# Long shared haplotypes identify the Southern Urals as a primary source for the 10th century Hungarians

**DOI:** 10.1101/2024.07.21.599526

**Authors:** Balázs Gyuris, Leonid Vyazov, Attila Türk, Pavel Flegontov, Bea Szeifert, Péter Langó, Balázs Gusztáv Mende, Veronika Csáky, Andrey A. Chizhevskiy, Ilgizar R. Gazimzyanov, Aleksandr A. Khokhlov, Aleksandr G. Kolonskikh, Natalia P. Matveeva, Rida R. Ruslanova, Marina P. Rykun, Ayrat Sitdikov, Elizaveta V. Volkova, Sergei G. Botalov, Dmitriy G. Bugrov, Ivan V. Grudochko, Oleksii Komar, Alexander A. Krasnoperov, Olga E. Poshekhonova, Irina Chikunova, Flarit Sungatov, Dmitrii A. Stashenkov, Sergei Zubov, Alexander S. Zelenkov, Harald Ringbauer, Olivia Cheronet, Ron Pinhasi, Ali Akbari, Nadin Rohland, Swapan Mallick, David Reich, Anna Szécsényi-Nagy

## Abstract

During the Hungarian Conquest in the 10th century CE, the early medieval Magyars, a group of mounted warriors from Eastern Europe, settled in the Carpathian Basin. They likely introduced the Hungarian language to this new settlement area, during an event documented by both written sources and archaeological evidence. Previous archaeogenetic research identified the newcomers as migrants from the Eurasian steppe. However, genome-wide ancient DNA from putative source populations has not been available to test alternative theories of their precise source. We generated genome-wide ancient DNA data for 131 individuals from candidate archaeological contexts in the Circum-Uralic region in present-day Russia. Our results tightly link the Magyars to people of the Early Medieval Karayakupovo archaeological horizon on both the European and Asian sides of the southern Urals. Our analyes show that ancestors of the people of the Karayakupovo archaeological horizon were established in the Southern Urals by the Iron Age and that their descendants persisted locally in the Volga-Kama region until at least the 14th century.

## Introduction

The Hungarians are the only Uralic-speaking ethnicity in Central Europe, with an early history that extends obscurely into the Early Medieval period, toward the east of the Carpathian Basin. Their history became richly documented beginning with the Hungarian Conquest period (895-1000 CE), which introduced striking innovations in burial rites and artifact assemblages to the Carpathian Basin. These cultural transformations are commonly interpreted as signatures of the arrival of a tribal alliance from the Eurasian Steppe, known as the early medieval Magyars (EMM)(*1–6*). Chronicles and oral tradition trace the origin of these Magyars to an eastern homeland(*1,2*) and a significant body of archaeological and linguistic research(*1,4,7–11*) points to the Cis-or Trans-Uralic regions as their likely homeland. Over the past century, the reconstruction of early Hungarian history has seen the emergence of diverse theories, as comprehensively reviewed by Zimonyi(*12*), all of which recognize the significance of the broader Volga-South Urals region in the ancestral formation process of the Magyars. However, the details of the migration speed and routes remain contentious. The Magyars likely encountered Turkic-speaking communities in both the Volga-Ural region and the North-Pontic steppe, based on material culture connections between these regions and the Carpathian Basin. The crossing of the Volga River by the Magyars in a westward direction has been estimated to have occurred between 460–830 CE(*1, 7, 13–17*), while their occupation areas in the northwestern Pontic region are inferred to have commenced between 670–860 CE(*7, 16–22*). Since these time ranges are broad, it is hard to date the beginning of this migration and its intermediate steps. Furthermore, it remains unclear where and how the language and community structure of the early Magyars was formed, as well as the roles the Circum-Uralic populations played in their ethnogenesis and confederation.

Based on parallels in material culture with the 10th-century Carpathian Basin, archaeologists have attributed some burial sites located around the South Urals to Magyars(*8*). We hereafter introduce the term ‘Karayakupovo Horizon’ (KH) to cover the diversity of the burial traditions and artefactual assemblages of the Southern Urals, including Cis- and Trans-Urals, dated to 750-1000 CE and associated with putative early medieval Magyars(*8,9*). East of the Urals, a reference cemetery of this horizon was excavated at Uyelgi, near Chelyabinsk(*23*). On the European side of the Urals, Bolshie Tigany in Tatarstan was a key site, and in the last decades, it was understood as a 9-10th century cemetery of Magyar groups that remained in the Volga-Urals(*3, 5, 8, 24–27*). People attributed to the Karyakupovo Horizont lived in a multilingual and ethnic context in the Circum-Uralic region, surrounded by Turkic, Finno-Permic, and Ugric-speaking people(*28*). Further evidence supporting the theory that Magyars settled in the Volga region during the Early Middle Ages are later reports of a Hungarian-speaking population in the Middle Volga and Lower Kama regions. This information comes from European travellers who visited an area known as *Magna Hungaria* in the 1230s (*29*), however, the survival of such communities has never been tested using ancient DNA data, which is the only direct way to verify population continuity and theories of ancestral origin.

Ancient DNA (aDNA) studies have generated large amounts of genetic data on ancient people of northern Eurasia which we co-analyze in this study along with our newly reported data(*30–68*). However, the Volga-Ural region from the Late Iron Age to Medieval times remained unstudied on the genome-wide level. *Csáky et al.* (2020)(*69*) and *Szeifert et al.* (2022)(*70*) provided insights into the connections between the 10th-11th century population of the Carpathian Basin and the Volga-Ural populations at the uniparental DNA level, while *Maróti et al.* (2022)(*65*) and *Gnecchi-Ruscone et al.* (2022)(*61*) generated genome-wide data for the Early Medieval Carpathian Basin itself. *Maróti et al.*(*65*) reported data from the 5th-10th centuries Carpathian Basin, showing that the Avars and Magyars represent distinct groups with East Eurasian genetic affinities. Based on their analyses, they argued that several sources were plausible for the immigrant 10th-century Magyars (named there as Conqueror Asia Core). This included modern Ugric-speaking Mansi proxy used in their canonical ancestry modeling, as well as groups descended from Huns/Xiongnu, and early and late Sarmatians. However, these sources do not align with prevailing linguistic and archaeological interpretations, so it is important to carry out tests with samples from the populations that are thought based on archaeological evidence to be the most plausible proximate ancestral sources.

The goals of the present study are twofold. First, we aimed to leverage the first genome-wide ancient DNA data from putative Volga-Ural source and adjacent populations of early medieval Magyars to understand their relationships to the new arrivals in the Carpathian Basin. Second, we attempted to model the deeper population history of those Volga-Uralic groups that showed especially strong connections to 10th-century Carpathian Basin Magyars and to document the extent of genetic continuity from the Iron Age to Medieval times in the Volga-Urals.

## Results

We used in-solution enrichment for more than 1.2 million single nucleotide polymorphisms (the “1240k” SNP capture panel(*71*)) to study the ancestry of 131 newly reported individuals from 31 archaeological sites in the Circum-Uralic area (see descriptions of relevant geography and sub-regions in the SI), dated from the Late Bronze Age (∼1900-1300 BCE) to the Late Medieval period staring ca. 1400 CE (see Figure 1, and Supplementary Text for detailed archeological descriptions of the newly sampled burials). In addition, we present data for six new individuals from the Carpathian Basin dated to the 10th century. For estimating genetic diversity and, in some cases, for modeling genetic origin, we grouped individuals by ecoregions/river basins and chronological periods(*72*); see Supplementary text, section II.A for details. For brevity, these periods are labeled by prevailing cultural groups in the region, e.g., *Russia_Belaya_Chiyalik* (Fig. 1), but cultural attribution did not play a role in the grouping process with one exception (the Karayakupovo Horizon).

**Fig. 1.**
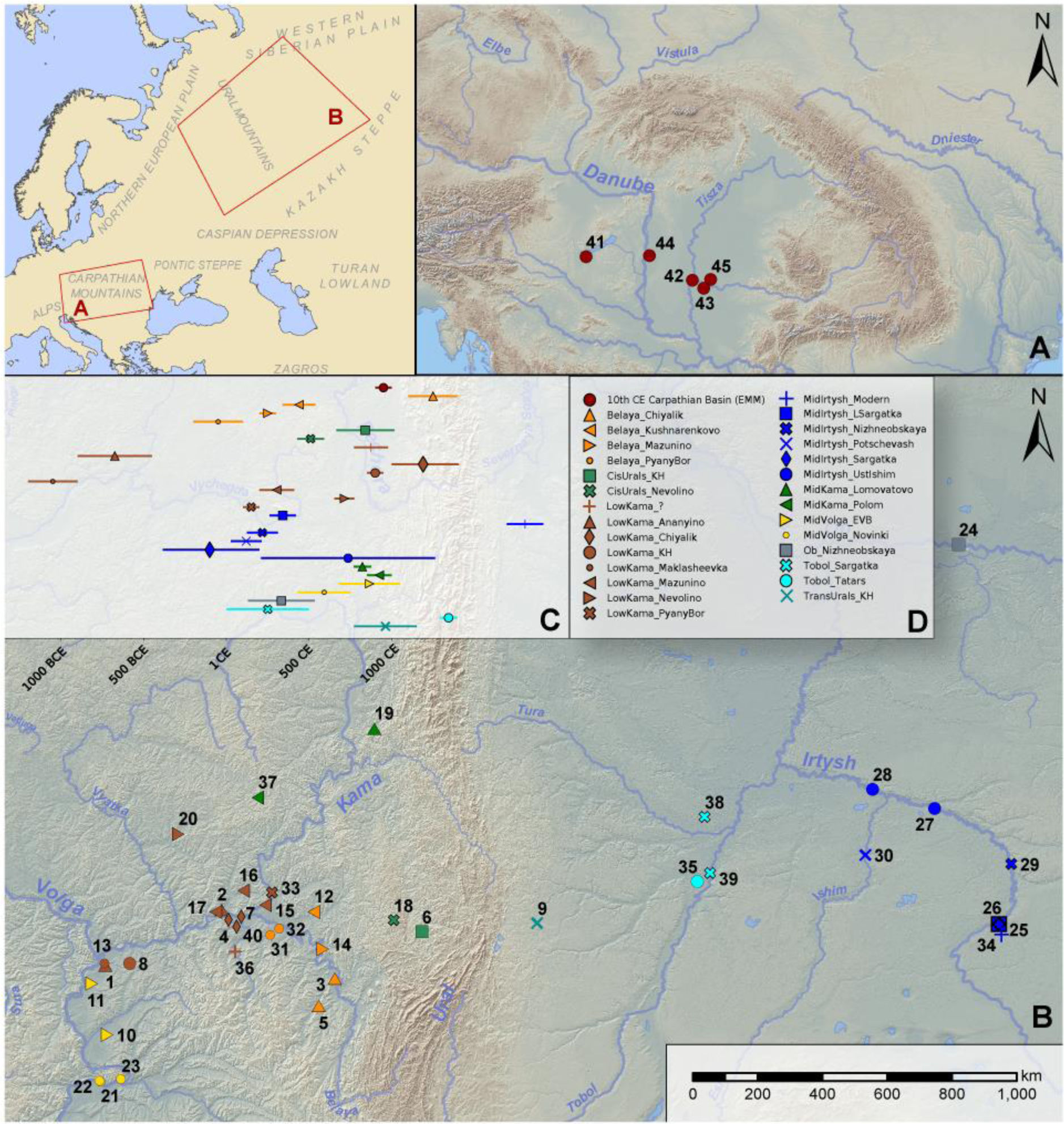
Locations and chronology of the studied burials. Archaeological sites in the Carpathian Basin (**A**) and in the Volga-Ural region (**B**) involved in this study, colored according to ecoregions: 1: Izmeri-7; 2: Rysovo-1; 3: Gornovo; 4: Gulyukovo; 5: Novo-Khozyatovo; 6: Karanayevo; 7: Zuyevy-Klyuchi; 8: Bolshie-Tigany; 9: Uyelgi; 10: Mullovka; 11: Tankeyevka; 12: Bustanaevo; 13: Devichiy-Gorodok-4; 14: Birsk-2; 15: Boyarsky-Aray; 16: Dubrovsky; 17: Turaevo-1; 18: Bartym; 19: Bayanovo; 20: Sukhoy-Log; 21: Brusyany; 22: Malaya-Ryazan’; 23: Novinki-1; 24: Barsov-Gorodok; 25: Borovyanka-17; 26: Borovyanka-18; 27: Ivanov-Mys-1; 28: Panovo; 29: Ust-Tarsk; 30: Vikulovo; 31: Kipchakovo; 32: Starokirgizovo; 33: Tarasovo; 34: Bogdanovo-2; 35: Putilovo; 36: Mellyatamak-3; 37: Varni; 38: Ipkul; 39: Starolybaevo-4; 40: Ust-Menzel’skoye; 41: Balatonújlak; 42: Szeged-Öthalom; 43: Kiszombor; 44: Harta-Freifelt; 45: Makó-Igási dülő Groups defined in this study are listed in panel **D** and their chronology is given in **C**.

Recent methodological developments have made it possible to detect long shared autosomal haplotypes between pairs of ancient genomes(*73,74*), often termed identical-by-descent (IBD) segments(*75*). Previously, this method was only applicable to high-quality genomic data for modern populations(*76, 77*). However, recent advancements allow its application to ancient individuals as well even if they have moderate fractions of their genome without high sequence coverage, leveraging the fact that human genetic variation is highly redundant so genotypes can be statistically imputed with high confidence from nearly incomplete genetic data(*74*). The IBD-sharing analysis is particularly useful for detecting distant relatives. We coupled this analysis with archaeogenetic methods relying on correlations of allele frequencies: PCA(*78*), *f*-statistics and derived methods(*31, 78–82*), as well as *ADMIXTURE*(*83*).

Our research protocol included several stages. First, we utilized PCA, supervised *ADMIXTURE* analysis, and network graphs visualizing individuals linked by shared IBD segments (see Methods for further details), to obtain a broad overview of the dataset. In the second stage, we focused on IBD connections between the Volga-Ural region and the population of the 10-11th century Carpathian Basin. In the third stage, we explored the genetic history of the Medieval Volga-Uralic groups using *f*-statistic methods(*31, 78, 81*), which allow formal tests of simple non-phylogenetic admixture models. To understand changes in population size and rates of close-kin marriages in this period, we explored runs of homozygosity (*hapROH*)(*84*).

### Genetic diversity in the Volga-Ural region

The Eurasian PCA in Fig. 2B reveals extraordinary genetic heterogeneity in the Early Medieval Volga-Ural region, with high variability in ancestry among individuals associated with certain regional and chronological groups. In the PC1/PC3 space (Fig. 2B), we observe an east-west genetic gradient from Northeast Asian (NEA) to Northwest Eurasian (NWE) genetic affinities. Most ecoregions of interest display high genetic diversity, with individuals from each region spread over large sections of the gradient (Fig. 2B). Notably, most of the newly sequenced 10th-century individuals from the Carpathian Basin are positioned along the NWE-NEA and NWE-Eastern Asian (EA) clines, with only two of them demonstrating a Central European genomic profile. We also conducted a supervised *ADMIXTURE* analysis (Fig. 2A), utilizing eight Neolithic and Early Bronze Age populations as proxy ancestry sources for the clustering algorithm. In our selection for the ancestral sources, we aimed to reflect the Neolithic/Bronze Age variation of North Eurasia (Fig. S1). Our findings reveal a widespread yet varying presence of Early Bronze Age Yamnaya-related ancestry across the region. This persistent Yamnaya-related ancestry(*30*), contrasted with the fluctuating levels of other ancestries, such as the Yakutia LNBA or Altai Neolithic(*68*), reflecting a patchwork of local genetic influences in the region.

**Fig. 2.**
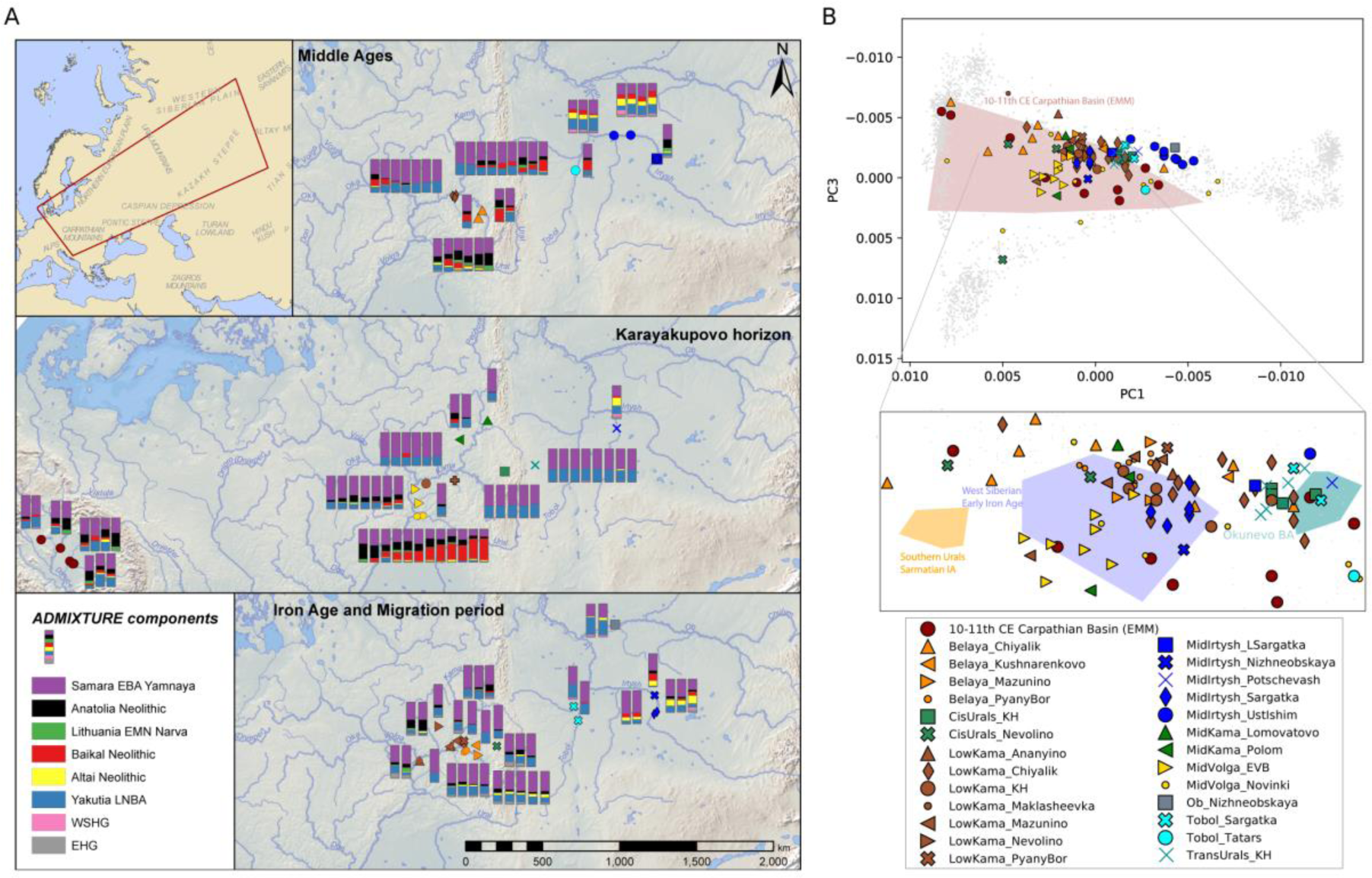
Principal component analysis and supervised *ADMIXTURE* analyses of the newly sequenced genomes. **A:** Supervised *ADMIXTURE* analysis (K=8) of the newly presented individuals, plotted on the map which shows their origin approximately. **B:** Eurasian-scale principal component analysis (PCA), with a projection of the newly sequenced individuals on modern genetic variation after Jeong et al. 2019(*50*). The PC1 and PC3 dimensions are depicted with the newly presented genomes and in polygonal representations with genomes from Early Medieval Magyars from the Carpathian Basin (*red*(65)), Early Iron Age Southern Urals ( *yellow*(49)), Iron Age Western-Siberia (*blue*(*56*)), and Bronze Age South Central Siberian (*green*(*38*)).

We applied genotype imputation(*73*), inferred IBD segments using the approach from(*74*), and constructed a network graph connecting individuals with shared IBD segments on a total of 1,333 individuals, comprising published data for 1,239 individuals from Asia and Europe and 94 individuals presented in this study (Fig. 3A). The graph’s edges were weighted based on the length of the most substantial IBD segment shared by two individuals (nodes). To de-noise the graph, we restricted the analysis to individuals connected by at least one 9 cM segment, were not separated in time by more than 600 years, and focused on the largest interconnected sub-graph. Details of the de-noising, visualization, and clustering approach are described in the Methods, for non-filtered network see Fig. S2. Twelve newly reported Iron Age individuals formed a cluster (with many previously published individuals) in the IBD network that we labeled *Eurasian steppe IA* in Fig. 3A (clusters were inferred with the Leiden community detection algorithm; we refer to them as “IBD-sharing communities’’ or simply “IBD clusters”). A total of 116 Early Medieval individuals from both the Volga-Ural region and Carpathian Basin formed another cluster (Fig. S3), labeled as *Urals-Carpathian EMA* in Fig. 3A. To discern and quantify the underlying differences among the identified network clusters, we analyzed network topology, similar to that described by Gnecchi-Ruscone et al. 2024(*85*), focusing on metrics such as degree of centrality (number of links held by a given node) or module strength measured based on summarized IBD-sharing between individuals (see Methods). The *Urals-Carpathian EMA* cluster’s average clustering coefficient reported by the Leiden algorithm was close to the mean of the other clusters. At the same time, its relatively high within-module (kw) and low between-module (kb) centrality exhibited distributions akin to the most cohesive clusters (Fig. S4 and Fig. S5). The *Urals-Carpathian EMA* cluster was loosely connected to the other IBD-sharing communities. Still, based on the low cluster coefficient, this separation could reflect gaps in sampling in time or space rather than true genetic isolation.

**Fig. 3.**
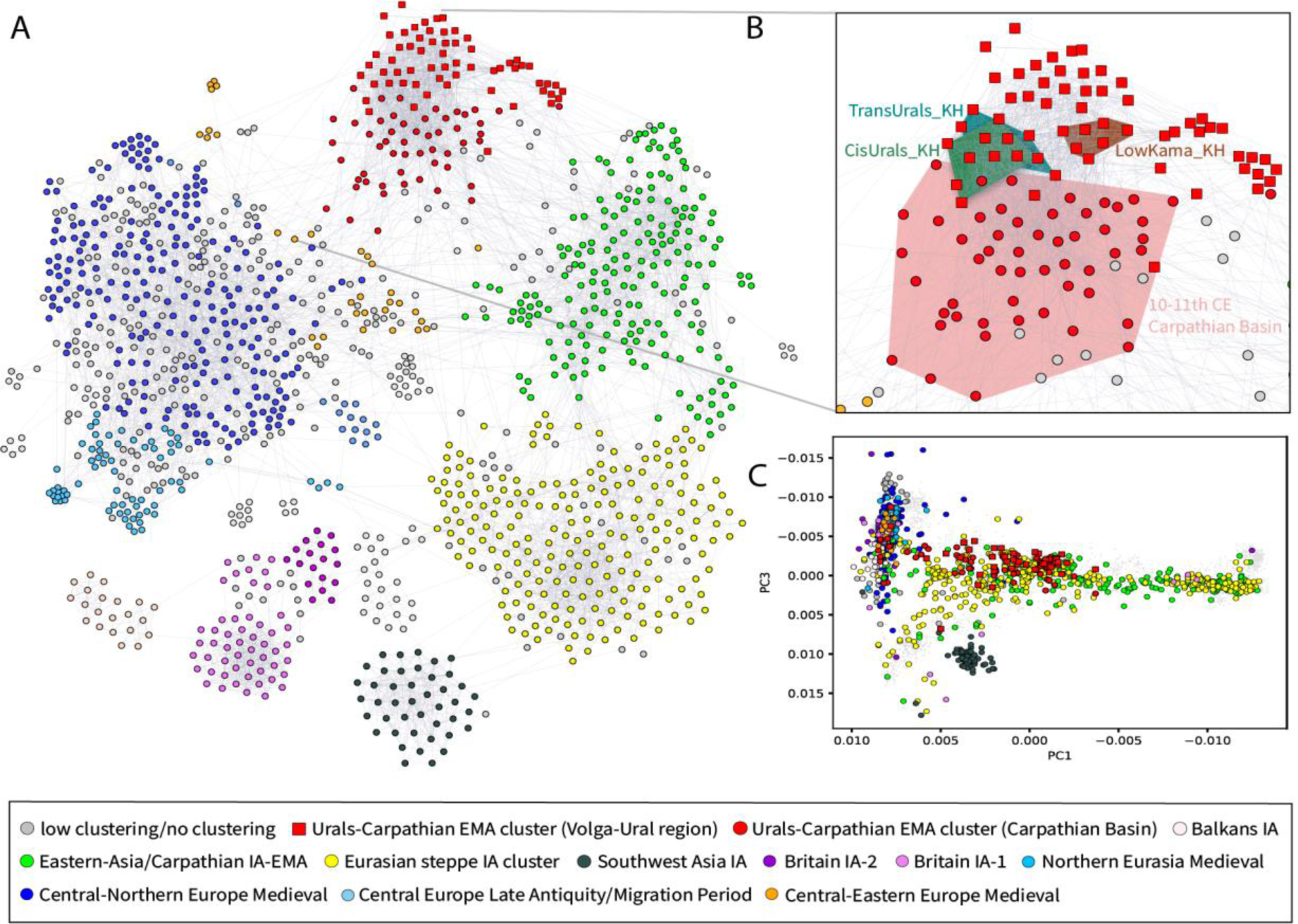
IBD network and visualization of the IBD clusters on PCA. **A:** A network graph of IBD sharing visualizing clusters of distant relatives for 1,333 ancient Eurasian individuals from the Iron Age to the Medieval Period (MultiGravity ForceAtlas 2, a force-directed layout algorithm(*86*) was used, and the Leiden algorithm(*87*) was used for clustering); **B:** Zoom in on the Urals-Carpathian EMA cluster within the network, highlighting the KH and 10-11th century Carpathian Basin individuals in the cluster; **C:** Individuals of the IBD-sharing network presented in PC1/PC3 spaces, projected on modern Eurasian individuals(*50*). The IBD clusters inferred with the Leiden algorithm are color-coded in all panels according to the legend in panel **A**.

Within the *Urals-Carpathian EMA* cluster, the published 10-11th century Carpathian Basin (CB) genomes(65) are grouped with our newly sequenced Volga-Ural Medieval samples. The Karayakupovo Horizon (KH) groups exhibited the highest degree of centrality (k) compared to other groups within the cluster (Fig. S6). In contrast, the early Medieval Carpathian Basin group exhibited a more diverse pattern. The strength (based on the summarized IBD-sharing) between and within the module links showed the high between-module connecting strength of the KH groups (Fig. S7). These findings highlight the ‘bridging’ role of the KH groups, linking the Volga-Uralic Medieval populations with the early Medieval Carpathian Basin individuals. However, some 10th-century Carpathian Basin individuals fall into the *East-Asia/Carpathian IA-EMA* cluster, reflecting a genetically diverse migration into the region. We have observed that PCA (and also the other allele-frequency-based methods) and the IBD network highlight different and complementary aspects of population structure: the former is more sensitive to East-West and North-South Eurasian genetic gradients, while the latter connects distant or close relatives who may have very different positions on these gradients (Fig. 3A, C; Fig S8).

### Early Medieval Magyars Fall within the Genetic Diversity of the Volga-Ural Region

We examined closely the genetic links between the Volga-Uralic groups and the 10th-century Carpathian Basin population forming the *Urals-Carpathian EMA* IBD cluster. The analysis showed that 10th-century Magyars in the Carpathian Basin exhibit significant genetic variation along PC1 (Fig. 2B), indicative of admixture during their migration westward or within the Carpathian Basin. As observed earlier, ancestries tracing back to the Baikal Neolithic and the Yakutia Late-Neolithic/Bronze Age varied across the EMM individuals. We mapped the proportions of these proxy ancestry sources onto our PCA (Fig. S9A). Consistent with the previously identified NWE-NEA and NWE-EA gradients, the EMMs demonstrate ancestry from two different East Eurasian sources. Specifically, those aligned with the NWE-NEA gradient exhibited a pronounced Yakutian Late-Neolithic/Bronze-Age ancestry, whereas those on the NWE-EA cline displayed higher levels of Baikal Neolithic ancestry. We note that these ancestry components do not reflect gene flows specifically from Yakutia or the Baikal region; rather, the proxy sources are reference groups for broad geographical regions and chronological periods. Males with distinct Y-chromosomal lineages from the Volga-Ural region showed a gradient along PC2 (Fig. S10) and the N1a∼ derived haplogroups seemed to be present at high frequency in the region in all periods explored (for mitochondrial DNA haplogroup frequencies, see Fig. S11). N1a-bearing EMM males were prevalent (Table S1), which also suggests their connection to the region. All of these results suggest that substantially different genetic sources on the Siberian genetic landscape could have contributed to the *Urals-Carpathian EMA* cluster of distant relatives in the 10th-century Carpathian Basin.

Next, we focused on specific cases of strong IBD links between early medieval Magyars and the population of the Volga-Ural region, providing case examples of long-distance migration within a few generations. We identified 28 pairs of individuals sharing more than two 12 cM or longer segments of their genomes (Table S1); of these, 11 pairs with the longest IBD segments are presented in Table 1 (for their ancestry proportions estimated with *ADMIXTURE* see Fig. S12). It is most likely that the degree of kinship for these pairs of individuals varied between the 6-8th degrees(*74*) (Fig. S13).

**Table 1:**
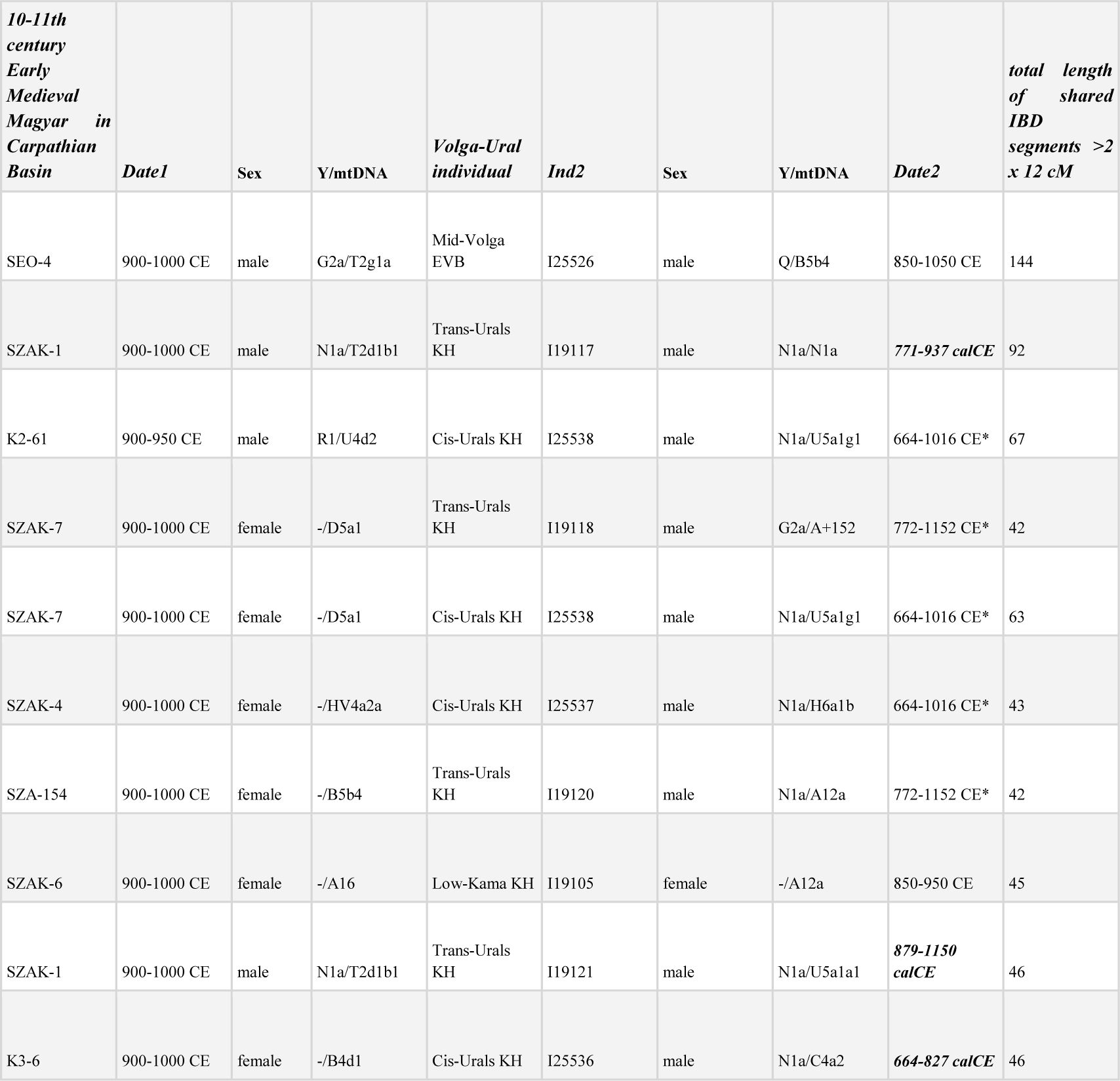
IBD connections between Medieval Volga-Uralic and Carpathian Basin individuals with ca. 6th to 8th degrees of kinship. Radiocarbon dates (calibrated, 95% confidence interval) are highlighted in bold. In other cases, the dating is based on the archaeological chronology of material culture. *summed probability densities, based on samples dated by radiocarbon data from the same site.

Archaeological and radiocarbon dating show that most IBD segments link individuals within a couple of hundred years of each other. Due to the wide ranges in radiocarbon dates, the connection between pairs of 6th-to 8th-degree relatives may stem from either a shared common ancestor or from ancestor-descendant relationships. The majority of the strong connections (>2 segments above 12 cM) of the EMM individuals are detected with the KH individuals (25 individuals) from various ecoregions. To better understand the connection between the two regions, we also conducted a qpWave analysis-based cladality test(*82*) (see Methods for details). This test evaluates whether the populations of interest (referred to as *left* populations) form a clade with respect to the *right* populations. We employed KH groups (Trans-Urals, Cis-Urals, and Low-Kama regions) and one joint group with European ancestry from the 10th to 11th centuries in the Carpathian Basin(*65*) (‘European cline’) as references. We used each group individually and tested whether they formed a clade with the *Urals-Carpathian EMA* cluster individuals from the Carpathian Basin. As *right* populations, we included early medieval contemporaneous groups spanning across the Volga-Ural region (Mid-Volga EVB, MidKama Lomovatovo, and Mid-Irtysh Potschevash), along with a group from Migration Period Buryatia, serving as a Central-Siberian reference point. Where a feasible model was lacking, we jointly tested with one KH and the European group. Our results showed feasible cladal structures for 17 individuals from the 10th to 11th century Carpathian Basin, and with the KH groups from the region (Table S2). We found that individuals sharing the highest levels of genomic segments shared identity by descent (IBD) with KH groups from both the Trans and Cis-Ural regions primarily showed feasible models with the Cis-Uralian KH group. Interestingly, Carpathian Basin individuals with lower levels of IBD sharing exhibited cladal structures linked to the Low-Kama KH group. Our cladality test provides a second and independent line of evidence, in addition to the IBD links for the connection between the two regions.

### Iron Age genetic continuity in the Medieval Volga-Ural region

To provide deeper insights into the genetic landscape of the Volga-Ural region, we applied *f4*-statistics, aiming to test if there was a significant genetic shift in this region since the Bronze Age. For this purpose, we compared allele sharing between the newly sequenced individuals and selected Bronze Age reference individuals from the Southern Urals (attributed to the Sintashta culture) and South-Central Siberia (attributed to the Okunevo culture, from the Minusinsk Basin), as shown in Fig. 4A. Our analysis revealed that during the late phase of the Early Iron Age, the level of the allele sharing was similar with both distant reference populations. However, as time progressed, an increasing number of individuals exhibited higher genetic affinity to one of these reference groups, suggesting that populations in the Circum-Uralic region experienced gene flows from nearby populations. These findings raise the hypothesis of shared ancestry for the Cis- and Trans-Ural individuals dated to the Early Iron Age (culturally from the Pyany Bor and Sargatka contexts), a conclusion further supported by our supervised *ADMIXTURE* analysis. A notable observation was the pronounced affinity of all of the Karayakupovo Horizon individuals to the South-Central Siberian BA reference group. This was also detectable in the case of the Low-Kama KH group. The significant allele sharing that prevailed in Low-Kama groups dates to the Medieval Period and is driven by individuals from the Chiyalik culture. These results highlight various population interactions during the Medieval period.

**Fig. 4.**
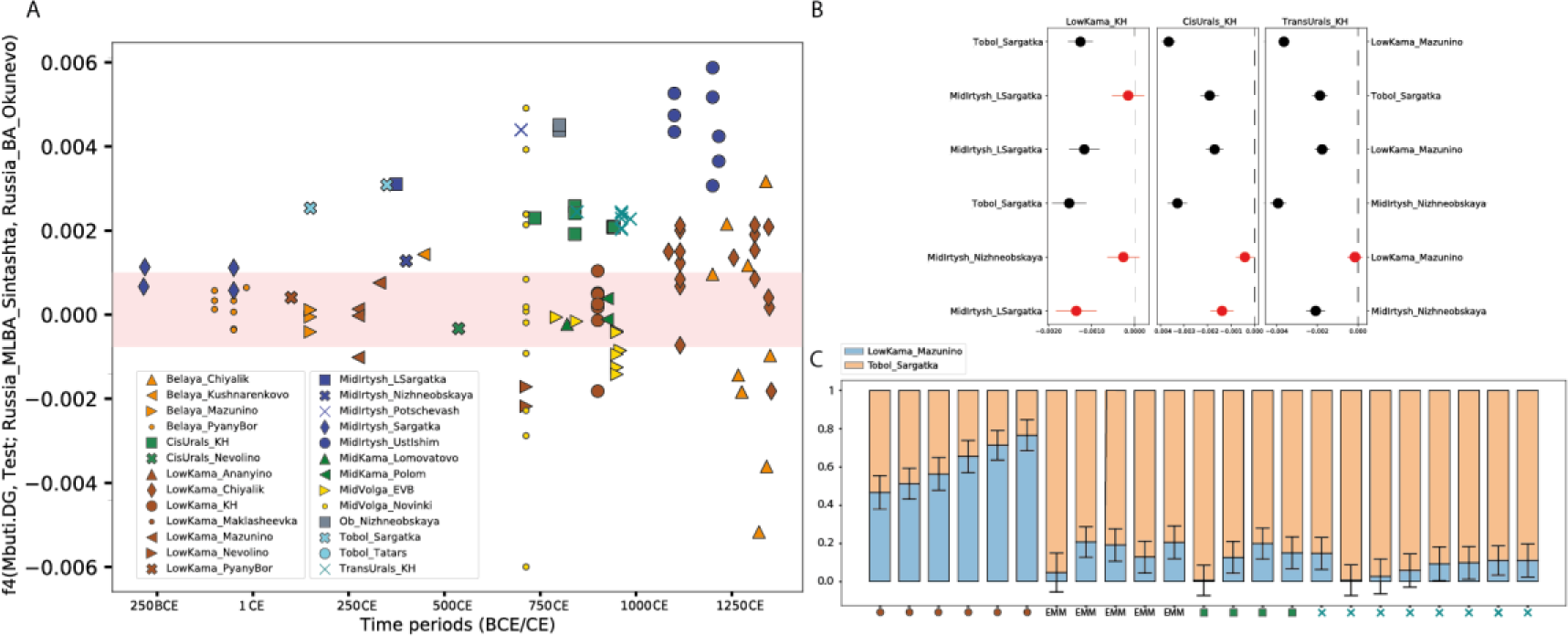
*f4*-statistics and admixture models illustrating allele-sharing and genetic affinities among newly sequenced individuals and Bronze/Iron Age reference groups. **A:** *F_4_*-statistics for the newly sequenced individuals, excluding those from the Maklasheevka and Ananyino cultural contexts. The Y-axis represents the allele-sharing values with two Bronze Age reference groups (red band indicates |Z-score| < 3). The X-axis shows the timeline. **B:** f_4_-statistics comparing allele sharing between KH groups and Migration Period Volga-Uralian reference groups (Mbuti.DG, KH_test_group; MigrationPeriod_reference_group1, MigrationPeriod_reference_group2). Markers indicate affinities with left and right reference groups. Red markers denote |Z-score| < 3. **C:** A two-way admixture model (*qpAdm*) for the Karayakupovo Horizon and early medieval Magyar individuals (with feasible *qpAdm* models) from the 10-11th century Carpathian Basin (from Table 1) that exhibited strong IBD sharing (>42 cM in IBD segments longer than 12 cM; see Table S1 for additional details). For additional EMMs modeled with this two-way *qpAdm* setup see Supplementary Dataset 6.

To test the Iron Age/Migration Period (for a detailed description of the archeological chronology in the region, see SI I.A) individuals for evidence of continuity with early Medieval KH individuals, we used two complementary f4-statistics. Initially, we tested allele sharing between our focal (KH), and both EIA Southern Uralic (associated with Sarmatian culture context) and Western Siberian groups (Sargatka horizon), which revealed reduced allele sharing with the former group (Fig S14). Furthermore, allele-sharing analyses among Western Siberian groups revealed significant affinity between the Cis and Trans-Urals KH groups and EIA groups in the Irtysh River region. In the second stage, we analyzed early Migration Period reference populations from the wider Volga-Ural region and allele sharing among KH groups (Fig 4B). This included the Low-Kama Mazunino group and groups from the Tobol and Mid-Irtysh regions from the late Sargatka horizon and the Nizhneobskaya culture. The latter is distinct both archaeologically and genetically from the local continuum. Compared to the other references, we observed significant allele sharing with the Mid-Irtysh and Tobol groups from the late Sargatka horizon. These findings indicate genetic continuity in the KH groups from the Early Iron Age, rooting their ancestry in the Irtysh and Tobol River regions.

To model possible admixture scenarios and quantify the proportion of the Migration Period ancestral sources (for KHs and EMMs with direct connections to KH individuals [Table 1]) we employed *qpAdm* analysis (Fig. 4C) (for the detailed settings, see Material and Methods). We purposely avoided rotating modeling approaches exploring large sets of alternative proxy sources(*88*). Instead, we utilized a two-way modeling strategy with proxy sources on both sides of the Urals in the Migration Period: the Sargatka cultural group in the Irtysh/Tobol basins, and Mazunino in the Low Kama basin. Their archeological importance in the late phase of the Iron Age in the Ural region and also their separation in the spaces of *f4*-statistics and outgroup *f3-* statistics (Fig. S15 and S16) justified the use of these sources for *qpAdm* analysis. Archaeological context also supports the significance of these groups as they potentially influenced the Kushnarenkovo and later Karayakupovo archeological cultures(*8*). In the case of the Mazunino group, we used the Low-Kama sub-group, which has sufficient coverage in our data. Out of the 26 analyzed individuals, the two-way model was a fit (p-value > 0.05) in 22 cases (for the list of outgroups see Material and Methods). The Tobol Late Sargatka ancestry was notably prevalent among the Trans-Ural KH, Cis-Ural KH, and early Medieval Magyar individuals, at least ∼70% (for detailed results, see Supplementary Dataset 6). While all EMM and KH groups likely share the same Trans-Uralic ancestry, some (Low-Kama KH, see Fig. 4C) mixed extensively with local groups to the west of the Urals.

A time-ordered IBD graph in Fig. S18 illustrates biological continuity, especially between the Early Medieval KH groups and those from the Late Medieval Chiyalik cultural contexts in the Belaya and especially Low-Kama regions. The similarity in *ADMIXTURE* profiles (Fig. 2A) further supports the continuity of the KH-type ancestry into the later Medieval period. In contrast, the Belaya region in the Late Medieval period is more diverse genetically, with several individuals having European and East Asian genetic profiles (supported by IBD connections outside the *Urals-Carpathian EMA* cluster).

To explore the demographic history of the Volga-Ural groups from a different perspective, we utilized the *hapROH* method to identify long runs of homozygosity (ROH), as shown in Figs. S18 and S19(*84*). This analysis revealed that KH individuals probably had a low effective population size (*Ne*), evidenced by the ROH segments in their genome (Fig. S19). The number of ROH segments per group correlated negatively with other estimates of genetic diversity used in this study. Our *Ne* analysis further indicated that both Early Medieval Low-Kama KH and Late Medieval Low-Kama Chiyalik groups had consistently smaller population sizes than neighboring groups across different periods.

## Discussion

In this study, we report genome-wide data for 131 ancient human genomes from 1900 BCE to 1400 CE in the Circum-Ural region and the Carpathian Basin. The genetic gradients displayed on the PCA by the Volga-Ural region groups (Fig. 2B) align with the modern genetic variation found in Eurasia’s forest and forest-steppe zones (the northern one) and the steppe zone (the southern one), respectively(*68*). The Asian end of the northern gradient is linked to the Yakutian LNBA population, which is a genetic „tracer dye” for Uralic speakers in North Eurasia(*68*). The analysis of identity-by-descent (IBD) chromosome segments revealed distant relatedness between Early Medieval Circum-Uralic individuals from the Karayakupovo Horizon sites and the EMM 10th-11th centuries population from the Carpathian Basin. We termed the IBD cluster of distant relatives as “*Urals-Carpathian EMA*” (Fig S5), which showed a genetic gradient stretching from Europe to Northeast Asia on PCA, and distinct from the *Eurasian steppe Iron Age* and *East Asia/Carpathian IA-EMA* clusters (Fig. 3B-C, Fig S3).

Our findings demonstrate that Cis- and Trans-Uralic Karayakupovo Horizon sites are linked to 10th-11th-century Carpathian Basin individuals via IBD. These IBD connections are supported by similarity in *ADMIXTURE* profiles and *qpWave* based cladality tests. Notably, individuals from the Hungarian Szakony-Kavicsbánya site displayed the highest similarities to the Volga-Uralic population in *ADMIXTURE* clustering and IBD sharing. Archaeological artifacts from this site and burial customs show direct parallels in Uralic cultural contexts(*89*). These combined findings provide the first compelling genetic evidence for a Uralic origin for an important part of the ancestry of 10th-century Magyars in the Carpathian Basin. EMMs from the Carpathian Basin mostly demonstrate Yakutian LNBA-type ancestry associated with the northern (forest and forest-steppe) Eurasian gradient. Still, some also demonstrate Baikal Neolithic-related ancestry associated with the southern (steppe) Eurasian gradient (Fig. S7).

These results imply that they (or their ancestors) have at least two genetic sources outside the Carpathian Basin, and we confirmed the Circum-Uralic one. Considering the archeological, historical, and genetic results, our findings are consistent with a scenario in which the initial area of the EMM migration to the Carpathian Basin was located in the Volga and Ural regions, where traces of admixture are not observable with Central/East-Eurasian ancestry bearing groups (such as people usually attributed to the Turkic speakers(*38, 50*). The results presented in our paper align with the Uralic (Ugric) basis of the Hungarian language, which has its first written documents only as late as 11th century Hungary(*90*). Among the possible Early Medieval influxes to the Carpathian Basin, the Hungarian language was most probably brought from the Southern Ural region (by descendants of the members of the Karayakupovo archaeological horizon), among others by those Magyars who shared the *Urals-Carpathian EMA cluster*. However, it is important to emphasize that the Magyars as steppe nomadic societies, had diverse cultural backgrounds and functioned as multiethnic/multilingual communities(*16, 91, 92*). The most recent reconstructions of the Magyar migration based on material culture evidence date the subsequent population movement from the Volga to the Pontic Steppe as late as the early 9th century CE, and from there to the Carpathian Basin by the end of that century(*3, 6*). The tight connectedness of the Urals-Carpathian EMA cluster and the genetic characteristics of a part of the EMM indicate a rapid migration from the Volga-Ural to the Carpathian Basin and a rather short stop in the North-Pontic area. This later area could have been the site for the integration and alliance with further Turkic-speaking tribes(*3*).

The first emergence of Karayakupovo-type genetic ancestry west of the Urals was detected by 550 CE. This ancestry did not extend as far west as the Volga-Kama confluence or the Volga’s west bank by the Samara Bend, as it is absent in the group with Novinki-type burial practices. Furthermore, our findings indicate that individuals from the Early Volga Bulghar Mullovka and Tankeevka cemeteries either show no genetic links to the Karayakupovo Horizon sites. Our analyses indicate little or no IBD connection between the EMMs and proto-Ob-Ugric groups in Western Siberia, despite their close geographical proximity for 1500–2000 years after their split estimated by linguistic models and chronology(*70*).

As the KH groups demonstrated notably strong IBD connectivity despite considerable geographical distances (Low-Kama, Cis-Urals, Trans-Urals), we investigated the extent of their shared population history. Using multiple f4-statistics, we demonstrated that the KH groups shared the most alleles with groups from the Irtysh and Tobol regions throughout the Iron Age and Migration Period. This evidence supports the hypothesis of a Trans-Uralian origin for the late Karayakupovo-type ancestry. Our proximal *qpAdm* analysis showed that the Low-Kama KH group could be modeled as a combination of Pyany-Bor/Mazunino and Tobol Late Sargatka-related ancestries, resulting in a distinct local KH variant. In contrast, the other KH groups have much lower Pyany-Bor/Mazunino ancestry. We demonstrate that the proxy ancestry sources we used in our *qpAdm* analyses (Pyany-Bor/Mazunino to the west of the Urals and Tobol Late Sargatka to the east of the Urals) are much closer to the actual sources than those used in the Maróti et al. *qpAdm* approach, which suggested using modern Mansis, early/late Sarmatians, and Xiongnu as proxies for modeling the ancestors of the EMMs. Based on the connections with the KH individuals, we show that an important stratum of the EMMs (named by Maróti et al. as ‘Conqueror Asia Core’) can be traced to the Early Medieval Circum-Uralic region. Also, with *qpAdm* modeling, we detected local biological continuity from the Iron Age to the Early Medieval times in these regions. However, we avoided extensive *qpAdm* screenings across multiple ancestry sources closely timed to target groups, similar to the approach used in Maróti et al. 2022(*65*), due to the high risk of false-discovery rates (FDR), as demonstrated by Yüncü et al. 2023(88). Additionally, archaeologists have determined that in the early Middle Ages, the area east of the Ural Mountains, extending to the Ob River in the present-day Omsk region, had an extremely low population density. The total number of excavated graves from the 6th to the 10th centuries AD does not exceed 300(*93*). We have detected extended genetic signals indicating small population sizes both east of the Urals and in the Cis-Urals KH group. These findings provide significant evidence of sparse and low population sizes in these regions during this period.

The Late Medieval Low-Kama Chiyalik group shows strong continuity within the *Ural-Carpathian EMA IBD* cluster. This is indicated by a high level of connectivity within the IBD-sharing community and limited IBD sharing beyond it. Moreover, they are similar to the KH groups on an allele frequency level. In contrast, individuals linked with the Chiyalik culture from the Belaya Region are more diverse genetically and fall outside the *Urals-Carpathian EMA* cluster. These findings suggest the potential influx of newcomers during the Golden Horde period, who likely introduced different East Eurasian genetic ancestries. Considering the late 14th-century radiocarbon dates for the Chiyalik individuals, it is reasonable to assume the presence of remaining Magyars, archaeologically represented by a local variety of the Chiyalik culture, mainly in the Lower Kama River Valley(*94, 95*). By analyzing the effective population size, we estimate that the Low-Kama Chiyalik group comprised at least a few thousand individuals during the Late Medieval times. These results suggest that descendants of the *Uralic-Carpathian EMA* IBD-sharing community survived in Late Medieval times in considerable numbers in the Kama region. We assume that the Low Kama region near the Belaya-Kama confluence was the area that was called Magna Hungaria by Friar Julian in the 13th century(*29*). In addition to this historically documented data, the regional toponymy suggests the presence of Hungarian-speaking groups there until the 16th century, when, after the collapse of the Golden Horde imperial space, they were absorbed into the Late Medieval populations of modern-day Bashkortostan, Tatarstan, and Udmurtia(*6, 96, 97*).

## Material and methods

### Sampling and sample selection

Based on years of collaborations with local archaeological experts governed by bilateral collaboration agreements, we selected the most relevant samples, which were verified by radiocarbon dating. We aimed to collect graves for this research with grave materials characteristic of local cultures. In the **Trans-Urals,** our sampling involves individuals buried in the Sargatka cultural context from the Middle Irtysh (300 BCE - 200 CE) and the Tobol (100-350 CE) river basins. The later Trans-Uralic population groups are represented by burials attributed to the Nizhneobskaya, Potchevash, and Ust’-Ishim cultures, and the Uyelgi cemetery attributed to the Karayakupovo Horizon. In the **Cis-Urals**, we undertook a dense sampling from sites attributed to the Maklasheevka Late Bronze Age (1100-900 BCE), Post-Maklasheevka Ananyino Early Iron Age (900-250 BCE), Pyany Bor Early Iron Age (250 BCE-150 CE), Mazunino (150-450 CE), and Nevolino (400-850 CE) archaeological entities. The Migration-period population archaeologically related to the Trans-Uralic groups is represented by one individual from the Kushnarenkovo cultural context (550-700 CE). The sampling of the Medieval individuals of the Volga-Ural region involves the peripheral regions of the Volga Bulgaria, to the east of main cities and densely populated areas. The cultural context of our Medieval samples can be mainly described as the “Muslim burials with pagan elements in burial rites”, and it is usually attributed to the Chiyalik culture. The sites of the **Karayakupovo Horizon** to the west of the Urals are represented by Bolshiye Tigany from the Lower Kama region (800-900 CE). We also included some sites contemporaneous with the Karayakupovo Horizon, but archaeologically attributed to other groups: the Novinki-type sites (700-850 CE) and the Tankeevka cemetery (850-1000 CE), a local group of the Khazar-Khaganate nomads and the Early Volga Bulghars (EVB) respectively (see further details in the SI). Two individuals from the Polom cultural context and one from Lomovatovo represent the Mid-Kama population groups that are contemporaneous with the people of the Karayakupovo Horizon sites.

### Ancient DNA data generation

117 samples were cleaned and powdered in the Budapest Laboratory of Archaeogenetics (Institute of Archaeology RCH) as described in Szeifert et al. 2022 and shipped to Harvard Medical School. Three samples were prepared in Vienna, and seven samples were prepared in Ostrava and shipped to the Harvard laboratory. In dedicated clean rooms, we extracted DNA manually with spin columns(*98, 99*) or automated using silica magnetic beads and Qiagen PB buffer on the Agilent Bravo NGS workstation(*100*) and converted it into barcoded double-stranded partial Uracil-treated libraries(*101*), which we enriched in solution for sequences overlapping 1.24 million SNPs [1240k: Fu et al. 2013(*33*), Twist: Rohland et al. 2022(*102*)] as well as the mitochondrial genome. For each library, we sequenced approximately 30 million reads pairs (median of 29.747M reads) of the enriched library using Illumina instruments [NextSeq500, HiSeq X]; we also sequenced several hundred thousand sequences of the unenriched library.

### Bioinformatic analysis

Samples were sequenced to generate raw paired-end reads; these were prepared for analysis by performing the following steps: preprocessing/alignment, and post-alignment filtering to enable variant calling. Raw reads were demultiplexed by using identifying barcodes and indices to assign each read to a particular sample, prior to stripping these identifying tags. Paired-end reads were merged into a single molecule using the base overlaps as a guide, Single-ended reads were aligned to the hg19 human reference genome (https://www.internationalgenome.org/category/grch37/) and the basal Reconstructed Sapiens Reference Sequence (RSRS)(*103*) mitochondrial genome using the samse aligner of bwa(*104*). Duplicate molecules were marked based on barcoding bin, start/stop positions and orientation. The compuational pipelines with specific parameters are publicly available on github at: https://github.com/dReichLab/ADNA-Tools and https://github.com/dReichLab/adna-workflow. For calling variants, a pseudo-haploid approach is used at targetted SNPs, where a single base is randomly selected from a pool of possible bases at that position filtering by a minimum mapping quality of 10 and base quality 20, after trimming reads by 2 base pairs at both 5’ and 3’ ends to remove damage artifacts.

### Principal component analysis(PCA)

PCA analysis was carried out with EIGENSOFT software(*105*) (version 5.0) with lsqproject: YES and shrink mode: YES settings. For projection, we used modern-day Eurasians from the Affymetrix Human Origin array and after merging our dataset with the array we restricted our analysis to 597573 SNPs.

### ADMIXTURE analysis

Before running ADMIXTURE(*83*) we pruned our dataset with plink (version 3)(*106*). We have used the-geno 0.95 option to ensure that we included sites where most individuals were covered at least once. After that we used –indep-pairwise 200 24 0.4 parameters for linkage disequilibrium (LD) pruning. We also removed individuals who were closely related (up to 3rd degree). We performed supervised ADMIXTURE clustering with K=8. We used Neolithic/Early Bronze Age populations as sources to reflect the overall distribution of different ancestries through Eurasia. We tried to involve well-represented groups (> 4 individuals) with high-coverage data. We intentionally aimed to reconstruct a similar ADMIXTURE reference set presented in Zeng et. al 2023(*68*). We have found this set useful in understanding the pre-historical genetic composition of our newly published individuals.

### Genotype imputation

For imputation, we applied the GLIMPSE (v.1.1.1)(*73*) software with the 1000 Genome Project as the reference panel on VCF files to estimate genotype posterior at bi-allelic SNP sites. For IBD analysis, we restricted to SNPs in the 1240k capture, which are informative for ancient DNA studies. These VCF files were generated using bcftools mpileup (v1.10.2)(*107*) applied on sequence data in aligned BAM format. A full description of the imputation pipeline is provided in Supplementary Note 3 and Figure 1b of Ringbauer et al., 2024(*74*).

### IBD-sharing analysis

We utilized the method described in Ringbauer et al.(*74*) to detect identity-by-descent segments. In the downstream analysis, we included samples that had sufficient coverage on the 1240k SNP positions and that matched our research criteria, focusing on geographical location (North Eurasia) and timeframe (∼1000 BCE to modern times).

### IBD-sharing network

All the IBD networks were built with Gephi (v.0.10.1) software(*108*). The graph’s edges were weighted based on the length of the most substantial shared IBD segment between two individuals, referred to as nodes. We removed IBD segments below a threshold of 9 cM and connections that spanned over 600 years for clarity and precision. Additionally, we maintained nodes connected by at least two edges and focused on the largest interconnected segment of the graph. Visualization was achieved using the MultiGravity ForceAtlas 2, a force-directed layout algorithm(*86*). In the processed graph, clusters were discerned using the Leiden algorithm(*87*), maintaining algorithmic independence. For further analysis of the clusters defined by the Leiden algorithm, we explored several key metrics: degree centrality (k), which measures the number of connections a node has; within-module centrality (k_w_), quantifying the connections within each defined cluster; and between-module centrality (k_b_), which assess the connections between different clusters. To calculate the strength (based on the summarized IBD-sharing) of within and between module links, we utilized the Python NetworkX package(*109*), considering our predefined groups as modules.

### f-statistics

We computed *f3* and *f4*-statistics with the ADMIXTOOLS software package(*78*) with the qp3pop (allsnsp:YES) and qpDstat (f4Mode: YES; printsd: YES) packages. For the *f3*-statistics we used an outgroup approach as follows (Test1, Test2; Mbuti). For *f4*-statistics we used (Mbuti, Target; Test1, Test2) to check the genetic affinities between two possible ancestral populations. For the pairwise cladality test, we used the ‘*qpWave pairs*’ test from the R software package *Admixtools 2* with default settings(*82*). We designated 10th to 11th-century Carpathian Basin individuals as *targets* and the KH groups as *left* populations. The right populations included Mid-Volga EVB, Mid-Kama Lomovatovo, Mid-Irtysh Potschevash, and the Buryatia Xiongnu group(*54*). For models that were unfeasible, we incorporated Early Medieval individuals with no Eastern Eurasian ancestry as *left* populations (Maróti et al., 2022)(65). We chose the one with the highest p-value when multiple feasible models were available.

### QpAdm analysis

For the qpAdm analysis, we used the *Admixtools 2* R package(82), with the following carefully selected(81) outgroups: Mbuti.DG, Ami.DG, Italy_North_Villabruna_HG, Turkey_N.SG, Russia_Ekven_IA.SG, Russia_DevilsCave_N.SG, Russia_Sidelkino_HG.SG, Russia_Caucasus_Eneolithic, Tarim_EMBA1. We avoided using the rotating approach as in complex demographic histories the direction of the geneflows cannot be defined accurately(88).

### Consanguinity test (ROHs)

Detecting runs of homozygous blocks with hapROH(84) software can provide signals of consanguinity, whereas small homozygous runs are indicative of a small recent effective population size. The program was used with default parameters for pseudo-haploid genotypes with at least 400k SNP covered. The *Ne* module of this program was also used to estimate effective population sizes with CI, considering 4-20cM ROHs.

### Radiocarbon dating

Radiocarbon dating of 10 DNA samples was performed in the Penn State’s Radiocarbon Laboratory (PSUAMS codes). The BP values were calibrated in the Oxcal program 4.4 with a calibration curve IntCal 20 (*110, 111*)

## Acknowledgements

We thank N. Adamski, V. Bódis, R. Bernardos, N. Broomandkhoshbacht, K. Callan, E. Curtis, M. Ferry, I. Greenslade, L. Iliev, A. Kearns, M. Michel, L. Qiu, K. Stewardson, N. Workman, F. Zalzala, and Z. Zhang for their work in sample management, processing, and laboratory work; A. Bogachev, E. Chernykh, O. Flegontova, R. Goldina, E. Kazakov, E. Kitov, A. Kochkina, and A. Tishkin for providing and collecting archaeological material; I. Lazaridis, M. Mah, A. Micco, and I. Olalde for their bioinformatic work; E. Szász for the visualization; and D. Gerber for essential feedbacks.

## Funding

US National Institutes of Health grant HG012287 (Ancient DNA research in Boston)

Allen Discovery Center program, a Paul G. Allen Frontiers Group advised program of the Paul

G. Allen Family Foundation (Ancient DNA research in Boston)

John Templeton Foundation grant 61220 (Ancient DNA research in Boston, L.V., P.F.)

Private gift (Ancient DNA research in Boston, L.V., P.F.)

Howard Hughes Medical Institute (HHMI)

Priority Research Theme proposal of the Eötvös Loránd Research Network (2019-2023 ELKH, 2023-HUN-REN), in the frame of the “Archaeogenomics research of the Etelköz region” project (A.T., A.Sz-N., B.G.M., B.Sz)

PPKE-BTK-KUT-23-3 project, funded by the Faculty of Humanities and Social Sciences of Pázmány Péter Catholic University (A.T.)

Czech Ministry of Education, Youth and Sports (program ERC CZ, project no. LL2103) (L.V., P.F.)

Czech Science Foundation (project no. 21-27624S) (P.F.)

Private support from Jean-Francois Clin (L.V., P.F.)

Russian Science Foundation grant no. 24-18-20055 (S. B.)

Russian Science Foundation grant no. 24-28-20283 (I. G.)

Russian Science Foundation grant no. 23-78-10057 (R. R.)

Research of O.P. and I.C. was supported by state assignment #FWRZ-2021-0006

## Author contribution

Designed the study: B.Gy., L.V., A.T., P.F., D.R., A.Sz.-N.

Collected/provided archaeological material: L.V., A.T., P.L., D.S., A.S., N.M., A.Z., S.B., I.G., M.G.B., I.C., R.P., O.C., O.P., R.R., E.V., M.R., A.Ko., A.C., A.Kh., I.G., S.Z., F.S.

Laboratory analysis: B.Sz., N.R.

Performed bioinformatics processing of the data: H.R., A.A., S.M. Performed analysis: B.Gy., L.V., A.Sz.-N.

Wrote the paper: B.Gy., L.V., P.F., B.Sz., V.Cs.

Wrote archaeological supplement: L.V., D.S., A.Z., S.B., I.G., O.K., D.B., A.Kr., O.P. Supervised the manuscript: A.T., D.R., A.Sz.-N.

