## Supplementary Material for "Long shared haplotypes identify the Southern Urals as a primary source for the 10th century Hungarians"

#### Supplementary Materials

##### Supplementary Text

I. Archaeological context 3

### I. Archaeological context

#### I.A. Chronological terms and definitions

The chronology of the population dynamics in the Circum-Uralic area over the first millennium BCE - early second millennium CE (~2900-650 yBP) is based on the archaeological data and reflects the main stages of the development of the material culture. Despite strong criticism from the last 60 years of archaeological discussion, the term “archaeological culture” still plays an important role as the main research category in the region of study. During the last 30 years, it lost its previously dominating "ethnical" sense and was partially pushed out from literature by using more diverse definitions such as "archaeological horizon" (a combination of similar features that can be observed at the same time in different cultures) and "cultural type" (an assemblage of cultural features which may not form a clearly defined area of sites, may be represented in a very limited area or even by single site). We summarized in the following sequence the spatial-temporal variations of those archaeological entities that are used for the general periodization of regional history.

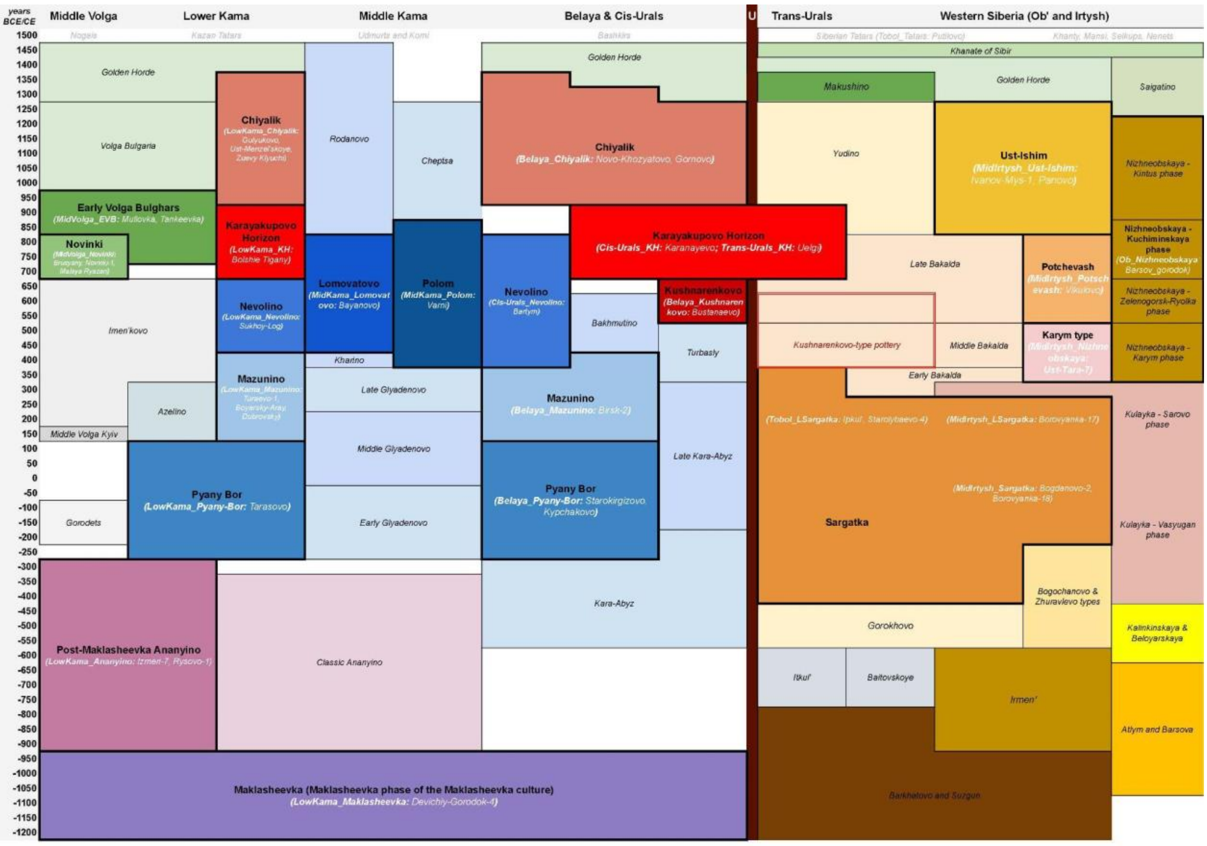

***Fig. SI.A.1.*** *Main archaeological entities of the Circum-Uralic region from the Early Iron Age to the Middle Ages.*

The chronological boundaries slightly differ in different areas of the region and the cultural transformations could last for several decades.

#### I.A.1. Early Iron Age, phase 1 (850 – 250/200 BCE)

The chronological frame of the earliest phase of the Early Iron Age in the Cis-Uralic part of the region is bounded by the cultures of the Ananyino area (Kuz'minykh, 2006; Kuz'minykh, Chizhevskiy, 2021), shaped by long-term interaction with the Scythian-period nomads of the steppe region. The upper chronological boundary of the period coincides with the Sarmatian migration and the establishment of the Sarmatian cultures (Prokhorovka) (Chizhevsky 2021).

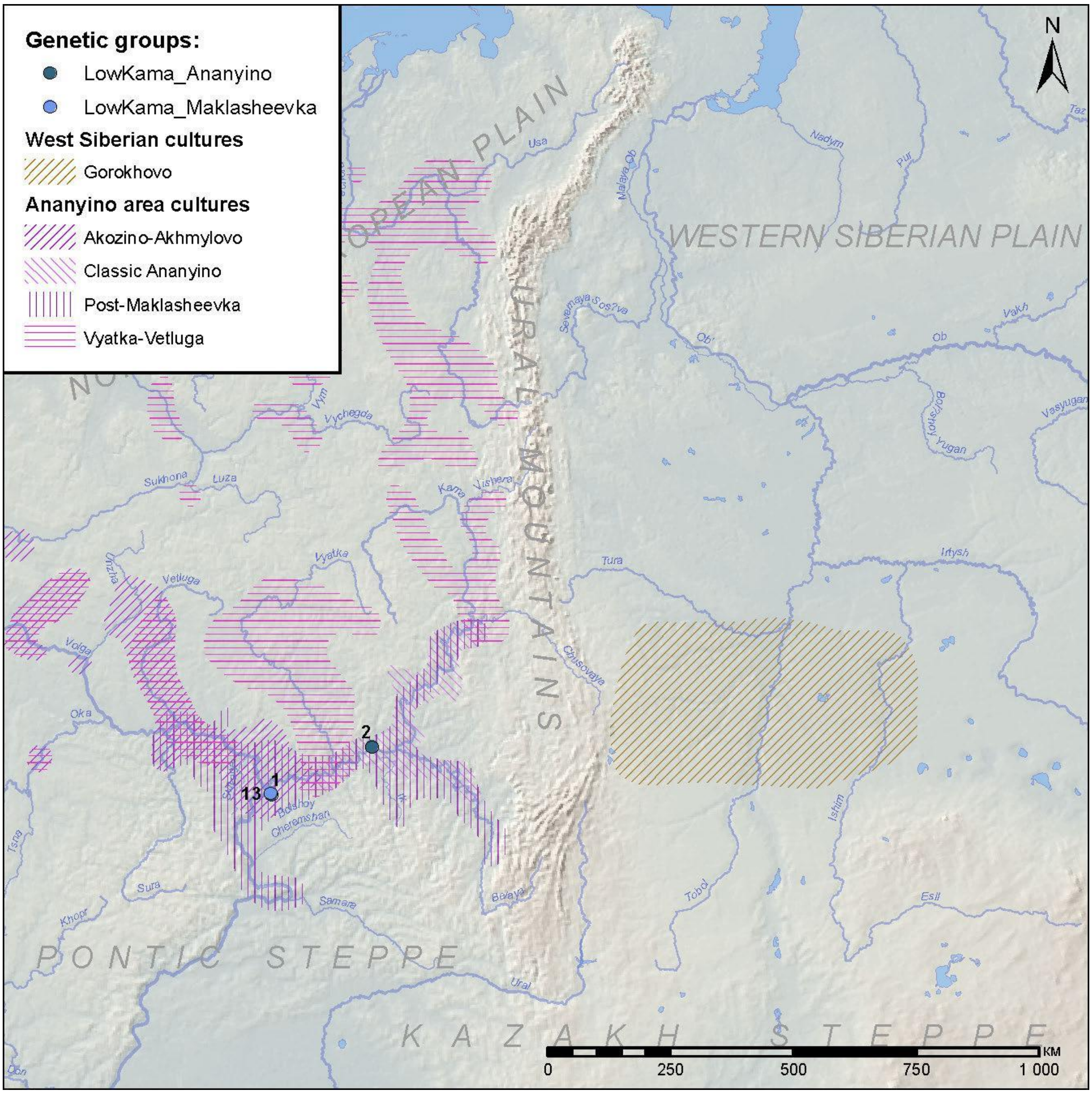

*Fig. SI.A.2. Archaeological cultures existed during phase 1 of the Early Iron Age in the Circum-Uralic region and the location of sites where samples were collected.*

In the Trans-Urals, the period is described as the Gorokhovo-Sargatka horizon (Botalov 2016) or as the time of the coexistence of the two cultures, the Gorokhovo and the Sargatka ones (Matveeva, 2019). The end of the period is marked by the disappearance of the Gorokhovo tradition from the Trans-Ural forest-steppe and emergence of its elements in the Cis-Ural-steppe Sarmatian population (Matveeva, 2018a).

To the north of the Gorokhovo and Sargatka area, a distinct kernel of bronze metallurgy developed in the area of the Itkul culture. The artifactual assemblage shared a common cultural subbase with the Ananyino people and interacted with the West Siberian groups which they supplied with metals (Savelyev, 2007).

#### I.A.2. Early Iron Age, phase 2 (250/200 BCE – 100/150 CE)

The second phase of the Early Iron Age in the Circum-Uralic region coincides with the Sarmation period in the Eurasian steppes. The Sarmatian population intensively interacted with their northern forest-steppe and forest periphery, mediating in trade exchange with the southern craft centers and shaping fashion.

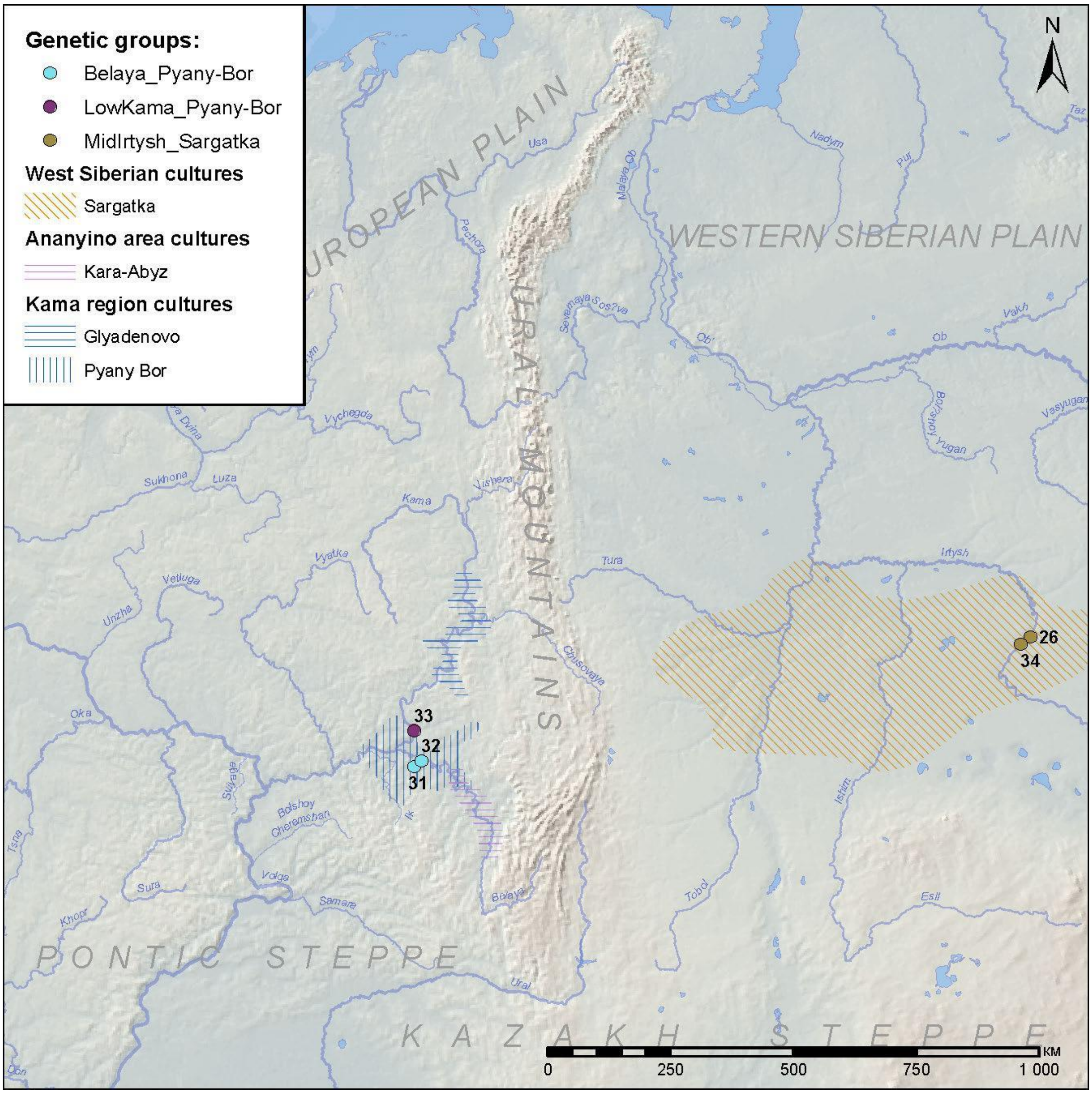

*Fig. SI.A.3. Archaeological cultures existed during phase 2 of the Early Iron Age in the Circum-Uralic region and the location of sites where samples were collected.*

The Trans-Urals and the West Siberian forest-steppe during the 2nd phase of the EIA were dominated by the Sargatka people, who reached their peak in social and economic development during that period (Matveeva, 2018a).

In the Cis-Urals, the Lower Kama was occupied by the Ananyino-rooted groups, attributed to the Pyany Bor culture. Further to the north, the Pyany Bor tradition was neighbored by the Glyadenovo one.

In the Mid-Volga, the period is represented by supposedly small populations of the Gorodets and early Old-Mordovian cultural contexts.

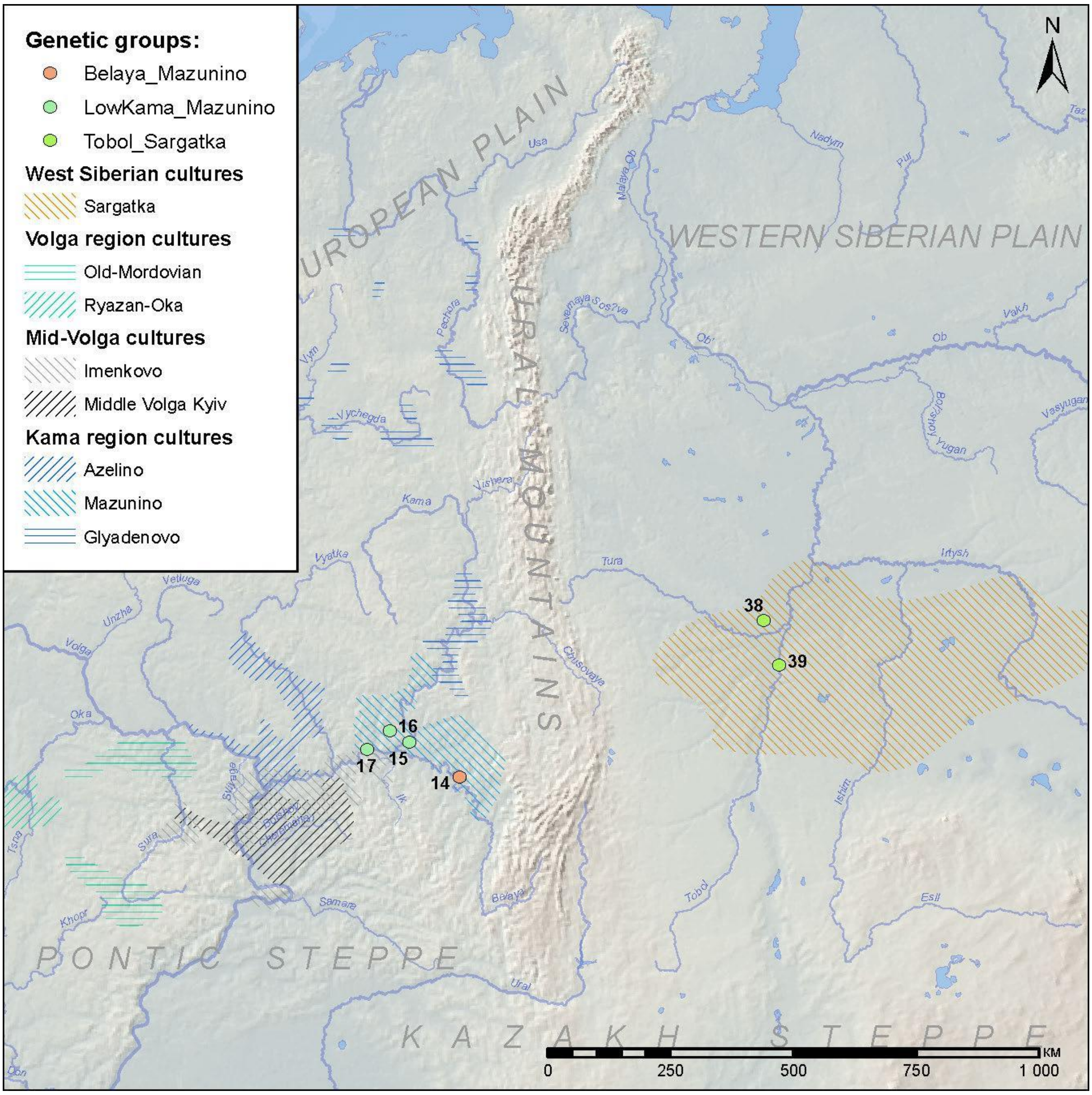

*__Fig. SI.A.4.__ Archaeological cultures existed the period of Roman influences in the Circum-Uralic region and the location of sites where samples were collected.*

The period of Roman influence for the Uralic region is a conventional term, marked by the domination of the Late Sarmatian culture in the steppes, which transmitted styles and innovations from the Mediterranean-Caspian civilization to the boreal zones of the Volga and Kama regions.

The distribution of new categories of imports and subsequent development of new local styles is reflected in regional cultural nomenclature, where the period from 100/150 to 350/400 CE is described as the coexistence of two Pyany Bor-derived cultures, the Azelino in the Vyatka basin, and the Mazunino in the Lower Kama and the Belaya basins. In the Middle Kama region, cultural development was more gradual, and the Glyadenovo culture continued to its late phase. Dramatic changes in the population pattern of the Mid-Volga forest-steppe from 100/150 CE resulted in the formation of several specific cultural groups, possibly reflecting not only cultural transmission but also a migration from the west to the east. All these groups (Kyiv, Early Imen'kovo, and Lbische) intensively interacted with the Late Sarmatian populations of the Volga-Urals.

In the regions to the east of the Urals, the period is associated with the late phase of the Sargatka culture, strongly influenced by the cultural elements from taiga (the Kulai culture). The details of cultural history are under discussion (Botalov 2016). Our periodisation is based on Matveeva (2017) and the set of the radiocarbon dates she collected.

The cross-Uralic cultural relationships during this period were scarce but already existed. In the Bolshoy Cheremshan River valley, a hillfort was excavated with a Late Sargatka-Bakalda architectural style house, dated to the latest edge of the Roman or the earliest phase of the Migration period (Stashenkov, 2020).

##### I.A.4. Migration period (350/400 – 650/700 CE)

The term Migration period is used to cover the Hun (Central European chronology) and post-Hun (Shipovo) periods of the East European steppes and the corresponding phases of development of the local forest-steppe cultural groups. In the Cis-Urals, this archaeological epoch is represented by the classic phase of the Imen'kovo culture in the Mid-Volga, the Late Azelino culture in the Vyatka basin, and the Bakhmutino phase of the Mazunino tradition in the Belaya basin.

In the Mid-Kama basin, the earlier part of the period is usually described as the "Kharino horizon" of the introduction of the Hun-period innovations, while the last stage is marked by the formation of local cultures: the Nevolino, the Polom, and the Lomovatovo.

In the areas to the east of the Urals, we use the term “Migration period” to determine the time span when the interaction of the Late Sargatka and taiga-zone traditions shaped the Bakalda and the Karym-phase Nizhneobskaya cultural groups in the Tobol basin, and the Potchevash one - in the Middle Irtysh area. In the boreal Urals, the period is represented with local population groups, which developed peripheral traditions of the Itkul metallurgists of the Early Iron Age.

Not later than around 550 CE the Cis- and the Trans-Urals manifested an increase in intra-regional cultural connections reflected in the spread of the Kushnarenkovo-type pottery. In the Belaya basin of the western South Urals, sites with the Kushnarenkovo-type pottery form a distinct cultural area, while further west in the Kama region and to the east of Urals it associated with a mixed type of pottery usually found in various contexts (Ivanov 2008; Zelenkov, 2017). Explanations of these processes differ greatly among various researchers and involve different models of interaction: ethnic, cultural, chronological.

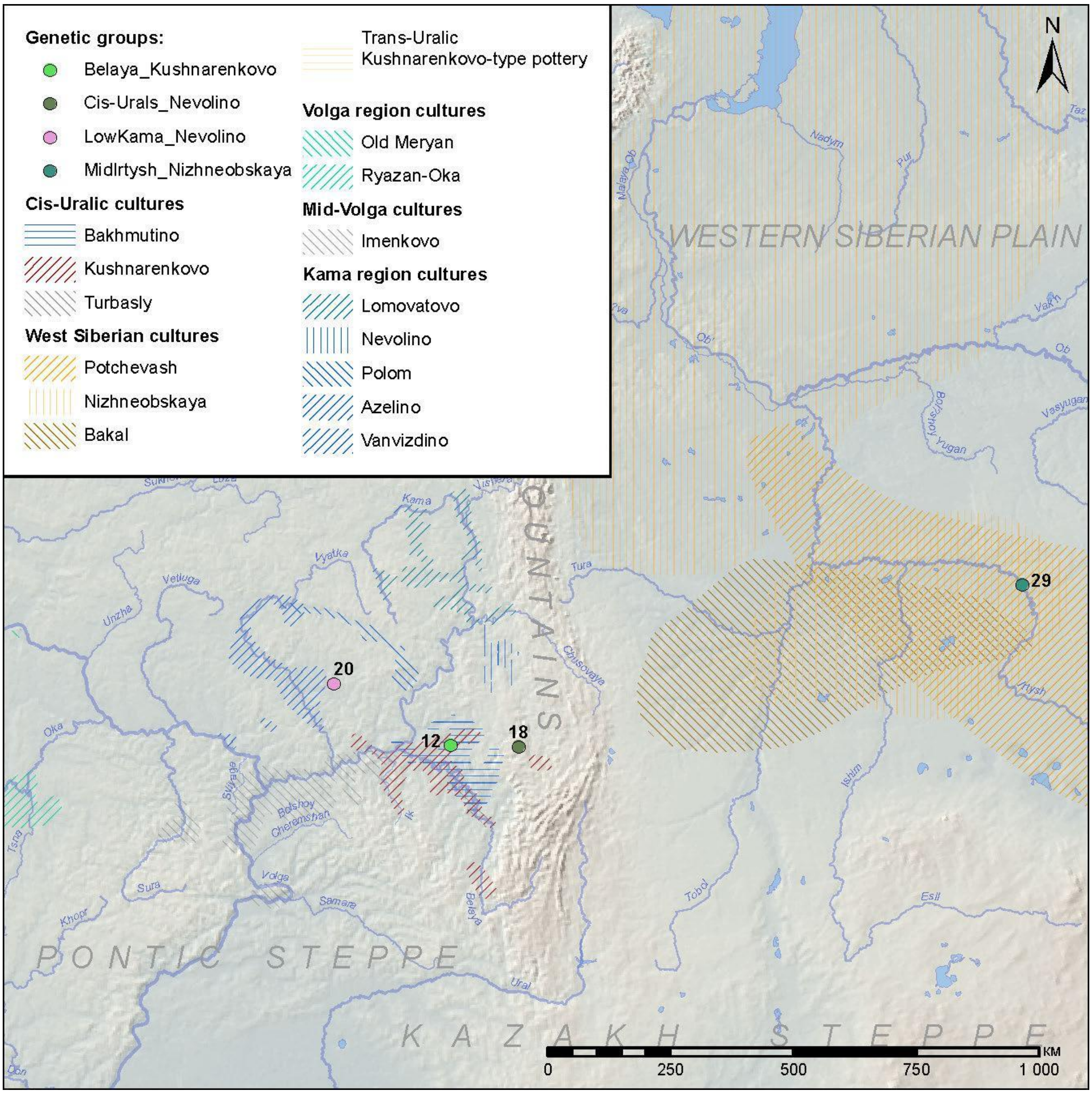

*Fig. SI.A.5.* *Archaeological cultures existed during the Migration period in the Circum-Uralic region and the location of sites where samples were collected.*

Kushnarenkovo-type ware is often combined in burial complexes with belt decorations of the "heraldic" style of the 6-7th centuries, while such cases are extremely rare for Karayakupovo-type ware. However, in burial grounds of the 8th-9th centuries in the Urals and Kama region, the Kushnarenkovo-type ware did not disappear but evolved into forms with poorer decoration and often combined with the Karayakupovo-type ware.

It is difficult to give an unequivocal answer whether this picture reflects the migration and infiltration of groups of the Karayakupovo population into the Kushnarenkovo area, or whether the spread of Karayakupovo-type ware was the result of some cultural processes. This is, in particular, the consequence of the difficulty in determining the original habitat of the Karayakupovo group in the 7th century which still remains hypothetical. However, the fact of the very close mutual influence of the Kushnarenkovo and Karayakupovo groups in the 8th-9th centuries prompts researchers to conclude that they are unconditionally related.

Simultaneously with the formation of the Kushnarenkovo horizon, some traces of cross-Uralic cultural interactions emerge in the boreal parts of the studied region. They are marked by finds of the forest Trans-Uralic comb-pit ware at some Nevolino sites (Pastushenko 2004), and penetration of the West Siberian population to the west of the mountain ridge.

Since these processes covered a wider area than the actual population concentration areas of the Kushnarenkovo and Karayakupovo groups, it is worth determining the Kushnarenkovo and Karayakupovo horizons (stages) in the archaeology of the Cis-Urals.

##### I.A.5. Karayakupovo Horizon (700/750 – 950/1000 CE)

We use "Karayakupovo horizon" to refer to a period when a specific trans-Uralic cultural area formed, connecting the Tobol and the Belaya basins and the northern parts of the Kazakh steppe. The detailed archaeological definitions of the period are still discussed (Ivanov 2022) and the researchers use various terms to define local Circum-Uralic cultural groups of the period (Petrogrom, Karayakupovo, Mryasimovo type, Early Chiyalik, etc.), but their cultural similarity and the proximity to traditions of the Trans-Urals are not disputed.

In the Mid-Volga basin, the formation of the Karayakupovo horizon was preceded by the collapse of the sedentary Imen'kovo-culture population and the inclusion of the area into a northern periphery of the Khazar Khaganate. The earliest penetration of the nomadic groups that are believed to be warbands of the Khazarian border guards (Novinki type) (Stashenkov, 2013) was followed by a more substantial migration process, commonly associated with the Turkic-speaking Early Bulghars (Vyazov et al., 2019).

The cultural influence of the Khazar Khaganate (reflected in the Saltiv-Mayaky archaeological culture) involved the population of the Lower Kama and the Cis-Urals as well and was reflected in a specific assemblage of the Nevolino and the Polom cultures in their late phases. In the remote forest areas, the local cultural groups were less influenced by the southern steppe styles, and the Lomovatovo culture mostly continued to develop the Kama traditions (Goldina 2022).

The Samara, Belaya, Upper Ural, and Tobol river basins in 700/750 – 800/850 CE experienced cultural unification and can be described as a core area of the Karayakupovo traditions. Those ones developed the pottery styles that were first attested as the Kushnarenkovo ceramic assemblage, but in contrast to the Kushnarenkovo period, after 650 CE, the Karayakupovo pottery is represented not only as an admixture to the other pottery types but as a dominating one in cemeteries, on both sides of the Urals.

In the 9th century, the population associated with the Karayakupovo traditions and its cultural descendants probably reached its maximum size. The appearance of the Karayakupovo-type ware is observed both to the east of the Ural Mountains in the Uelgi cemetery (Botalov 2017), and to the west of the mountain ridge, where such ceramics coexisted with the latest Kushnarenkovo pottery in the Bolshiye Tigany site (Khalikov 2022). Several single graves in the Samara Valley part of the Middle Volga rerion were also attributed to the Karayakupovo horizon (Stashenkov 2020).

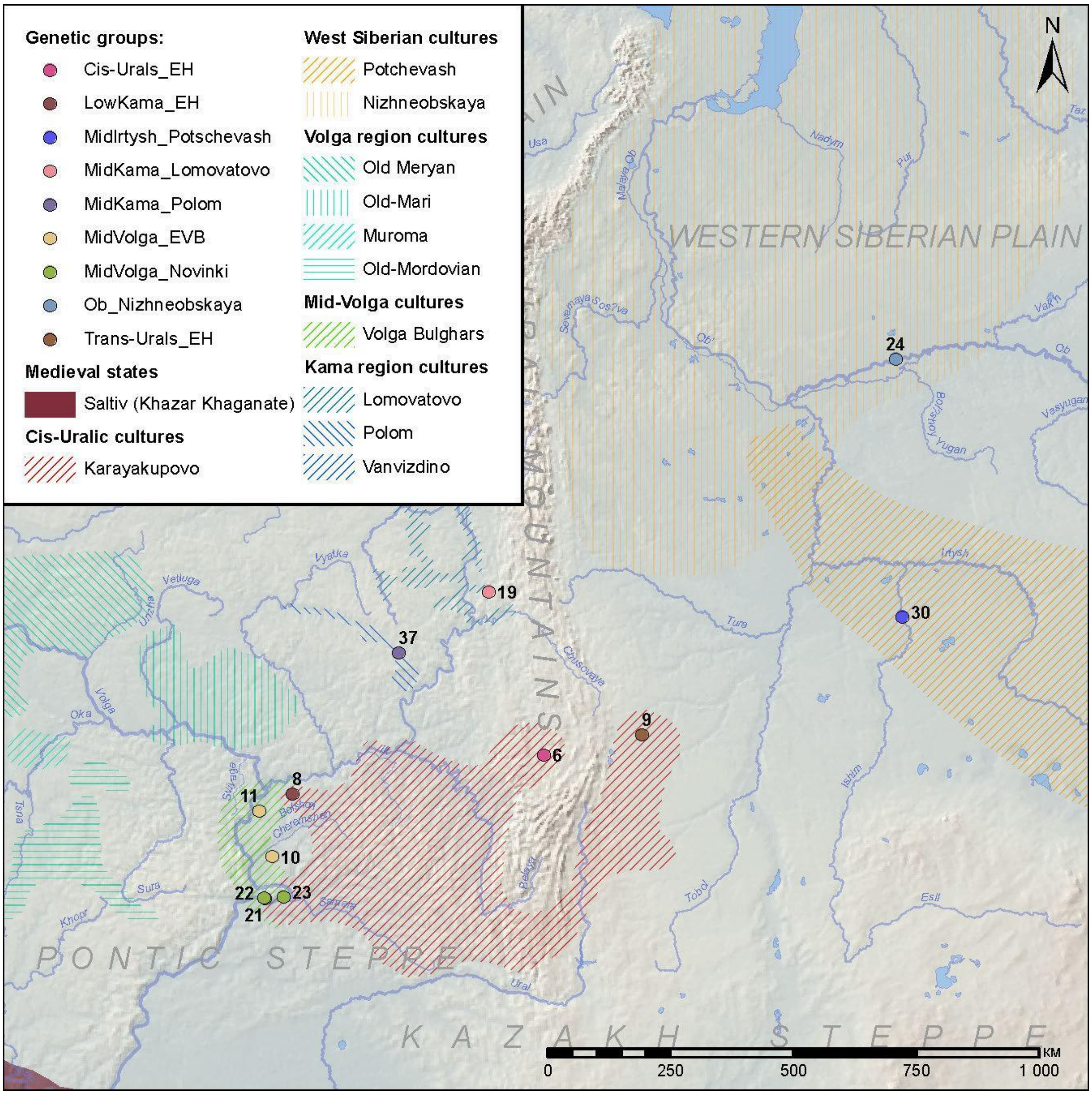

*Fig. SI.A.6. Archaeological cultures existed during the Karayakupovo horizon period in the Circum-Uralic region and the location of sites where samples were collected.*

The appearance of the Karayakupovo-type ware at this time is observed both to the east of the Ural Mountains, e.g. Uelgi cemetery (Botalov 2017), and to the west, up to the Lower Kama region, where such ceramics together with late Kushnarenkovo ware were discovered on the Bolshiye Tigany cemetery (Khalikov 2022). Also several single graves from the Volga region (in the Samara Valley) to the Karayakupovo-type were attributed (Stashenkov 2020).

During the 9th century, the cultural influence of the Karayakupovo traditions involved various neighboring populations, including the ones archaeologically represented by the Lomovatovo and Polom cultures, and the Early Bulghars in Tankeevka.

On the eastern side of the Urals, the cultural development did not experience dramatic changes. In the Mid-Irtysh area, from the 8th century, the influence of the Srostki cultural group on the

Potchevash traditions is recorded, but it was weaker in the boreal zone of the Urals, in the areas occupied by the Nizneobskaya and, later, Yudino cultural areas. The influence of Srostki in also sufficient in Uelgi, where it effectively prevails in the complex of horse equipment and belt decorations. In the Southern Urals, this impulse also dominates from the second half of the 9th century, completely displacing both the Saltiv influence and local types of bridle and belt decorations.

By the 10th century, the common Circum-Uralic cultural environment formed by the Karayakupovo horizon started to disintegrate. This process was possibly driven by the Early Hungarian migration to the Carpathian basin and the occupation of the North Kazakh steppe by the Central Asian nomads, usually attested to as Oghuzs and Patzinaks (Ivanov 2022) or Kimaks and Kipchaks (Botalov 2017). In the southern Cis-Urals of the second half of the 9th century - the 10th century the picture of the influence of both the Srostki culture and the Oghuz-Patzinaks culture is observed in burial complexes, while the specific Ural ware and personal ornaments gradually disappeared during the 10th century. These processes likely mark the Turkization of the region’s population.

##### I.A.6. Middle Ages in the Volga-Ural and Trans-Ural regions (950/1000 – 1400/1500).

Due to the asynchronous socio-political development of the main part of Europe and the Volga-Ural region, we conventionally start the Medieval period with the establishment of Volga Bulgaria in the Lower Kama and the Mid-Volga basins in 900/950 CE. This turning point coincides with the start of the decline of the Karayakupovo Circum-Uralic environment and the formation of continuous and stable cultural groups in the Cis- and Trans-Uralic boreal zones. The development of these groups continued mostly gradually, with a catastrophic breaking point caused by the Mongolian invasion in the 1230s, which, however, impacted mostly urban centers and southern regions on the border of the steppe.

From the 10th century onwards, the cultural development of the Circum-Uralic area was highly influenced by the centers of technology and craft that emerged at the confluence of the Kama and the Volga. The main innovations included the spread of arable farming and trade; from the 10th century, the Great Volga trade route started connecting the remote areas of the East European North with Iran and Central Asia.

Despite the formation of Volga Bulgaria, the Belaya River basin and the eastern part of the Low Kama region were not developed by sedentary agriculturalists, and the traditions of urban life were not introduced there. Those areas were occupied by the people of the Chiyalik culture, commonly interpreted as descendants of Karayakupovo-horizon populations. The chronological frame of the Chiyalik tradition covers both the pre-Mongolian and the post-Mongolian eras.

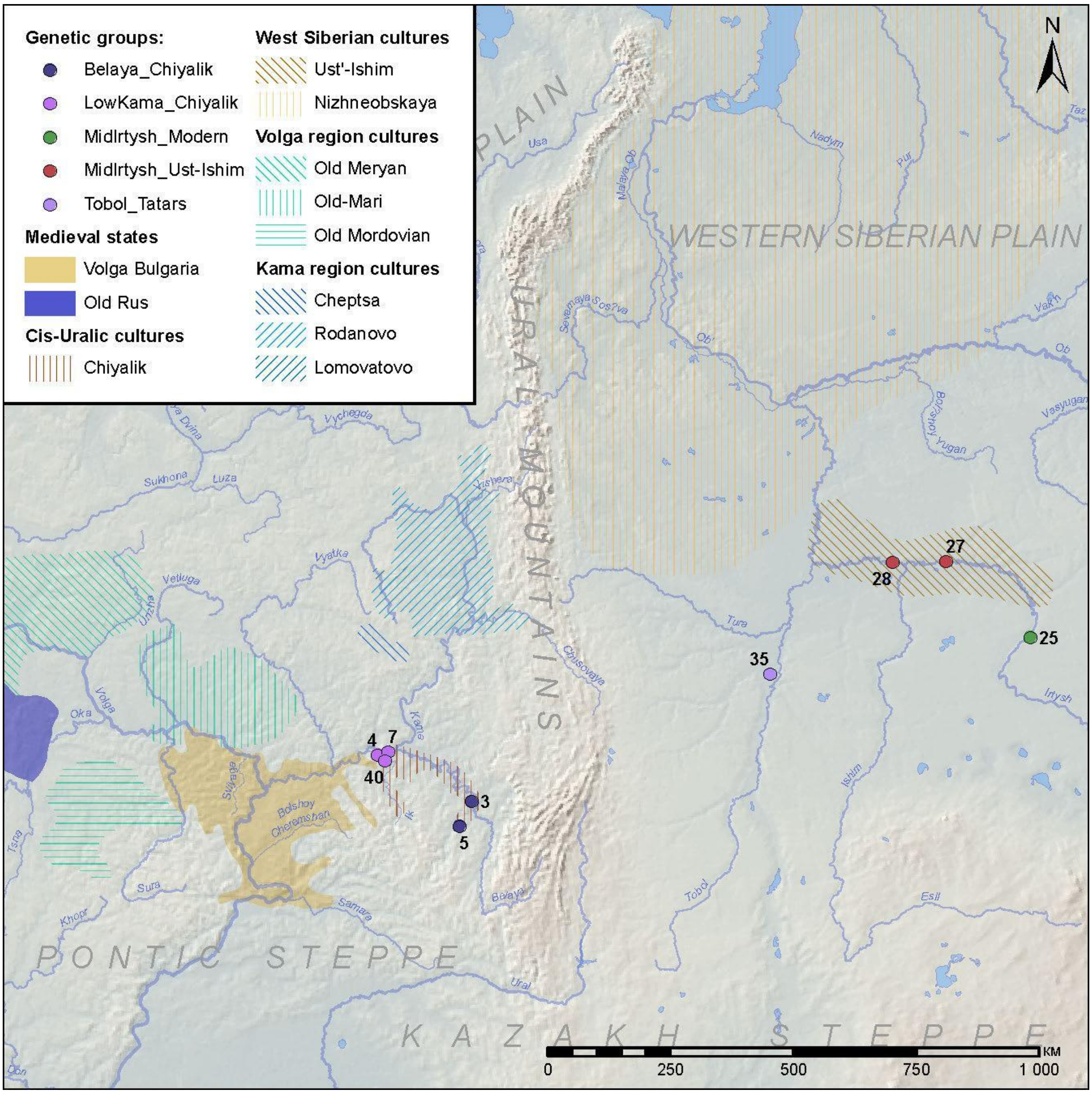

*Fig. SI.A.6. Archaeological cultures existed in the Middle Ages in the Circum-Uralic region and the location of sites where samples were collected.*

In the Mid-Kama region, the cultural tradition called Rodanovo derived from the Lomovatovo, and the Cheptsa culture succeeded the Polom one.

Only during the gradual collapse of the Golden Horde in the 15th century, peripheral populations of the former imperial space transformed into the ethnic groups recorded by the late medieval and early modern travelers and naturalists: the Kazan Tatars, Bashkirs, Udmurts, Khanty, and Mansi.

***Archaeological sites labelled on the maps Fig. S2-7: 1 - Izmeri-7; 2 - Rysovo-1; 3 - Gornovo; 4 - Gulyukovo; 5 - Novo-Khozyatovo; 6 - Karanayevo; 7 - Zuyevy-Klyuchi; 8 - Bolshie-Tigany; 9 - Uyelgi; 10 - Mullovka; 11 - Tankeyevka; 12 - Bustanaevo; 13 - Devichiy-Gorodok-4; 14 - Birsk-2; 15 - Boyarsky-Aray; 16 - Dubrovsky; 17 - Turaevo-1; 18 - Bartym; 19 - Bayanovo; 20 - Sukhoy-Log; 21 - Brusyany; 22 - Malaya-Ryazan'; 23 - Novinki-1; 24 - Barsov-Gorodok; 25 - Borovyanka-17; 26 - Borovyanka-18; 27 - Ivanov-Mys-1; 28 - Panovo; 29 - Ust-Tarsk; 30 - Vikulovo; 31 - Kipchakovo; 32 - Starokirgizovo; 33 - Tarasovo; 34 - Bogdanovo-2; 35 - Putilovo; 36 - Mellyatamak-3; 37 - Varni; 38 - Ipkul'; 39 - Starolybaevo-4; 40 - Ust-Menzelya. Maps in higher resolution are shown as Supplementary Figures S1-S6.***

### I.B. The Middle Volga and Lower Kama

#### I.B.1. The Late Bronze Age Maklasheevka culture (LowKama_Maklasheevka)

##### I.B.1.1. Devichiy Gorodok-4

The Devichy-Gorodok-4 burial ground of the late stage of the Maklasheyevka culture was excavated over the course of four years (1981, 1985, 1989, 2001) by Evgeny Kazakov.

Despite its small size (about 20 burials), its materials significantly changed modern ideas about the funeral rites of the Maklasheevka culture. New details of cultural traditions and artifacts, recorded at the site for the first time, include a cluster of four horse legs, laid in the infill of a grave pit, and a bone bridle set (burial 16), which, in addition to the widespread three-hole rod cheek-pieces, contained additional elements of the harness, such as rings and a distributor made from the shuttle bone of a horse (Kazakov, 2002; Chizhevsky et al., 2019).

From Devichiy-Gorodok-4 burial site, we sequenced one individual from **burial 21** (individual ID I32628, Male).

#### *I.B.2. The Early Iron Age Post-Maklasheevka culture of the Ananyino area (LowKama_Ananyino)*

The first archaeological records later attributed to the Ananyino group of cultures became known in the 1850s. A significant accumulation of data is associated with the preparation of the 4th Archaeological Congress in Kazan, Russia, in 1878 and the organization of the Society of History, Archeology, and Ethnography at the Kazan Imperial University. In the 20th century, A.V. Zbrueva (1952), A.Kh. Khalikov (1977), and V.S. Patrushev (1984) summarized previously accumulated data and elaborated reconstructions of cultural development of the population, associated with the Ananyino sites and artifacts.

S.V. Kuzminykh and V.N. Markov (2007), and A.A. Chizhevsky (2008) presented modernized concepts of the Ananyino cultural complex, emphasizing its heterogeneity. According to their models, the term "Ananyino" covers an area, where four different cultural groups developed separately from different local sources and are characterized by a specific pottery tradition, but share some common traits (Kuzminykh S.V., Chizhevsky A.A., 2009).

1. The Post-Maklasheyevka culture occupied the Lower Kama and the Belaya River valley. This group developed the traditions of the Volga-Kama Maklasheyevka culture of the Terminate Bronze Age. The Post-Maklasheevka bronze celt-axes had a lens-shaped or oval cross-section of the sockets (Ananyino type). The pottery lacked ornamentation, and only at the latest stage, the population started to decorate vessels with sparse cord imprints.

2. The Akozino-Akhmylovo culture spread in the areas of modern Chuvashia and Mari El as well as in the lower reaches of the Oka. The cultural assemblage demonstrates a mixture of elements characteristic of the "textile ware" cultural area in Central Russia and traits of the Maklasheevka culture of the Volga-Kama, the Western component dominated. The Akozino-Akhmylovo cultural assemblage is indicated by celt-axes with a long socket and one ear (Akozino-Mälar type), iron and bimetallic pins, and pottery with imprints of textile or network-like stamp.

3. The Classic or the Corded Ware Ananyino culture is characterized by multi-row corded ornamentation on ceramics and Ananyino-type celt axes with a hexagonal cross-section of a socket developed from local Late Bronze Age cultures in the Middle and Upper Kama region.
4. The Vyatka-Vetluga, or the Comb-Corded Ware Ananyino culture, occupied the vast region from the Vetluga and Vyatka rivers in the south to the Pechora and Vychegda basins in the north. The assemblage of this group ascended to the local Late Bronze Age Lebyazhskaya culture and yielded mostly bone artifacts while the iron smelting was poorly developed.
The chronology of the Anayino cultural area is based mostly on relative dating. Stage 1 is marked by the appearance of Koban (Koban-3) types of artifacts and dates to 900-700 BCE. Stage 2 is associated with the spread of Caucasian imports from the Novocherkassk and Chornohorivka groups during 725-650 BCE. Stage 3 is characterized by Scythian, Sauromatian, Saka, and Itkul imports and dates to 650-400 BCE. Later periods lack imports relevant for precise dating. The Lower Kama region and the Vyatka basin were depopulated at that time, and in the Perm Cis-Urals they possibly continued to exist, but this assumption of the local archaeologists remains putative.

##### I.B.2.1. Izmeri-7

The site is located 2 km northwest of the village of Izmeri, Spassky district, Tatarstan, near the edge of a promontory, separating the vast floodplains of the Kama and the Volga rivers, now flooded by the Kuybyshev reservoir of the hydroelectric station. It was discovered by E.P. Kazakov in 1985 and excavated during 1986-1990. Most of the burial ground was destroyed by the reservoir.

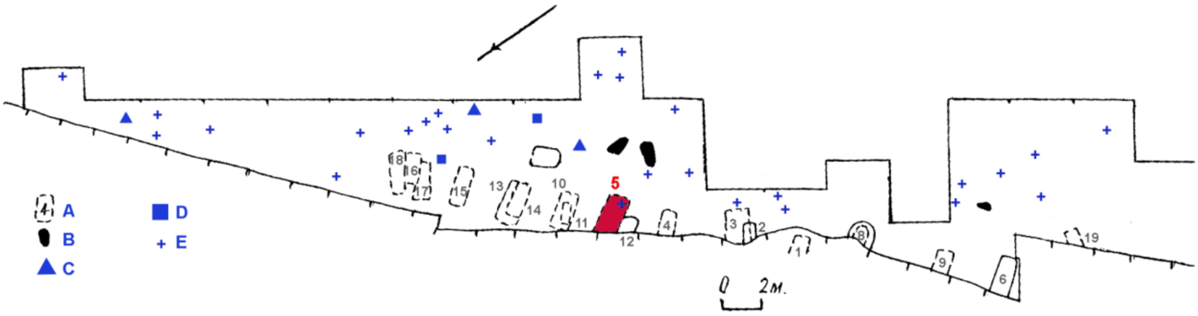

***Fig. SI.B.1.*** *The burial ground Izmeri-7 (16, 17). A plan of the excavated area. The burial 5 is marked in red.*

The burial ground, denoted as Izmeri-7, Izmeri-16, or 17 in different publications, occupies an eroded edge of the former river terrace, elevated for 8-9 m above the reservoir level. E.P. Kazakov collected artifacts from destroyed burials on the shallows under the cliff and traced exposed profiles of grave pits on it. He also unearthed about 30 graves in an excavation area of 283 square m (Kazakov, Lyganov, 2014; Kazakov, 2017). One more paired burial numbered 20, was excavated in the cliff of the bank 20 m northeast of others.

##### burial 5(II) (individual ID I32632, Male)

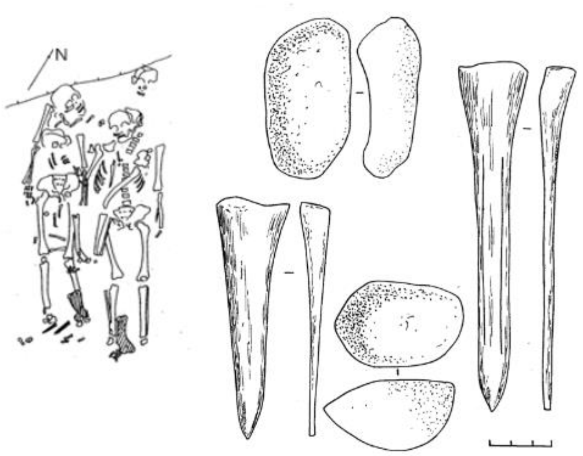

*__Fig. SI.B.2.__ Burial 5 at Izmeri-7 site.*

In the collective burial 5, the bones of a child and an adult were laid over two skeletons of adults. The upper layer of the skeletons reflects a secondary burial, the positions and the orientations of the skeletons do not coincide.

#### I.B.2.2. Rysovo-1

The Rysovo archaeological ensemble was discovered in 1985 by E.P. Kazakov (Rysovo-1 burial ground on the island part) and V.N. Markov (Rysovo-3 occupation site on the bank of the reservoir). Since 1995, the Rysovo sites have been inspected annually by A.A. Chizhevsky. In 2001, he discovered a new burial ground there, Rysovo-2.

Rysovo-1 burial ground is located 8-8.5 km southwest of the village Salaush, Agryz district of the Republic of Tatarstan, 3 km to the southwest of the Rysovo-3 occupation site and 12 km to the east of the village Izhevka, Mendeleevsky district of the Republic of Tatarstan. The site was situated on an elevated remnant of a river terrace, at the height of 8-10 m above the floodplain, at the confluence of the river Izh and the Kama. After the filling of the Nizhnekamsk reservoir, the site became an island, surrounded by water. In the eastern part of the island, E.P. Kazakov discovered Burial 1 in 1985.  A.A. Chizhevsky later found another burial and called it Burial 2.

##### burial 2 (individual ID I32630, Male)

The boundaries of the grave pit have not been traced. At a depth of 55 cm, a skull was recorded, lying with his head to the northeast. The postcranial bones were destroyed by the bank abrasion. Possibly, the skeleton was laid in a supine position with his feet towards the river. I.R. Gazimzyanov examined the skull and attributed it to a young woman who died as a result of damage to the cervical vertebrae. No grave goods accompanied the burial in situ, but a bronze cast plaque in the form of a griffin with an eyelet on the reverse side was found under the cliff. The plate is attributed to the Ananyino period. As far as the pottery vessel unearthed from Burial 1 by E.P. Kazakov belongs to the Ananino Corded Ware tradition, we date Burial 2 to the Ananyino period as well.

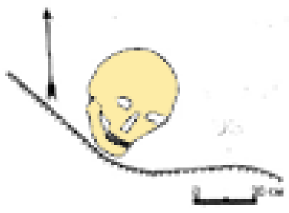

***Fig. SI.B.3***. Burial 2 from Rysovo-1 site.

##### *I.B.3. The Early Iron Age Pyany Bor culture in the Lower Kama area (LowKama_Pyany-Bor)*

The first findings later attributed to the Pyany Bor culture became known in 1878. A significant volume of materials was acquired as a result of A.A. Spitin's extensive expedition throughout the Vyatka Province in 1898. Initially, all post-Scythian period sites in the Perm Cis-Urals were unified within a single culture. It was subsequently designated as an independent culture by A.V. Schmidt in the 1920s. Key consolidations were achieved through the dissertation research of V.F. Gening (1959; 1970; 1971), B.B. Ageev (1983; 1992), D.G. Bugrov (2006), and R.R. Sattarov (2019).

Burial grounds have been extensively studied, they are flat and have an average size of 200-400 graves. Burials were arranged in rows, in shallow, often simple pits, sometimes accompanied by poorly preserved wooden structures of uncertain form. Adjacent to the skeletons (extended in a supine position), numerous costume ornaments (worn during life) were found, alongside tools and weapons.

Finds and features of settlements have scarcely been described, largely due to the peculiarities of layer formation and the poor methodologies of early excavations. Economic aspects remain understudied, although evidence suggests advanced animal husbandry and hunting practices. Agriculture, while probable, lacks detailed investigation. Pottery vessels are handcrafted, round-bottomed, and ornamented with belt-like rows of pits around the neck.

Chronology: The core of this culture's formation lies within a local group of sites in the lower and middle reaches of the Syun' River (a left tributary of the Belaya River, itself a left tributary of the Kama River). These sites are challenging to date precisely. The Ik burial ground, located significantly apart from the "core," provides insights into early complexes but poses difficulties in attribution. The inception of the culture may be traced to 150 BCE. A chronological gap exists between it and the preceding Ananyino cultures. A distinct period of peak development for the classical Pyany Bor culture is evident in 200-225 CE, while the delineation of materials from 100 BCE to the 50 CE is more intricate. A substantial portion of the 1st century BCE likely comprises unadorned burials, identifiable primarily through their planimetrics.

During 200-250 CE, some of the population remains in their previous habitation areas, gradually transforming into the Mazunino culture. Simultaneously, another segment shifts westward, founding sites associated with the Azelino culture.

###### I.B.3.1. Tarasovo

Tarasovo village, Karakulinsky District, Udmurt Republic. Located on a high promontory of the right bank of a stream, a tributary of the Tarasovka River, a right tributary of the Kama River. Discovered by N.L. Reshetnikov in 1979 during the installation of an oil well, 1 km southeast of the village of Tarasovo. The site was excavated from 1980 to 1997 by the Kazan Military-Archaeological Expedition under the direction of R.D. Goldina. A total of 1880 burials were studied over an area of 16,000 square meters. This is the largest Finn-Ugric burial ground of the first half of the 1st millennium CE. The site represents a transition from the early Cheganda culture to the Mazunino culture. Burials are arranged in rows with varying orientations. Numerous weapons were found, including swords, spearheads, and arrowheads, as well as defensive items such as helmets, armor, and chainmail. Some burials contained "sickles-humps". Ornaments include plaques, brooches, overlays, pendants used for headwear, chest ornaments, belts, and footwear accessories, as well as clothing adornments. Many different types of fasteners were found, including buckles, buckles with fixed hooks, belt loops, and butterfly fibulae.

##### burial 731 (individual ID I32623, Female)

The pit has a flat bottom and is oriented along the northeast-southwest line. The upper part was damaged during the installation of an oil well. At the bottom, poorly preserved skeletal remains of a female individual 25-30 years old were found, suggesting that the deceased was placed lying on her back, with her head to the northeast. No artifacts were found.

##### burial 1133d (individual ID I32624, sex unidentified)

The pit has a flat bottom and is oriented along the southwest-northeast line. The upper part of the pit was damaged by plowing. At the bottom, four poorly preserved skeletal remains were cleared. Skeletal remains A: the deceased was laid out lying on her back, arms along the body, head to the northeast. A fragment of a bronze object was found to the right of the skull bones. Skeletal remains B: the buried individual was laid to the left of buried A; the preserved skull allows determining the orientation of the deceased - head to the northeast. No artifacts were found. Skeletal remains C: only the skull remains, indicating that the deceased was laid with the head to the southwest, between buried A and D. No artifacts were found. Skeletal remains D: the deceased was laid out lying on her back, arms along the body, head to the southwest. An iron buckle was found on the pelvic bones, on the right side, and a fragment of an iron knife was found between the femur bones.

Anthropologically, the excavated skeletons were determined as: A - female, 40-50 years old; B - sex?, 8 years old; C - sex, age?; D - female, 25-30 years old.

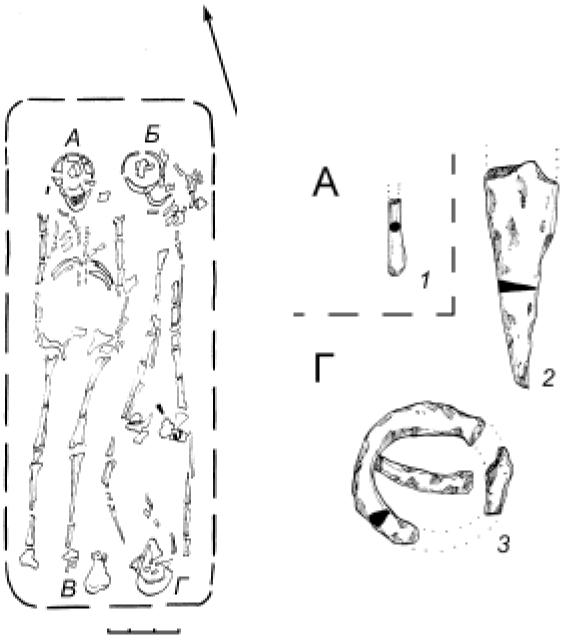

***Fig. SI.B.4.*** *Burial 1133d at the Tarasovo cemetery.*

#### *I.B.4. The Migration period Mazunino culture in the Lower Kama area (LowKama_Mazunino)*

The earliest finds of the artifacts later attributed to the Mazunino culture were first described in the late 19th century, and extensive excavations, culminating in a systematic description of the cultural assemblage, started in the second half of the 1950s (Gening, 1958; 1967). The unearthed data was summarized at the turn of the 1970s/1980s (Ostanina T.I., 1983a/p), but the results (Ostanina T.I., 1998a/p) were only published 15 years later without updates (Ostanina T.I., 1997).

The earliest manifestations of the elements of the Mazunino assemblage are recorded at the late Pyany Bor sites located on the right bank of the Kama River opposite the mouth of the Belaya and dated to 150-200 CE by the Sarmatian-derived elements of belt sets. One can find there the earliest forms of Mazunino-type temple pendants, a rite of placing a belt in the grave not worn by the deceased but stretched along the body develops, characteristic "butterfly-shaped" fibulae evolved from “Sarmatian” prototypes (with a coil at the end of the receiver, Ambroz-13 group).

In 200-250 CE, the Pyany Bor tradition gradually declined, while the Mazunino traditions strengthened. Overall, by the mid-3rd century CE, a classic set of all ritual and inventory features has developed.

The settlement clusters comprise hillforts of two distinct types: smaller ones characterized by insignificant cultural layers, and larger ones with thicker layers, alongside unfortified settlements. Fortifications may entail multiple rows of ramparts and ditches. The habitation areas within these settlements feature rectangular on-surface houses equipped with hearths and cash pits.

Pottery is predominantly handmade, incorporating a mixture of bird droppings, crushed shells, sand, dry clay, and calcined bones in the pottery clay. Vessels typically exhibit flattened round bottoms and rims, occasionally adorned with notches and pinches, while their bodies are embellished with round, square, or triangular corded stamps, arranged in one or several horizontal lines, or less frequently, in clusters of pits.

Cemeteries are characterized by flat grave pits sized according to the sex and age of the deceased, arranged in rows and groups. Skeletons are typically placed in supine positions, primarily within individual graves, occasionally within wooden coffins, or covered by timber overlays or logs. Collective burials have also been documented, and interpreted as family graves. Costume details and jewelry are often found on the bodies of the deceased, although belts are frequently aligned along the body. A common feature of burial practices involves the placement of decorations and small tools in separate niches (referred to as "mortuary assemblage"). Additionally, small vessels were sometimes included as part of grave goods. Finds of weapons and tools are infrequent, predominantly consisting of imported swords and helmets, with approximately 10-30% of burials lacking any inventory.

Costume decorations of the Mazunino culture include various items such as temple pendants, chainmail braid covers, torques, necklaces made of beads and threads, fibulae (including later, more specific types like "butterfly"-shaped, curved "bow"-shaped, and abundantly decorated rectangular-shaped), belt sets, numerous pendants and overlays, as well as knife sheaths typically featuring protrusions and pendants.

The economy of the Mazunino population was primarily based on livestock farming and beaver hunting, likely supplemented by slash-and-burn agriculture.

Around 300-350 CE, the cultural assemblage of the Mazunino underwent significant transformation, possibly triggered by the breakdown of interactions with the steppe Sarmatian population. There was a decline in the number and variety of bronze decorations, while iron artifacts began to increase in quantity. The newly introduced artifacts continued the stylistic development of Late Sarmatian prototypes, which had influenced Mazunino craftsmanship in the period leading up to the breakdown of communication. Concurrently, Late Sarmatian-style weapons disappeared, replaced by common types from the Migration period, which became more numerous and prevalent.

By 300 CE, the Mazunino population had vacated the right bank of the Kama River, with no documented finds of Mazunino-attributed artifacts dating from 300-450 CE in that area. However, a group of sites in the Belaya River valley continued to be inhabited, and influences from the Mazunino tradition persisted in the Middle Kama region (Nevolino culture) and the Cheptsa basin (Polom culture), as well as sporadically in Western Siberia.

In the later stages of evolution, the Belaya Mazunino group introduced innovations in pottery production, resulting in new vessel shapes such as ribbed bodies adorned with stamp decorations (referred to as the "Imendyashevo" type) and ceramics featuring numerous, irregularly arranged or row-like notches across the vessel body (known as the "Bakhmutino" or "Chandar" type) (Ovsyannikov, Sungatov 2004). These innovations coexisted with traditional elements of the cultural assemblage, suggesting ongoing cultural development within the population. However, the origins of these pottery-making innovations remain unclear. The most recent burials associated with the Mazunino tradition date back to 500-550 CE. During this final stage of cultural development, there was a gradual replacement of traditional styles with more "fashionable" items.

It is noteworthy that certain regional archaeological schools tend to identify distinct cultures within the Belaya Mazunino group, primarily based on variations in pottery styles (Belyavskaya, Protsenko, 2018). However, these approaches overlook the typological consistency observed in other categories of artifacts, suggesting the cultural coherence of the Mazunino Belaya group, despite the presence of numerous specific ceramic types.

###### I.B.4.1. Boyarka ”Aray”

The site is located 4 km northeast of the village of Boyarka (Karakulinsky district, Udmurt Republic) and 5 km south of the Kama River, on an elongated promontory of the river bank known as "Aray." The site surface is flat and gently slopes southeastward, measuring approximately 50 m in length and 10-15 m in width, with steep, actively eroding slopes, and ravine bottoms overgrown with forest.

In 2002, N.L. Reshetnikov from the Sarapul Museum collected artifacts from citizens of Boyarka, including iron knives and their fragments, iron temple pendants with bronze wrapping, and 2 bronze “butterfly” fibulae. Rescue excavations at the site were initiated by E.M. Chernykh of Udmurt University, leading to the discovery of 183 burials.

###### burial 155 (individual ID I32631, Male)

The grave pit found at a depth of -30 cm was oriented along the southwest-northeast line. The northeastern part of the grave pit was disrupted by burial 156. The surviving portion of burial 155 had a rectangular shape with slightly concave long walls, measuring 150x80 cm and reaching a maximum depth of 96 cm. At a depth of 90 cm, small traces of decayed organics up to 0.3 cm thick were found. The skeletal remains of the deceased were unearthed at the bottom of the grave pit.

Initially, the deceased was placed supine with the head to the northeast, arms alongside the body. The bones of the upper half of the skeleton were in correct anatomical order except for the rib cage, part of which was found near the feet of the deceased. The leg bones were shifted towards the pelvis, with the tibiae seemingly detached and placed on the femurs. The radius of the right arm of the deceased and several smaller bones of the skeleton were found to the left, near the skull, with the sternum occupying the position of the radius of the right arm.

In the area of the skull, an iron buckle and a bone arrowhead were found on the left side. A fragment of an iron object of uncertain purpose, which disintegrated during cleaning, was found near the pelvic bones.

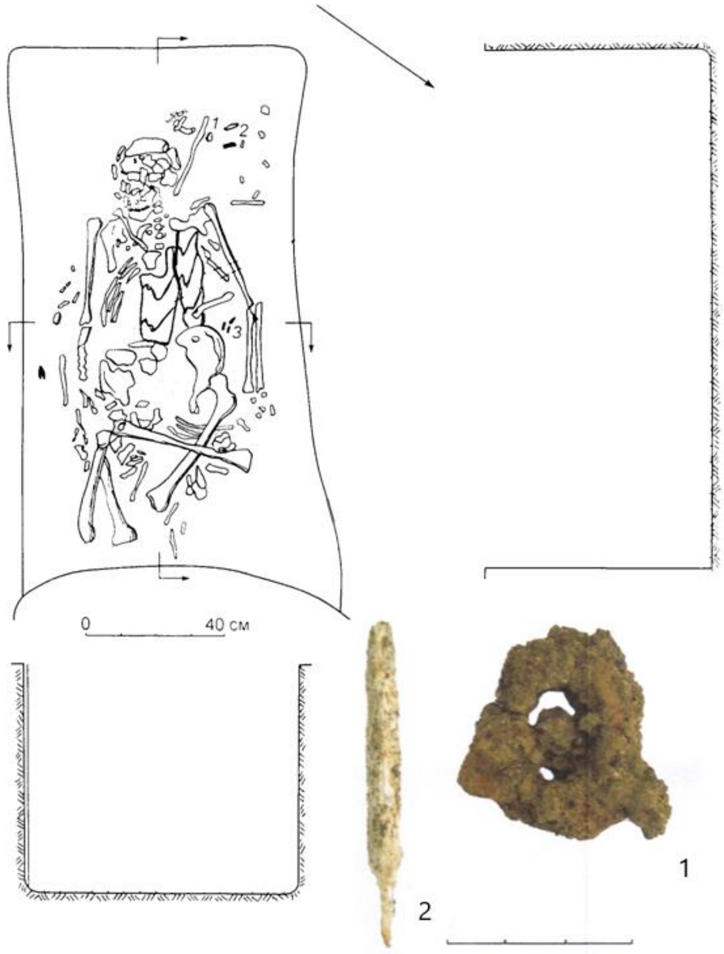

***Fig. SI.B.5.*** *Burial 155 at the Boyarka "Aray" cemetery.*

##### I.B.4.2. Dubrovsky

The Dubrovsky burial ground is located 0.5 km south of the settlement of Dubrovsky and 1 km south of the village of Kalashur in the Kiyasovsky district of the Udmurt Republic, on an elevated promontory of the terrace on the left bank of the Shekhostanka River. At this location, the river makes a sharp loop, creating whirlpools and rapids in the channel. A total of 204 burials have been excavated at the site.

###### burial 52 (individual ID I32627, Male)

The grave pit (excavated in two years, 2010 and 2012) was unearthed at -30 cm from the modern surface. The shape of the pit was rectangular with rounded corners and a slight widening in the northern half, measuring 224x85 cm. The walls of the pit were vertical to the south and west, and inclined to the north and east. The bottom was flattened along the longitudinal axis and flat along the transverse axis. The orientation of the pit was along the southwest-northeast line. The maximum burial depth was -65 cm from the modern surface. At the bottom of the grave, closer to the southern and western walls, a skeleton of satisfactory preservation was cleared. Based on the position of the bones, it can be assumed that the deceased was laid out supine with the right arm along the body, and the legs bent at the knees to the left so that the shins touched. The head was facing south-southeast. Discoveries at the bottom included a fragment of an iron buckle near the right thigh and an iron knife along the femur below the buckle.

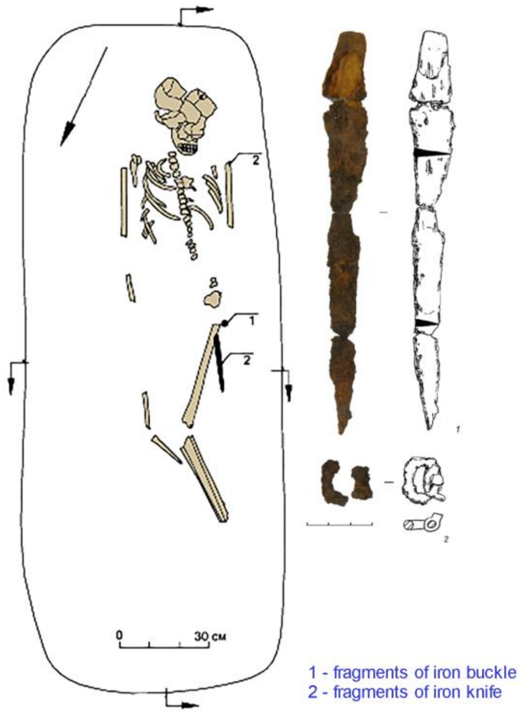

***Fig. SI.B.6.*** *Burial 152 at the Dubrovsky cemetery.*

##### I.B.4.3. Turaevo-1

The burial site is located on the right bank of the Chukma River (also known as Chemek), a right tributary of the Kama River, on an elevated plateau of the terrace gradually descending towards the Kama River, not far from it, to the southwest of the village Turaevo (Mendeleevsky District, Republic of Tatarstan). The cemetery includes two parts, a group of kurgans excavated by Vladimir Gening in 1969, and a flat part, excavated by N.V. Vodolago from 1986 to 1990. The latter excavation of an area of 2688 square meters yielded 269 burials attributed to the Mazunino culture. The burials are arranged in rows and oriented along the southwest-northeast line. Numerous artifacts were found near the skeletons, including belts laid along the body of the deceased, pendants, beads, overlays, bracelets, fibulae of local forms, and clay vessels.
In this paper, we publish two individuals buried in the flat graves of Turaevo, one from burial 66 (ID I32629) and another one from unidentified burial (ID I32621, Male).

###### burial 66 (individual ID I32629, Female)

Anthropologically identified as female, 18-20 years old. The grave pit had sloping ends, vertical walls, a flat bottom, and was oriented along the southwest-northeast line. Judging by the poorly preserved skeletal remains, the deceased was laid out supine, arms along the body, with the head to the southwest. To the left at the head of the grave, a mortuary assemblage was found, consisting of fragments of bronze spiral-threaded pins (7 fragments preserved), glass beads (5 specimens), fragments of fused bronze and iron chains, an iron awl with fragments of a wooden handle (attached to it was a fragment of a bronze spiral-threaded pin), and an iron knife with fragments of a wooden handle and a bronze pin; underneath the items, fragments of birch bark (a round box) were found. In the shoulder area (to the right and left of the mandibula), four iron temple pendants with bronze wrapping were discovered (two specimens on each side). The sleeves of the deceased's garment were adorned with large round bronze pins "beads" (116 specimens preserved), arranged in the area of the arms from the shoulders to the elbows; fragments of fabric and fur were also found there. Bronze and iron necklaces with bronze wrapping lay in the chest area. Additionally, slightly below the necklaces, an iron three-part fibula with bronze hemispheres, iron and bronze pendant chains, was found. Four bronze pins (pipe pins), zoomorphic: a bear pin and two duck pins, were attached to the chains, as well as six pendants, five of which were bronze (3 lobed), one trapezoidal and one zoomorphic pendant (horse), and one square iron pendant. The deceased wore three bronze bracelets with flattened ornamental ends on her arms: two specimens on the right arm and one on the left (found in the hip area). Furthermore, to the left along the body of the deceased, from the chest to the feet, a unfolded leather belt was laid. Preserved from it were an iron buckle with six bronze rivets in the chest area, bronze overlays (a total of 85 specimens), and an assembled tip made of four iron plates (each with four bronze rivets) with four iron pendants; the tip lay between the shin bones. Of the 85 bronze belt overlays, one had two bronze pins, twelve had three bronze hemispheres (nail caps), and the remaining 72 overlays had two bronze hemispheres (nail caps) and a bronze ring. The total recorded length of the belt was at least 95 cm, and the width of the leather base of the belt was 3.0-3.5 cm.

##### *I.B.5. The Migration period and Early Medieval Nevolino culture in the Lower Kama area (LowKama_Nevolino)*

Finds later attributed to the Nevolino culture are known from the mid-19th century. The eponymous site, Nevolino burial ground, was studied in the 1920s and 1950s (materials were published unsystematically). Extensive field research, nearly fully published to date, led to the substantiation of the characteristic features of the monument group, conducted by Rimma Goldina (Goldina & Vodolago 1990).

The sites attributed to the Nevolino culture usually form spatial clusters which include hillforts and unfortified settlements. In the occupation area, separate homesteads are distinguishable, often arranged in rows. Houses were small, semi-subterranean or slightly dug into the ground, square or rectangle with rounded corners in shape. There were two round (diameter 4-5 m) sunken structures with 1-2 main pillars in the center. Sometimes, pillar constructions and porches can be traced. A hearth constructed with stones was usually located in a corner of a dwelling. Pottery was handmade and round-shaped; by proportions, there were pot-shaped, bowl-shaped, and dish-shaped vessels; with or without neck; paired handles-ears were rare. In later periods, the number of ornamented vessels increased; the upper part was decorated with notches, pinches, cord impressions, comb stamps, depressions, incised lines forming bands of horizontal lines, zigzags, fir trees, nets. There were wooden vessels with metal inlays, including those with zoomorphic handles, made on a potter's wheel from southern centers, including the Saltiv-Mayaky culture, and others. Burial mounds (small mounds with ring ditches) around the beginning of the 7th century were replaced by flat cemeteries, where graves formed groups and rows. Deceased were placed in rectangular pits, often in wooden coffins or frames; laid out on the back, the predominant orientation varied across necropolises. The inventory was mainly arranged in the order of wearing, but belts could be stretched along the body, and some items were placed in boxes. Male burials often contained weapons, horse gear (bridle could also be found with women). Artificial cranial deformation is known. There were many pendants in the decorations. Belt set arrangements are typical for the Perm region. Female belts, dated to the second half of the 7th-8th centuries, with wide pendant strips, abundantly decorated with overlays resembling an inverted heraldic shield with slits resembling a face, Ж-shaped overlays, round overlays, "triplets," slotted tips, and others, are attributed to the specific “Nevolino type”. A relatively large number of weapons and horse gear sets distinguish the Nevolino culture from other traditions of the Upper and Middle Kama regions. About 20 hoards

with metal utensils and coins (mostly Sasanian silver; there are also from Central Asia, Byzantium), jewelry, and metal ingots are known. The economy was based on agriculture (slash-and-burn), animal husbandry (horses, cattle, less sheep and goats, few or no pigs), hunting (furs were important), and crafts.
Five phases of the development were distinguished by Rimma Goldina, they are as follows: “Brody”, late 4th-5th centuries; “Verkh-Saya”, mainly 6th century; “Bartym”, late 6th-7th centuries; “Nevolino”, late 7th-8th centuries; “Sukhoy-Log”, late 8th - 1st half of the 9th centuries.
Some elements of the Nevolino pottery derive from the traditions of the Glyadenovo population. The Mazunino component may have had some influence as well. In the mid-9th century, marked by the spread of Saltiv types, the territory became deserted. In the late stage, the spread of typical Nevolino ornaments (belts, pendants) over a vast territory to Finland, Sumy Oblast of Ukraine, Omsk Oblast of Russia, and Kazakhstan was recorded. Sites directly left by late Nevolyinsk groups spread southeast of the original territory, to the headwaters of the Ufa River with its tributaries (settlements of Rakhmangulovo, Abdulinsky, Rakhmangulovskoe, Tornalinsky, Bolshoustikinsky hillforts, Kzytausky, Yangatausky settlements) and southwest (settlements Osh-Pando-Ner, Podgory-1).

##### I.B.5.1. Sukhoy Log

The Sukhoy-Log burial ground is located in the Kishertsky District of the Perm Krai, on the right bank of the Sylva River, a left tributary of the Chusovaya River, which is in turn a left tributary of the Kama River. The site is situated on the southeastern outskirts of the village of Sukhoy Log, on a high 17-meter terrace of the right bank of the Sylva River. At this location, the terrace takes on the shape of a rectangular promontory, formed by the steep bank of the river and a depression perpendicular to it, transitioning into a hollow that cuts through the terrace and defines the platform of the monument from the west. To the east, the terrace gradually rises. The platform of the monument is occupied by buildings and gardens of the modern village.
The burial ground was discovered in 1986 by E.E. Kirzhner. A trench with an area of 9 m2 was excavated at the edge of the terrace, revealing fragments of pottery, bronze, and iron artifacts, as well as two burials (1-2). In the same year, a larger excavation, covering an area of 95.5 m2, was conducted by N.V. Vodolago, where seven more burials were found (3-9). In 1987, 10 burials (10-19) were discovered on an area of 157.5 m2. In 1988, E.E. Kirzhner continued the investigations, excavating an area of 125 m2 and studying three burials (20-22). As a result, over the course of three years, an area of 387 m2 was investigated, revealing a total of 22 burials.

###### burial 13 (individual ID I19111, Female)

A 14-16 years old female was identified in burial 13. Along the presumed southern longitudinal wall of the burial, at the boundary of the mainland, at a depth of 40-56 cm, the remains of a fragmented skeleton were found in disorder. The bones of the pelvis and legs were relatively undisturbed. Based on their position, it appears that the deceased was laid with her head to the northwest. No inventory was found in association with this burial.

###### burial 22 (individual ID I19112, Male)

A male, aged 40-50 years old, was identified in burial 22. The burial was documented at a depth of 100 cm. The grave pit had a rectangular shape, with dimensions of 270x87x118 cm, oriented along the northwest-southeast axis. At the bottom, remnants of a wooden coffin measuring 171x38 cm were found. Upon clearance, a moderately preserved skeleton was uncovered, including cranial bones, partial rib cage, arm bones without hands, and leg bones. Based on the arrangement, it appears that the deceased was laid with his head to the northwest, lying on his back, with the skull facing upwards, arms alongside the body, and legs extended. Vertebrae and likely parts of the scapulae were found at the head. A gold pendant with an amber insert was discovered on the cranial bones, while iron fittings, fragments of clothing, belt buckle, and bronze fittings and tweezers were found around the waist and pelvis area. A bronze collar and scabbard tip, as well as iron knives were found near the right femur. An iron saber was placed near the left foot, an iron axe near the right knee, and an iron buckle near the left knee. Iron spurs and stirrups were detected near the right foot.

#### *I.B.6. The Early Medieval Novinki-type sites (MidVolga_Novinki)*

See Szeifert et al., 2022 for the general description of the Novinki-type sites.

##### I.B.6.1. Mound groups at Brusyany

See Szeifert et al., 2022 for the general description of the kurgan groups at Brusyany.

###### Brusyany-2 mound group

mound 22, burial 3 (individual ID I19085, Female)

At the southern border of the kurgan mound, in the layer of buried soil, poorly preserved bones of a child were found, placed on its back with the head to the west. To the left of the skull, there was a rounded-bottom pot vessel.

mound 23

burial 3 (individual ID I32663, Male)

The burial was unearthed in the northeast sector of the mound, outside the rubblework. The skeleton of a female person lay stretched out on its back, with the head to the northwest in the center of the rectangular burial with rounded corners. The bones of the ankles were strongly pressed together (bound?); the skull was turned to the right side. Near the wrist of the right hand, there was a spindle; near the skull and the right shoulder - bronze pendant balls with loops and bronze open rings-earrings.

burial 5 (individual ID I19086, Male)

Fragments of the infant's skull were found in the northwest sector outside the stone rubblework. The determined radiocarbon age of the burial is 600-666 calAD (1400±30 BP, Poz-136205) (based on human bone).

##### I.B.6.2. Malaya Ryazan'

See Szeifert et al., 2022 for the general description of the sites at Malaya Ryazan’.

burial 4 (individual ID I19082, Female)

The determined radiocarbon age of the burial is 576-654 calAD (1440±30 BP, Poz-136234) (based on human bone).

I.B.6.3. Novinki-1

See Szeifert et al., 2022 for the general description of the sites at Novinki.

mound 7, burial 2 (individual ID I16639, Male)

A burial was performed in a rectangular pit measuring 260 cm in length and 70-80 cm in width. The burial has been disturbed, with the skull broken into two parts. Presumably male, the age of the deceased is estimated to be around 40 years. Anthropological analysis indicates that the bones have not been preserved. Fragments of iron objects (plates) and two bronze overlays were found in the burial.

mound 8, burial 6 (individual ID I19104, Male)

The burial occupies a central position within the mound and is oriented in an east-west direction. It likely intersected another burial pit oriented to the south, or this pit may represent a cenotaph.

A small recess was made in the northern wall of the burial. The skeleton is fragmented, with the bones scattered. Only the lower leg bones remained in place, indicating that the burial was oriented with the head to the west. Two items were found in the burial: a piece of charcoal at the feet and an iron rod.

The determined radiocarbon age of the burial is 610-680 cal AD (1377±24 BP, DeA-16588) (based on human bone).

mound 9

burial 1 (individual ID I16640, Male)

The determined radiocarbon age of the burial is 656-775 calCE (1320±20 BP, PSUAMS-10312) (based on human bone).

burial 6 (individual ID I19102, Male)

The burial of a 5-6-year-old child was in a rectangular pit excavated 0.2 meters into the subsoil and oriented east-west. A sparse stone rubblework layer was observed in the vicinity of the burial. The burial, likely oriented to the west, is heavily damaged. The skull and part of the tubular bones are concentrated in the western part of the pit. The burial inventory includes a bent bronze earring of Saltiv type, fragments of an iron knife, and a multi-component iron object of unclear purpose.

The determined radiocarbon age of the burial is 576-654 calAD (1440±30 BP, Poz-136244) (based on human bone).

mound 12, burial 1 (individual ID I19103, Female)

The outlines of the burial pit are not discernible. The buried individual, a child aged 6-7 years, was oriented with the head to the west, lying supine with arms extended along the torso. The spine is poorly preserved.

Two large buttons made of bronze and covered with gold foil were found under the mandibula. The buttons were filled with a white paste on the inner side, into which bronze loops were inserted for suspension. A glass bead with a bronze lining was found on the chest of the deceased. At the bottom, underneath the spine, there are several poorly preserved glass beads. No other items were present.

The determined radiocarbon age of the burial is 433-592 calAD (1545±30 BP, Poz-136245) (based on human bone).

#### *I.B.7. The Early Volga Bulgaria burials (MidVolga_EVB)*

##### I.B.7.1. Mullovka

##### I.B.7.2. Tankeyevka

See Szeifert et al., 2022 for the general description of the Tankeevka burial ground.

###### burial 961 (individual ID I25532, Female)

The determined radiocarbon age of the burial is 710-957 calAD (1185±30 BP, Poz-136155) (based on human bone).

###### burial 1139 (individual ID I25527, Male)

The determined radiocarbon age of the burial is 892-1023 calAD (1080±30 BP, Poz-136157) (based on human bone).

#### *I.B.8. The sites of the Karayakupovo horizon in the Lower Kama area (LowKama_KH)*

##### I.B.9.1. Bolshiye Tigany

See Szeifert et al. (2022) and Khalikova & Khalikov (2018) for the general description of the Bolshiye Tigany site.

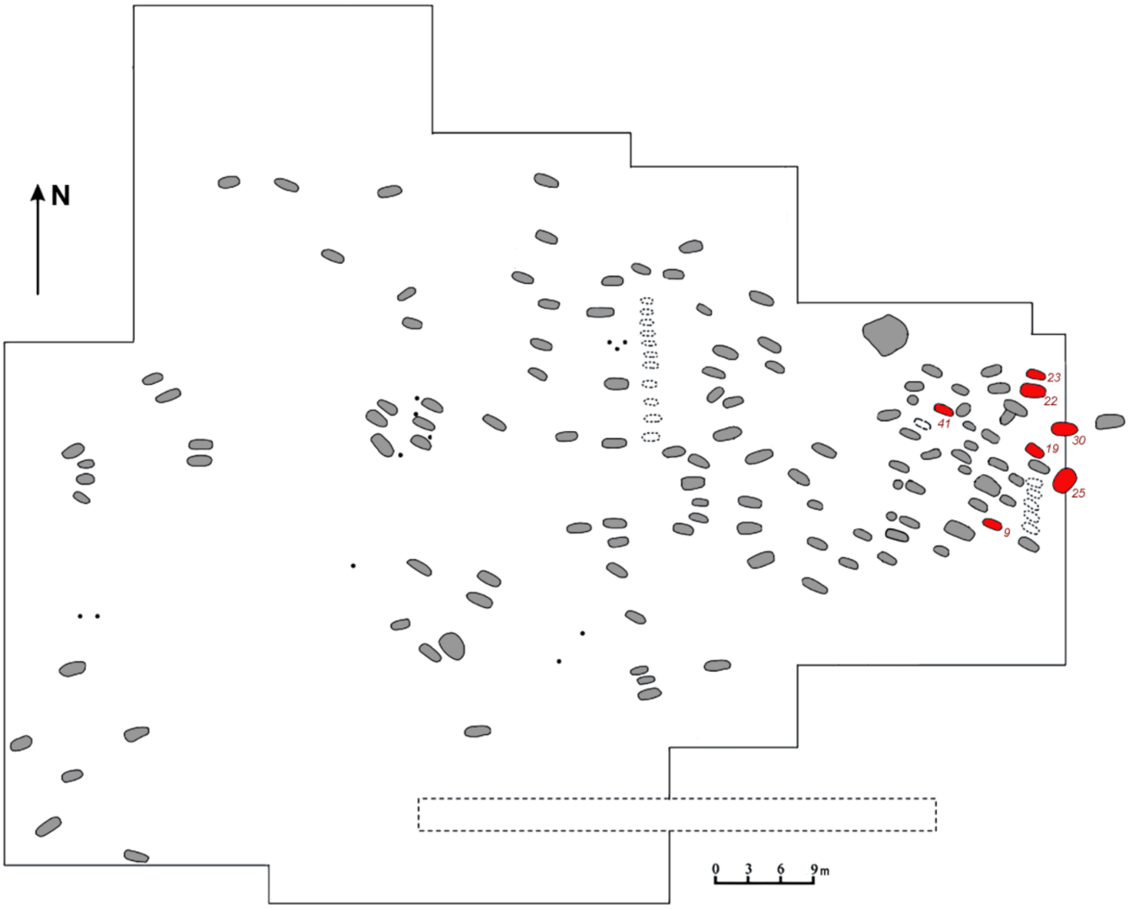

***Fig. SI.B.6.*** *A plan of excavated burials at Bolshiye Tigany.*

burial 9 (individual ID I25489, Male)

Grave pit (230×85 cm) was oriented along the west-south-west - east-north-east line. At the bottom, the poorly preserved skeleton of a male aged 30-40 years was found lying stretched out on his back, with his head to the west-south-west, faced upwards.

To the right of the skull, a horse femur (number 15 on the figure) and sheep tail vertebrae (16) were found. To the right of the mandibula of the deceased there was a small vessel (14) and a silver Saltiv-type earring (1). On the left side of the jaw, a leaf-shaped pendant (2) and a bronze bead (3) were discovered. On the elbow bones of the right arm, a bronze bracelet made of flattened wire (8) was found, and on the phalanx of the right middle finger, a silver ring with a pink stone in a bezel setting (9). Nearby, an iron knife (13) and an awl (12) were located. On the belt, remnants of a leather strap with two end straps decorated with silver fittings (10) were found. To the right of the belt, there was presumably a leather purse, from which a part of the round bronze lid with a punch ornament remained (11).

To the left of the skeleton, an iron saber, presumably placed in a wooden scabbard (6, 7), was found. The total length of the saber is 76 cm, with a slightly curved blade length of 67 cm. The sharp end of the blade, 11 cm long, is double-edged, while the rest is single-edged. The width of the blade is 3.0 cm. The crossguard has expansions at the ends. The handle is positioned at an angle towards the blade relative to the crossguard. The scabbard had a width of up to 4.5 cm. At a distance of 38 cm from the end, along the edge of the scabbard, there was an iron loop with a wooden attachment (7), and another one at the crossguard.

At the feet on the right side, the skull of a foal was uncovered. The bottom of the grave was lined with bast, and traces of fur and wooden coverings were preserved on the metal objects (Khalikova & Khalikov 2018).

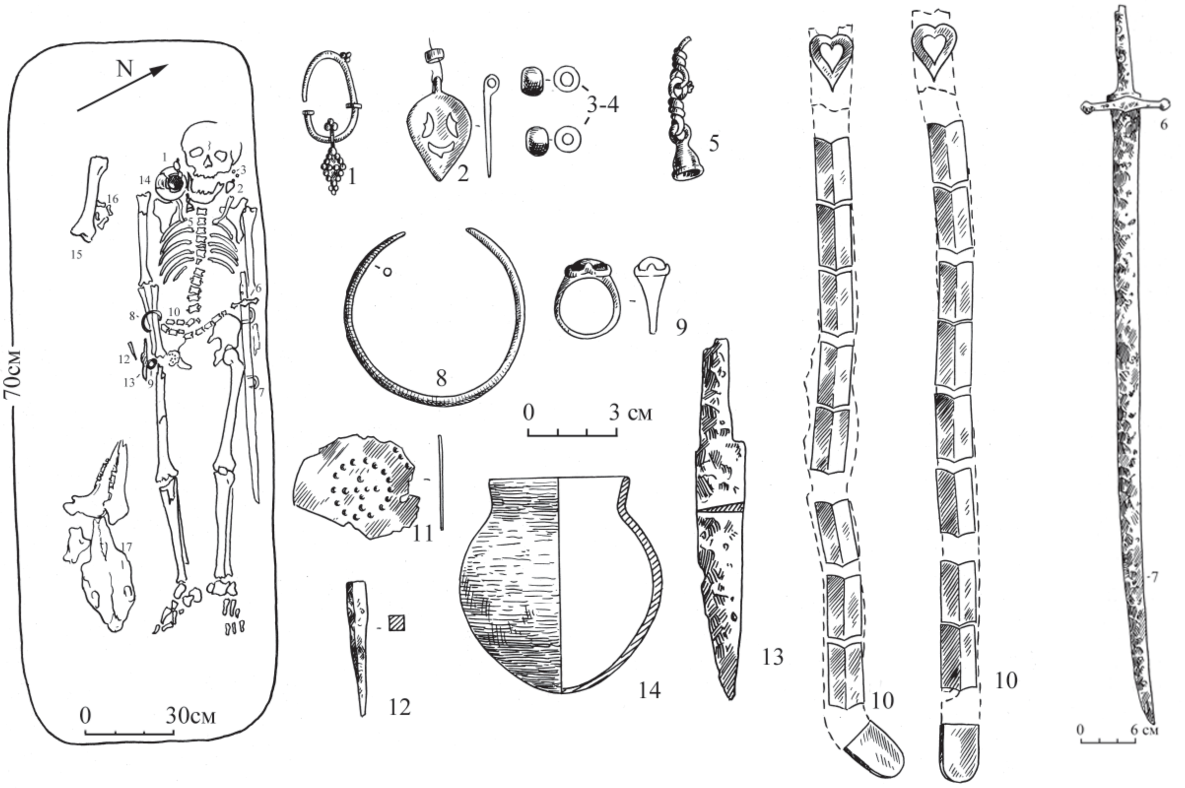

***Fig. SI.B.7.*** *Burial 9 at Bolshiye Tigany.*

burial 19 (individual ID I19105, Female)

The grave pit was initially traced at a depth of 70 cm as a sub-rectangular dark spot (80-100×215 cm) against subsoil loam. This spot was oriented from east to west and contained charred wood inclusions along the northern edge. At a depth of 77-80 cm, the pit narrowed on the western side, forming a ledge 25 cm wide. At the bottom of this ledge, a horse femur and small bones (36) were uncovered. At a depth of 80-90 cm, the main pit revealed: in the northern half, an “equine complex” consisting of a skull (35) and separately placed lower limbs (34) (horse was aged 5-6 years); in the southern half, a moderately preserved skeleton of a young female aged 16-17 years, lying on her back, head to the west, face to the south, with arms extended along the body. The deceased was apparently placed on a bedding of wood, felt, and fur. Carbonaceous inclusions were noted in the southeastern part of the grave (37).

The deceased was interred in relatively rich clothing, from which the following items were preserved: remnants of a headdress on the skull, with a forehead band adorned with leather and copper fittings (1-37 pieces), and a strap with silver buckles (3) and a tip (2) on the back of the head and crown; bronze earrings with metal beads ( 5, 6) and a simple gold ring (7) at the temples; remains of two necklaces on the neck – one with beads (white and blue glass – 37 pieces, 11), yellow beads (150 pieces, 11), multi-linked glass beads with silver lining (9 pieces, 10), bi- and tri-linked glass beads (8 pieces, 9), and five leaf-shaped bronze pendants (4 pierced - 13; 1 solid cast - 14), spiral-wound bronze beads (18), and three leaf-shaped pendants (15, 16, 17); nine cowrie shells on the left side of the chest (12), possibly adorning the collar or edge of the clothing; the back of the shirt collar was adorned with silver leaf-shaped appliques and pendants (22, 23); the clothing was belted with a leather strap with a silver buckle (26), two pierced fittings (27-28), and an end strap with a silver tip (29). The good preservation of the details allows for a proposed reconstruction of the woman's clothing based on the materials from this burial. Accompanying the deceased were: silver oxide powder from an applique in the left eye socket (4); a bronze bracelet with notches and bumps on the wire and bent ends (19) on the right hand; a pair of bronze rings with bezels on the fingers; two spherical bone beads (25) and an iron buckle (24) on the left hand; an iron knife pointing towards the feet (32) and a bronze sub-triangular ring (31) between the shins; fragments of an iron awl (30) and a round-bottomed cylindrical-necked vessel (33) near the left temple and back of the head (Khalikova & Khalikov 2018).

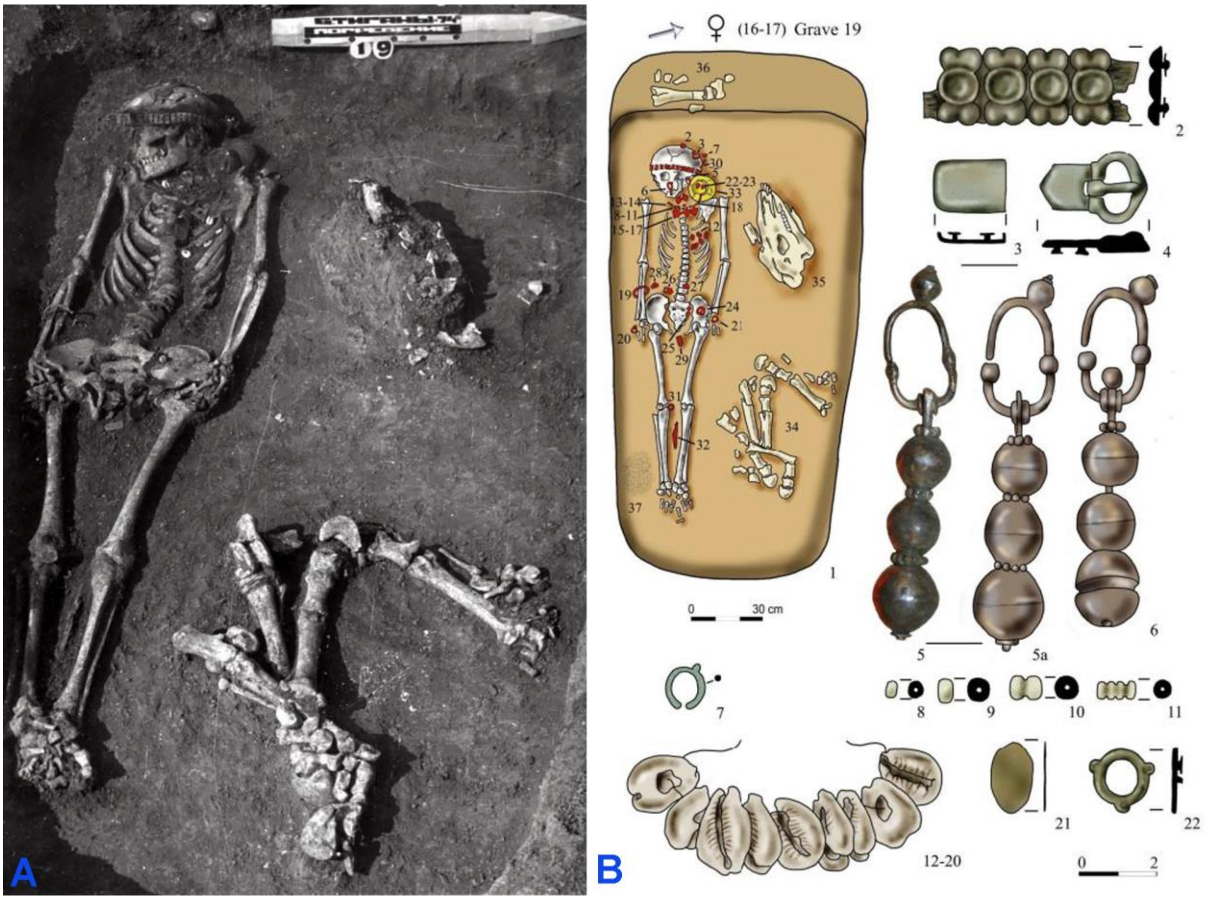

***Fig. SI.B.8.*** *Burial 19 at Bolshiye Tigany. A - photo, B - plan and grave goods.*

burial 22 (individual ID I19106, Male)

The grave pit was initially traced at a depth of 70 cm as a large dark spot (280×175 cm) oriented east to west. At a depth of 97-100 cm, the pit narrowed sharply towards the southwestern corner and continued as an oval spot (90×240 cm) down to a depth of 115 cm, leaving a large empty space in the rest of the pit (figure 26). The overall fill contained carbonaceous inclusions and a fragment of a handmade vessel (13). At the bottom of the main pit, at a depth of 105-115 cm, the moderately preserved skeleton of a male of 50-60 years old were uncovered, lying on his back, head to the west. Accompanying goods were: a horse femur at the head (1); a round-bottomed, thin-walled vessel with elegant carved Kushnarenkovo-type ornamentation under the right side of the skull (2); a saber in scabbard decorated with silver fittings along the left side (6); a bridle made of leather straps decorated with bronze fittings and connected by triplets was presumably thrown over the right knee and thigh (8) – its good preservation allows for a proposed reconstruction; two bone fittings from a bow were presumably placed on the left thigh (7), and a quiver with four iron socketed arrowheads of different types was placed on the right thigh (9). From the clothing and personal adornments of the deceased, the following items were preserved: a bronze Saltovo-type earring found near the right temple (14); remnants of a necklace with a large coin-like silver pendant (3) and a bronze bead (15); a bronze bracelet made of a sub-rectangular wire with bent ends on the right hand (10); a belt set consisting of six round silver fittings, six semicircular fittings with a ring, and three semicircular slotted fittings ( 4), and a bronze tip (5); an iron knife in the remnants of a leather sheath (11), and a pouch with a fire striker, from which a flint and rust from the steel were preserved, were presumably suspended on the right side of the belt (17).

Traces of fur and wooden bedding were observed under the metal items (Khalikova & Khalikov 2018).

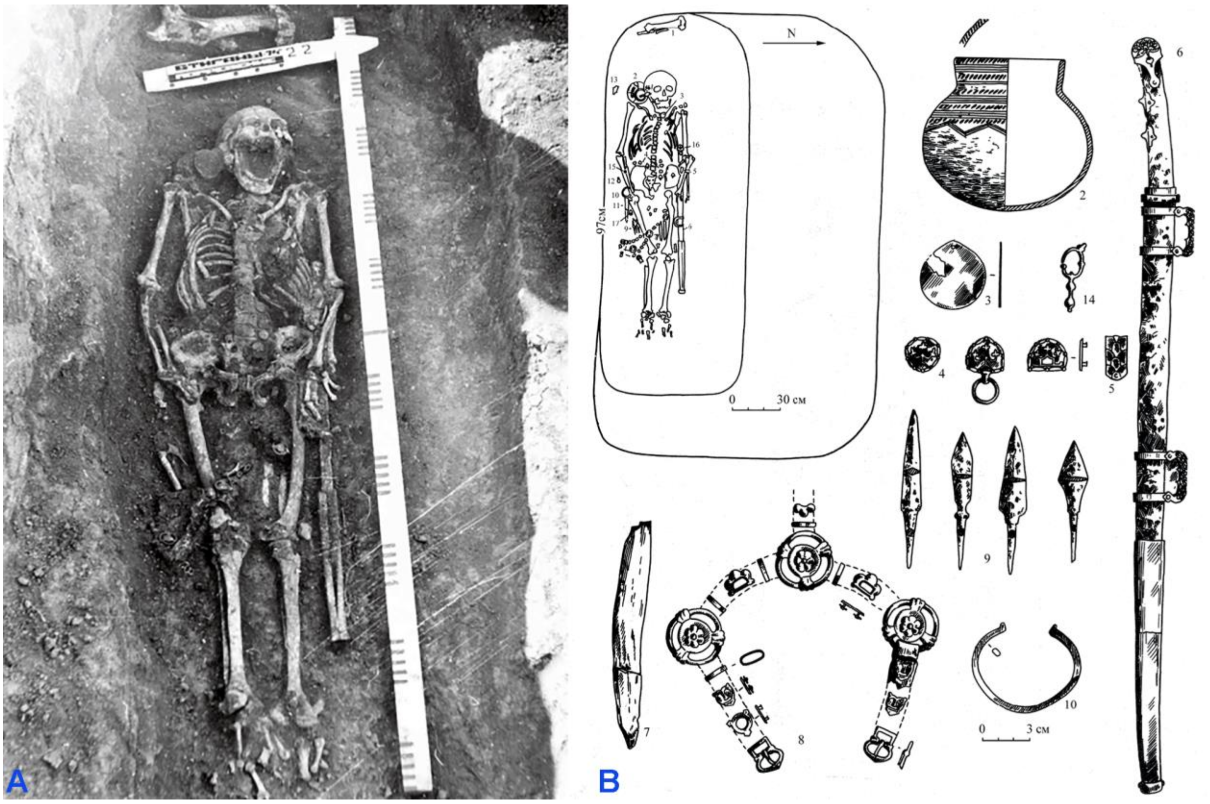

***Fig. SI.B.9.*** *Burial 22 at Bolshiye Tigany. A - photo, B - plan and grave goods.*

##### burial 23 (individual ID I19107, Male)

The grave pit was initially traced at a depth of 70 cm in the subsoil loam as a spot with rounded ends (65-80×200 cm) oriented east to west was outlined. At a depth of 90 cm, the pit revealed the moderately preserved remains of an elderly male (50-60 years old), laid on his back, head to the west, with the skull slightly turned to the right. The deceased was accompanied by grave goods. On the left side of the head, a cluster of horse bones (burial food?) (9) and a Kushnarenkovo-type clay vessel (11); on the right leg's shin bones, a horse skull (8); on the right forearm, remains of a bridle, from which several bronze fittings and a bit ring were preserved (2a-2c); along the right side of the upper torso, presumably a quiver, from which three iron socketed arrowheads (1) and an iron tip (10) were preserved. There was tarnish from silver eye covers on the skull's eye sockets (12). From personal adornments and clothing of the deceased, the following were preserved: a Saltiv-type earring near the right temple (13); around the neck area, three spherical bells possibly from clothing fasteners or a necklace (3); on the waist, remnants of a belt with a buckle (5), a semicircular fitting (4), and two sub-rectangular fittings (6), the surfaces of which were decorated with an original ornament depicting winged horses; on the index finger of the right hand, a ring with a stone (7); on the right side of the belt, an iron knife (15) and a fire striker, from which a flint and remains of a steel striker (14a-14c) were preserved (Khalikova & Khalikov 2018).

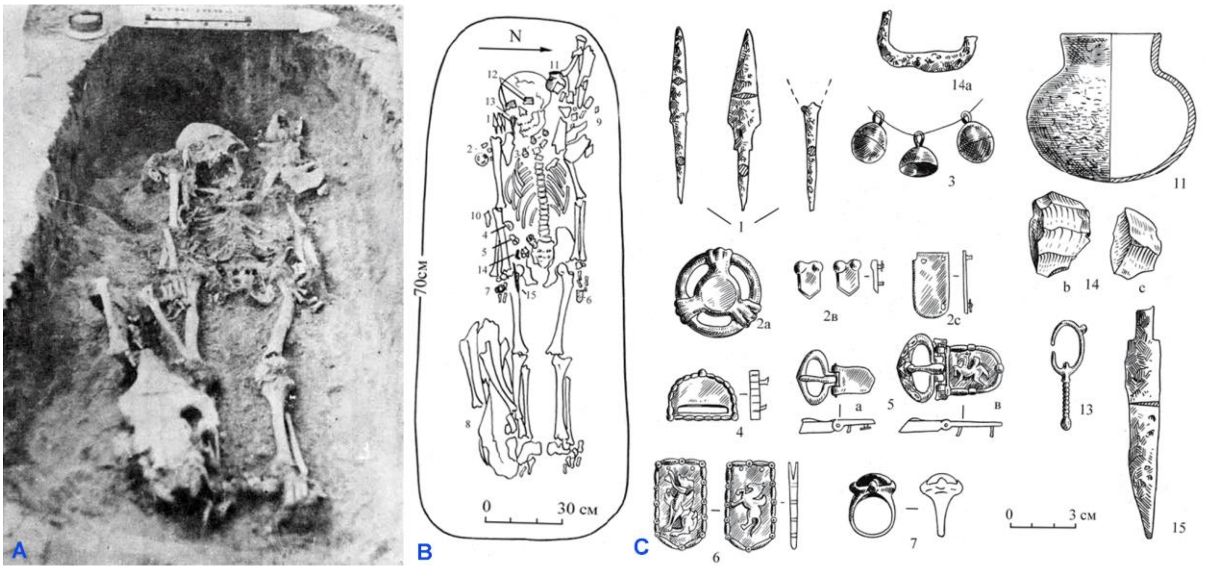

***Fig. SI.B.10.*** *Burial 23 at Bolshiye Tigany. A - photo, B - plan, C - grave goods.*

##### burial 25, individual B (individual ID I19108, Male)

The grave pit was initially traced at a depth of 65-70 cm as an oval-shaped spot (120×185 cm) oriented east to west.

Individual A. In the southern part of the spot, at a depth of 75 cm, the poorly preserved remains of an infant were uncovered, apparently laid on its back with the head to the west (only the decayed bones of the upper half of the torso were preserved). At the head, a large Kusnarenkovo-type clay vessel was found (1), along with beaver bone pendants (5). On the right side of the skull, there was a bronze Saltiv-type earring (2); around the neck area, remains of a necklace made of small glass beads with silver inserts and paste beads (3); on the waist, remnants of a belt with copper hemispherical fittings (3 pieces) and a trefoil fitting (4).

Individual B (individual ID I19108). The northern part was deepened by 20 cm, and at a depth of 90 cm, the relatively better-preserved remains of a 6-7-year-old boy were uncovered, also laid on his back with the head to the west (the skull was crushed and turned to the left temple). The deceased was accompanied by: at the head and above the skull, a fragment of a large horse bone (shoulder blade?) (1); in front of the skull, a crushed clay vessel (2). From the clothing and personal adornments of the deceased, the following were preserved: on the right side of the skull, a bronze Saltovo-type earring (3); under the mandibula, remains of a necklace with a silver coin-like pendant (6) and glass beads with silver inserts (4); on the neck, a bronze button with a convex head (5); on the lower chest, a leaf-shaped openwork pendant (7) and a bronze spherical button (8); on the right wrist, several dark blue paste and glass beads with silver inserts (9). An iron knife was found on the left side of the pelvic bone (10) (Khalikova & Khalikov 2018).

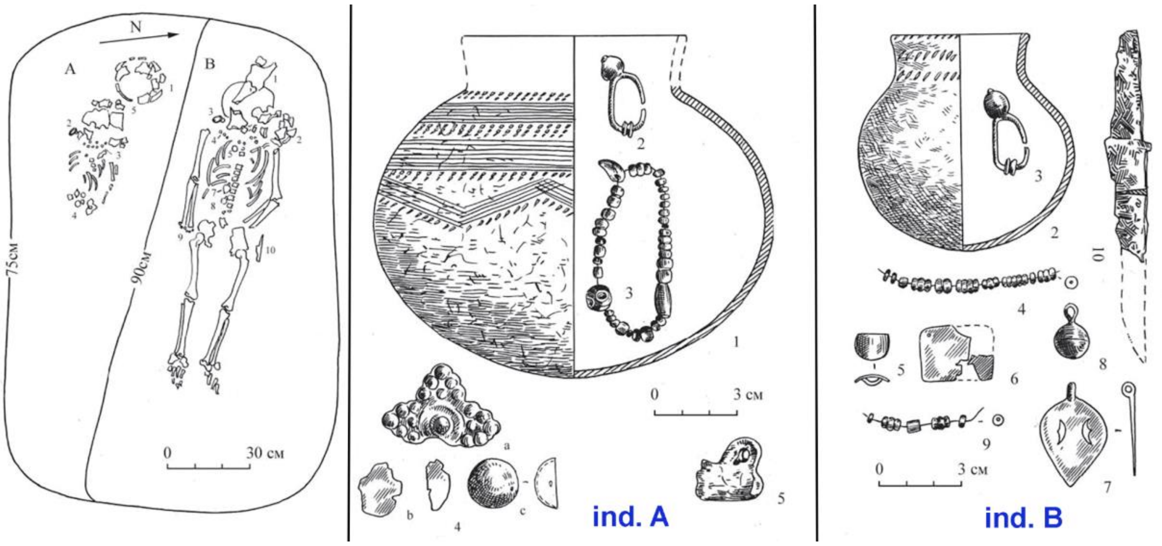

***Fig. SI.B.11.*** *Burial 25 at Bolshiye Tigany, grave goods from indivials A and B.*

burial 30 (individual ID I25490, Male)

The grave pit was initially traced at a with a depth of 80 cm, oriented fairly strictly from east to west. At the bottom of the grave pit, the remains of a female aged 35-45 were uncovered, laid on her back with her head to the west. A round-bottomed cylindrical-necked vessel was placed between the left side and the left elbow (9).

From the clothing and personal adornments of the deceased, the following were preserved: on the forehead and occipital parts of the skull, remnants of a headband made from appliqués attached to a leather strap with paired protrusions (1 - 27 pieces) and a bronze bead (16); under the right part of the occiput, a part of a bone comb with fine carved ornamentation was found (18); on both sides of the jaw lay massive earrings made of poor-quality silver with pin-shaped pendants (2); on the left eye socket, a silver eye guard fragment remained (4); under the jaw, in the upper part of the chest, remnants of a necklace made of beads and three bronze leaf-shaped openwork pendants (3, 12); in the upper part of the chest, an iron plate (fastener?; 5) and two small bronze spirals (7); below them, fragments of a bronze stick (6) and two bronze cast buttons (8); on the radial bones of the left hand, a bronze octagonal bracelet at the ends (10); on the middle finger of the right hand, a poor-quality silver ring with a dark stone in a bezel setting (11); in the upper part of the pelvis, remnants of metal belt details were preserved - a bronze buckle and bronze appliqués - four-part and heart-shaped openwork (12), a silver tip with an oval end (14) and two bell-shaped pendants (13); to the right of the belt, an iron knife was suspended, sheathed in a leather scabbard with a silver overlay (17), with a bronze bracket at the top of the scabbard (15).

Under the skeleton, especially in places where metal items were located, traces of bedding made from wood, felt, and fur were found (Khalikova & Khalikov 2018).

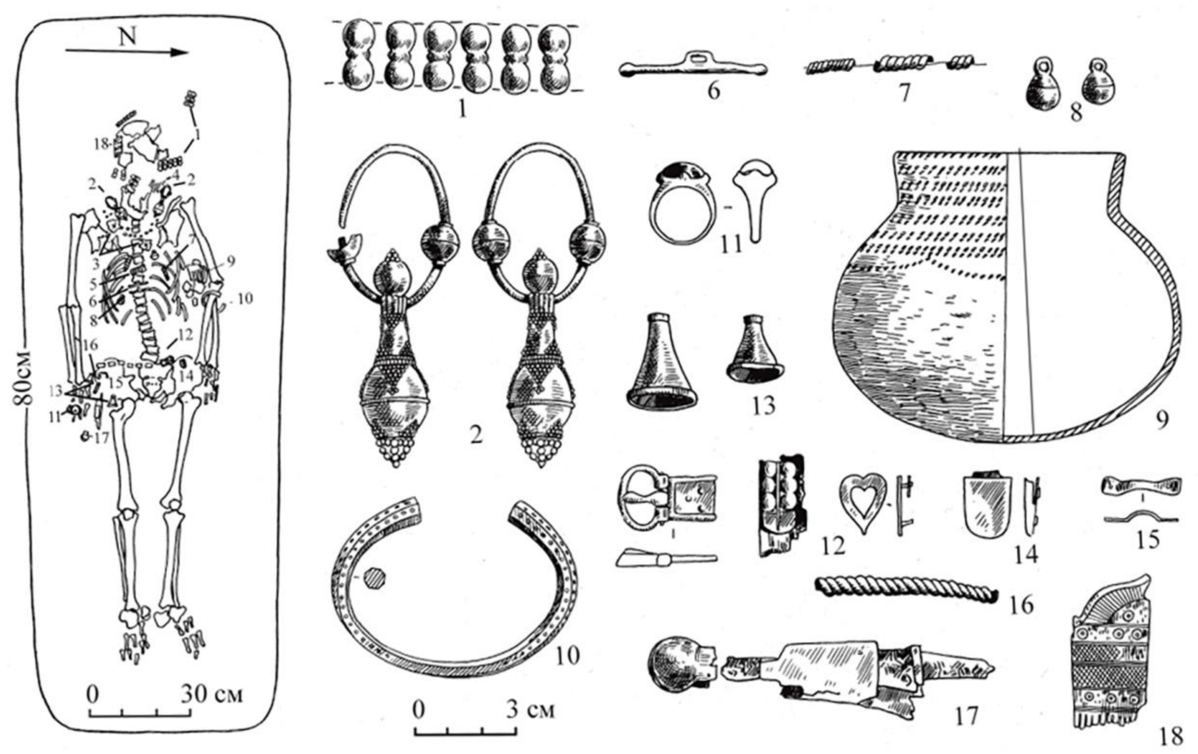

***Fig. SI.B.12.*** *Burial 30 at Bolshiye Tigany. A - photo, B - plan and grave goods.*

burial 41 (individual ID I19109, Male)

The grave pit had dimensions 230×85 cm. At a depth of 65 cm, the poorly preserved remains of a male individual (aged 45-55 years) were unearthed, positioned supine with the head extended towards the west. The skull is turned to the left temple, with the left leg bones showing signs of rodent damage.

Accompanying the interred were placed: along the left side of the torso - a quiver, from which a set of six iron tang arrowheads (8) and an iron loop (11), likely from the binding of the upper part of the quiver, were preserved; a bow (bone overlays preserved - 9) and possibly a bridle with two bronze bit rings (10); at the right ulna - an iron knife (5).

Fragments of a silver plate from an eyeshield (3) were discovered around the left orbit.

From the details of the clothing and personal adornments of the deceased, the following were preserved: on the right temple - a bronze ring-shaped earring (4); around the neck area - remnants of a necklace made of glass beads (1) and fragments of two copper rattles (2); on the wrist of the right hand - a bronze bracelet with flattened and curved ends (6); on the right side of the pelvis - a bronze buckle (7).

Metal objects preserved traces and fragments of wood on top, likely from the lid of the burial or the wooden covering of the grave.

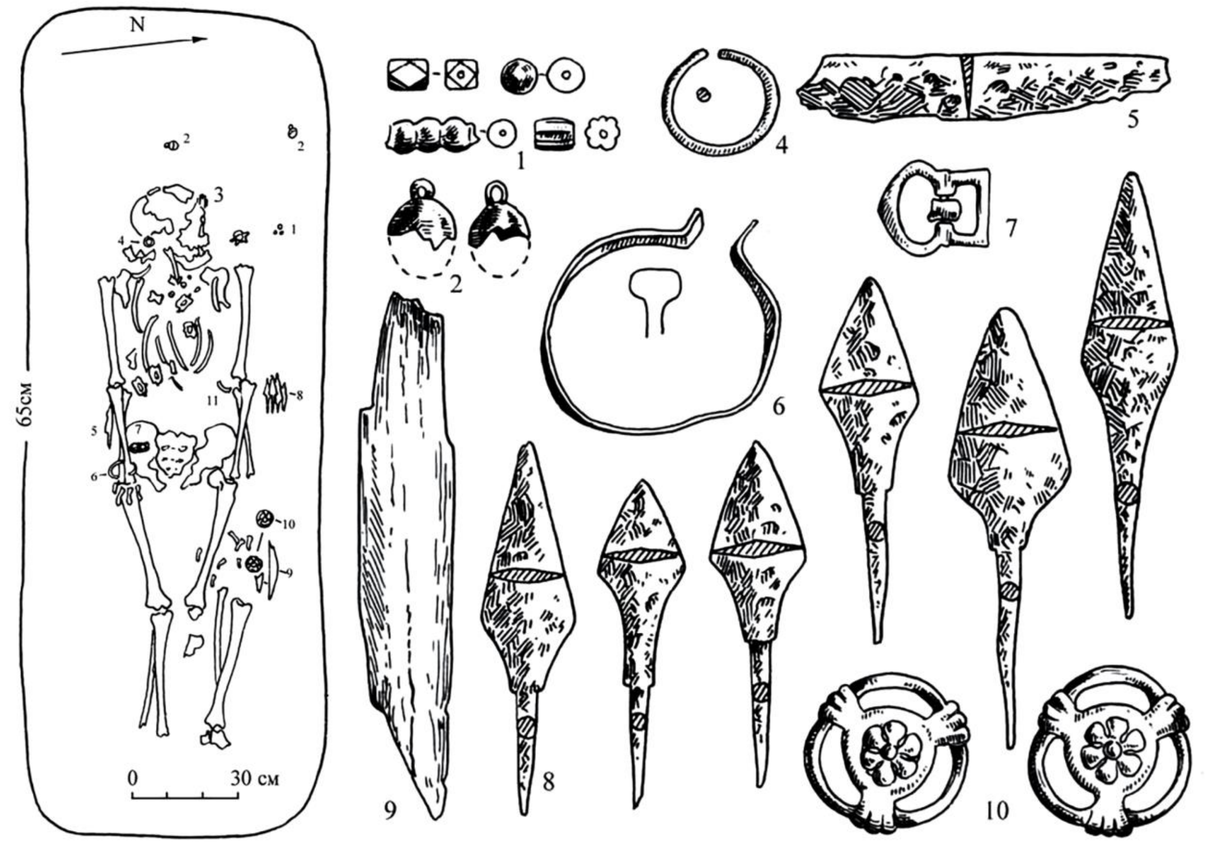

***Fig. SI.B.13.*** *Burial 41 at Bolshiye Tigany.*

#### I.B.10. The Medieval Chiyalik culture in the Lower Kama area (LowKama_Chiyalik)

##### I.B.10.1. Gulyukovo

The archaeological site is located at the northern tip of the watershed between the Igat and Tirgaush rivers, left tributaries of the Ik River, where they flow into the Nizhnekamsk Reservoir (formerly, before the reservoir's formation, where they entered the low floodplain of the Ik River). The relief of the watershed from the village of Gulyukovo to the reservoir consists of very sloping ridge with elevations ranging from 85 to 70 meters above sea level. The surface features gentle slopes (10-12°) sloping towards both the reservoir and the valleys of the side streams. Towards the reservoir, the surface gradually levels off (down to 3°), and as it approaches the edge, within a strip 10-20 meters wide, it becomes almost horizontal (1-3°). However, in cross-section, the final segment retains a gently convex profile, visually well observed from the reservoir shoreline and recorded by height measurements of the erosion ledge: it varies from 5-6 meters in the area of excavations 4 and 5 in 2009 (at the center of the terraced slope) to 1 meter near the modern channels of the Igat and Tirgaush rivers (on the flanks) (Khisyametdinova, 2012).

The opposite banks of the Igat and Tirgaush rivers, facing the watershed, consist of the ridge tips extending to the Ik River with elevations of 100-150 meters and steep slopes complicated by gullies and ravines. As a result, the Igat-Tirgaush interfluve represents the most convenient (and perhaps the only sufficiently wide and low) passage into the Ik-Kama floodplain from the village of Kulushevo (Tukaevsky district) to the town of Menzelinsk.

The occupation site at Gulyukovo attributed to the Neolithic and Late Bronze Age are known since 1979. The burials eroded by the abrasion of the Nizhnekamsk Reservoir were first documented in the Gulyukovo area by Andrey Chizhevsky in 1995-1996, who excavated 8 of them and called the site the Guluykovo Cemetery. From 1997 to 2001, 2003-2004, and 2007, N.M. Kaplenko, a teacher from Secondary School No. 8 in Naberezhnye Chelny, excavated approximately 538 m2, investigating 56 burials identified as "early Muslim" and attributed to the period from the 12th to the early 14th centuries CE (4 of them belong to the late Sarmatian part of the cemetery and are dated to the mid-second half of the 4th century CE). In 2006, Dmitry Bugrov and Ayrat Sitdikov excavated approximately 293 m2, studying 43 burials (7 identified as late Sarmatian, 36 as "early Muslim" from the second half of the 11th to the beginning (first third?) of the 13th centuries CE. In 2007 Dmitry Bugrov excavated another 103.5 m2 of the site area, revealing 15 more "early Muslim" burials. Further rescue excavations covering an area of approximately 225 m2 were conducted by Ruslan Matveyev in 2009, Anton Lyganov and Renat Valiev in 2012, Pavel Krasilnikov in 2014 under the supervision of Dmitry Bugrov. These excavations did not yield any medieval burials; instead, layers of the settlement, 6 Late Sarmatian burials, and a Medieval household were discovered. In total, at the moment approximately 1160 m2 area are unearthed, revealing 111 burials attributed to the Chiyalik culture (see Bugrov, Asylgaraeva, 2020 for detailed information and references).

The total area of the Medieval cemetery at Gulyukovo reaches 2 hectares. The burials were arranged in more or less dense groups within the excavation area, organized into 2-4 vague rows, with rows stretched parallel to the edge of the terrace (along the SSE-NNW and SSW-NNE lines), with gaps between groups ranging from 3-4 to 10-15 meters.

The burial rite of the site is characterized by individually arranged inhumations in simple graves (elongated rectangular in shape with rounded corners or ends), approximately corresponding to or slightly exceeding the body size. In 8 cases (burials 5, 7, 8, 19, 20, 39, 44, 47), a "leaning" of the northern long wall of the pits was observed (similar to the "lahd" feature). The buried individuals were laid out in the supine positions stretched on their backs, with their heads mostly oriented towards the southwest and south-southwest (36 and 32 cases, respectively), followed by west (12 cases), south-southwest (11 cases), and one case each towards the south, northwest, and west-northwest. In 42 cases, a pronounced turning of the head (less frequently the torso) of the deceased was observed: in 30 burials to the right, facing south and southwest (towards Mecca), and in 12 cases to the left, facing south and south-southwest and facing north and north-north-west.

In three cases (burials 49, 51, 54), a specific disturbance of the anatomical order of the skeletons was observed: some bones, primarily limbs, retained their position, while the bones of the chest, spine, pelvis, and less frequently the skull and part of the limb bones, were mixed and lay in disorder, but approximately in their anatomical positions (possibly reflecting a "neutralizing the deceased" ritual).

Inventory items were found in 16 burials, including iron coffin nails in one burial (burial 31K). In other cases, the grave goods are represented by decorations, clothing accessories, and tools: burial 1K - bracelet, round pendant-button, beads; burial 2K - iron knife; burial 8K - bone plate, iron knife, and two poorly preserved arrowheads (?); burial 11K - pendant-amulet, bracelet; burials 15K and 19K - one belt (?) plate in each; burials 18K and 12 - one each of bronze temple rings; burial 21K - a pair of similar rings and a knife; burial 2K - bronze ring, glass bead; PB (burial in the bank exposure) 3K - bronze ring, glass bead, and a knife; burial 4 - metal plate, possibly a pin (?), burin (?); burial 25 - fragment of a bronze ornament with grain, copper plate; burial 38 - knife, glass bead, a broken object made of lead-tin alloy; burial 51 - plate-like pendant made of copper. Items from destroyed burials were also present in the excavation material: temple rings made of "white bronze" and gold, a bronze drop-shaped faceted pendant-button, a bone needle, and iron knives.

The reference assemblages for the chronological interpretation of the site were found in burials 1K, 11K, and 4 (Bugrov et al., 2010, fig. 2).

In burial 1K, the earliest chronologically indicating item is a glass bead in the form of a "lemon" type (Bugrov et al., 2010, fig. 2: 6). Similar beads made of yellow semi-transparent glass with protrusions around the holes are characteristic of the second half of the 10th to the early 11th century in Eastern Europe, including Volga Bulgaria, persisting until the end of the 11th century. Individual finds in layers from the 12th century are considered residual. Spherical beads made of rock crystal and bipyramidal beads made of carnelian from burial 1K have their peak distribution in the Volga-Kama region in the 11th-13th centuries.

Round pendant-buttons from burial 1K and drop-shaped faceted "buttons" from the excavation material were common from the late 10th to the 11th century and were found in Bulgarian sites and the periphery of Volga Bulgaria, including the Mari Volga region, Permian Cis-Urals, Trans-Urals, and Western Siberia.

Bracelets with expanded rounded ends from burials 1K and 11K are represented in sites dated from the 11th-12th centuries in the Southern Urals, Upper Kama region, and the Vyatka basin.

The pendant-amulet from burial 11K dates within the 11th-12th centuries.

The bronze plate with a wide plate loop for suspension from burial 4 is similar to silver plates from the Butaikha hoard, interpreted as decorations for horse trappings and dated from the 11th to the first half of the 13th century. In the context of the Chally Ensemble, such plates serve as female necklaces, and the complexes with them are dated to the turn of the 11th-12th centuries to the first third of the 13th century or the 12th-13th centuries. Such decorations are one of the culturally defining features in the women's costume of the Chiyalik culture. Similarly, as the central medallion of a necklace or chest pendant, bronze and silver plaques of various types were used by the population who left the burial grounds of Selyanino Ozero, Kishert, Kushulevo, Azmetyevo-1, Taktalachuk, Derbyoshki in the Cis-Urals (Pastushenko, 2005, p. 37-40), Pylayevo (Pastushenko, 2005, p. 39) and “Kozyr” (Garustovich et al., 2008, p. 57) in the Trans-Urals. Some of them are extremely similar to the specimen from burial 4 in size and design (Pastushenko, 2006, Fig. 2: 2; 9: 6; 14: 15; Yutina, 2007, Fig. 7: 1, 8; Garustovich et al., 2008, Fig. 2: 23).

Thus, the overall dating of the site falls within the period from the second half of the 11th century to the beginning (possibly the first third?) of the 13th century. Narrower chronological phases of the site have not been delineated due to insufficient evidence. Presumably, the earlier part (associated with "inventory" burials) may have been obliterated by abrasion, while the later part (lacking inventory) does not have indicators for precise dating.

Specific elements of material culture left by the population buried at Gulyukovo include round bronze plaque with a plate loop, used as a neck-chest pendant (burial 4). Such ornaments are one of the culture-defining traits in the female costume of the Chyalyk culture. Similarly, bronze and silver plaques of various types were used by the population of other burial grounds in the Urals and Trans-Urals as central medallions of necklaces or chest pendants. Some of them closely resemble the specimen from burial 4 in size and construction.

Cultural influences on the population that left the Gulyukovo burial ground are traceable through characteristic elements identified in burial 31K. In this burial, the deceased was placed in a coffin secured with 11 iron nails, which is typical for Muslim cemeteries in the central regions of the Volga Bulgaria during the pre-Mongol period (11th – first third of the 13th centuries) and is not characteristic of the Chyalyk culture.

Taken together, based on the combination of burial ritual features (overall Muslim tradition with some manifestations of paganism - deviations in orientation and position of the buried, presence of inventory, post-burial manipulations with the skeleton), as well as the composition and chronology of the artifact complex, the monument can be attributed to the early stage of the Chiyalik culture.

##### burial 5 (individual ID I19078, Female)

The burial was excavated in 2006. The grave has an elongated sub-rectangular shape with rounded corners, oriented towards the south-southwest (azimuth 216°), and measures 185x50 cm at the fixation level. The filling consists of dense gray loam with traces of reddish-brown clay. The walls are sheer, with the northwest (long) side slightly overhanging at the upper part, while the southwest side is disturbed by a rodent burrow. The depth of the pit from the fixation level is 58 cm, and the bottom is flat.

The skeleton of a female, approximately 25 years old, as anthropologically identified by Ilgizar Gazimzyanov, and poorly preserved, was positioned on her back with the head facing southwest, and the arms extended along the body. The remains of the right forearm are placed on the right pelvic bone. The skull lies with the cranial part upwards, while the mandibula is displaced to the right, resting on the shoulder joint. The legs are slightly bent at the knees and turned to the left.

In the fill of the burial pit was found a sherd of a Late Bronze Age pottery vessel, originating from the cultural layer of the settlement.

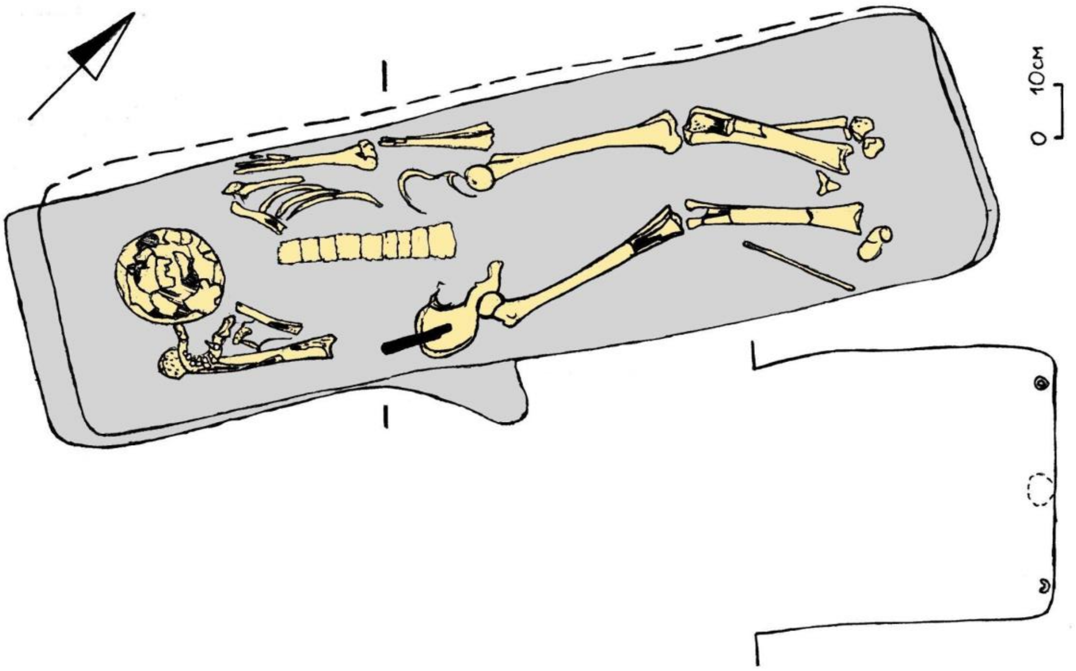

***Fig. SI.B.13.*** *Burial 5 at the Gulyukovo cemetery.*

##### burial 17 (individual ID I34074, Male)

The burial was excavated in 2006. The grave pit is elongated in shape, with a rounded southwestern end and an irregular northeastern end, oriented southwestward (azimuth 230°). The dimensions of the pit at the fixation level are 200 x 70 cm. The fill consists of heterogeneous soil (dense gray loam with traces of reddish-brown clay). The walls are sloping, and the depth of the pit from the fixation level is 25 cm, with a concave bottom.

The skeleton was positioned elongated on its back with the head facing southwest; the arms were stretched alongside the body, with the radial bones passing under the edge of the pelvic bones. The skull lies on the occiput facing upwards, slightly tilted to the right (southeast); the feet are turned outwards.

The buried person is male, aged 35-45 years, as anthropologically identified by Ilgizar Gazimzyanov. The right ulna is fractured, with non-union of the fracture (the ends of the fracture overlap, a large bony callus "false joint" formed on the upper fragment of the ulna, and a bony outgrowth formed opposite the fracture on the radius). The vertebrae are deformed (flattened), and the vertebral body edges are sharpened; there are traces of fused fractures on the right ribs.

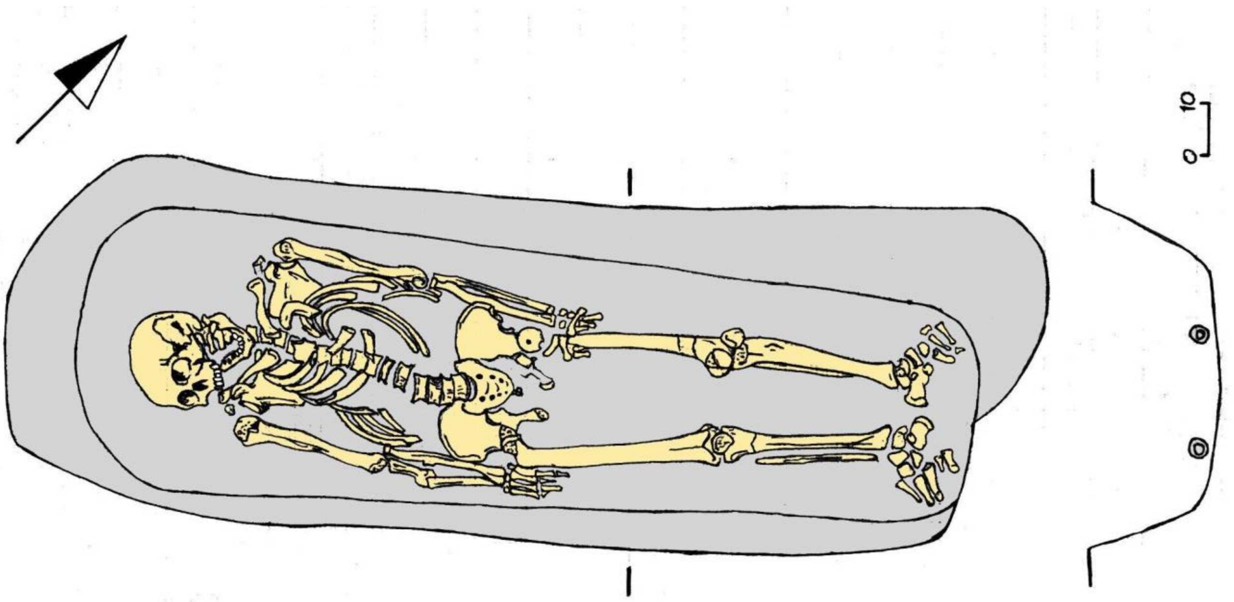

***Fig. SI.B.14.*** *Burial 17 at the Gulyukovo cemetery.*

##### burial 20 (individual ID I19079, Male)

The burial was excavated in 2006. The grave pit is elongated in shape with rounded corners, oriented southwestward (azimuth 235°), measuring 225 x 56 cm at the fixation level. The fill consists of heterogeneous soil (dense gray loam with traces of reddish-brown clay). The walls are steep, with the northwest wall overhanging; the eastern corner of the grave pit up to a depth of -152 cm is damaged by post hole 11. The depth of the pit is 48 cm.

The skeleton of a male, aged 20-30 years, as determined by Ilgizar Gazimzyanov, was positioned elongated on its back with the head facing southwest with a slight turn to the right (southeast); the arms were stretched alongside the body, with the left forearm passing under the edge of the pelvic bone. The right leg is slightly bent at the knee. The skull lies on its right side, facing southeast.

In the fill of the grave pit a fragment of a vessel rim of unclear cultural and chronological affiliation (Late Sarmatian period?; Middle Ages?) was found, with the inclusion of coarse grog in the molding mass.

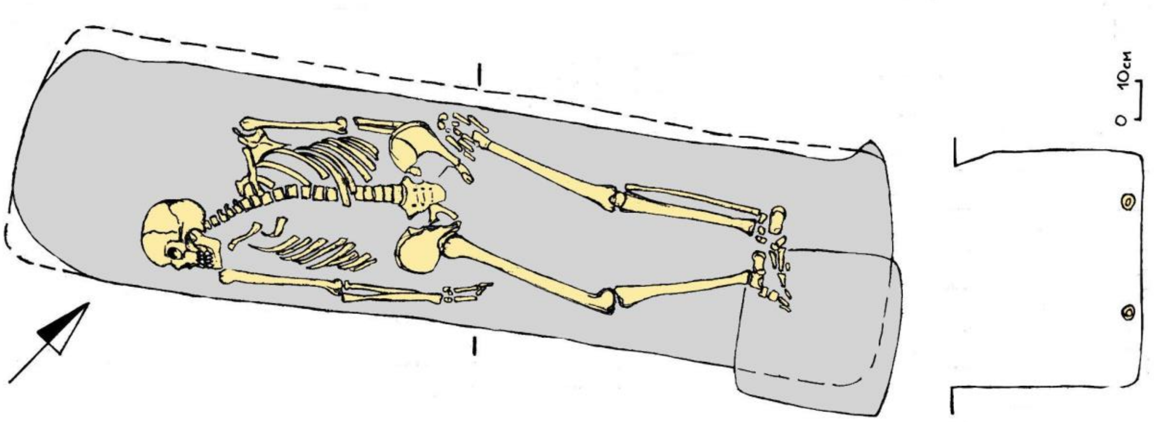

*Fig. SI.B.15.* Burial 20 at the Gulyukovo cemetery.*

##### burial 31 (individual ID I25503, Male)

The burial was excavated in 2006. The grave pit is elongated in shape with rounded ends, oriented southwestward (azimuth 233°), measuring 195 x 50 cm at the fixation level. The fill consists of heterogeneous soil (dense gray loam with traces of reddish-brown clay). The walls are steep, and the depth of the pit from the fixation level is 12 cm, with a flat bottom.

The skeleton of a male (?), aged 16-20 years, as determined by Ilgizar Gazimzyanov, was positioned elongated on its back with the head facing southwest; the arms were pressed against the body (the deceased may have been wrapped?), with the left shoulder bone lying on the ribs, and the hands placed on the pelvis (the bones of the left forearm are displaced and lie on the sacrum and the right side of the pelvis, to the right of the spine). The skull lies on its right side, facing south. The feet are turned to the right (southeast).

In the fill of the grave pit, fragments of a vessel of unclear cultural and chronological affiliation (Late Sarmatian period?; Middle Ages?) were found, with the inclusion of coarse grog and sand in the molding mass (possibly natural impurities in the clay): 2 fragments of a flat slightly concave bottom and 2 fragments of walls.

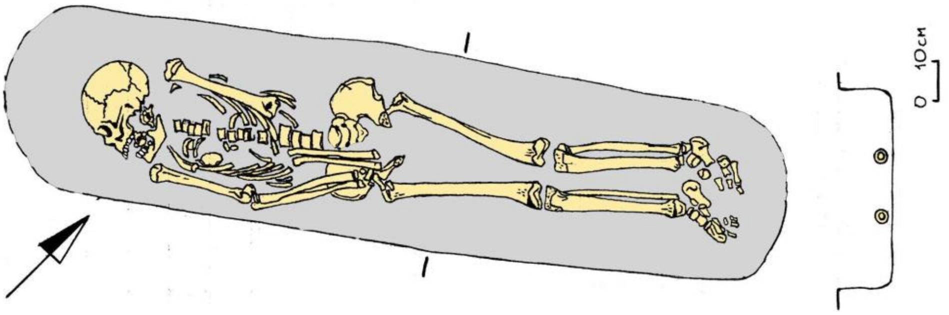

***Fig. SI.B.16.*** *Burial 31 at the Gulyukovo cemetery.*

##### burial 51 (individual ID I19080, Female)

The burial was excavated in 2007. The grave pit is elongated with a nearly rectangular shape and rounded corners in the western part, and a rounded eastern end, oriented southwestward (azimuth 227°), measuring 220 x 55 cm. The fill consists of heterogeneous soil (dense gray loam and reddish-brown clay in approximately equal proportions). The walls are slightly sloping, almost vertical, with a depth of 30 cm from the fixation level and a flat bottom.

The skeleton of a female, aged 30-40 years, as determined by Ilgizar Gazimzyanov, was positioned elongated on its back with the head facing southwest, and the anatomical order is partially disrupted. The preserved bones include the shoulder blades, right ulna, right pelvic bone, and leg bones (except for the left femur); the feet are turned to the right. Partially preserved are the original positions of the left femur (rotated 90° around its longitudinal axis, with the greater trochanter upward and the head of the femur downward) and the hands (located on the outer side of the pelvic bones). Based on the position and orientation of the preserved in situ long bones, the arms of the deceased were likely extended along the body. The skull is inverted, lying on the right temporal bone with the facial aspect facing southwest; other bones such as the mandibula, ribs, sternum, scapulae, clavicles, and vertebrae (including the sacrum) are scattered in the chest area of the deceased; the left pelvic bone is rotated with the iliac crest upward. Overall, the displaced bones are close to their original positions. It is difficult to determine whether the disruption of the anatomical order of the skeleton was caused by animal burrowing activity or intentional human actions (robbery? ritual post-mortem treatment?) with certainty.

In the western part of the grave pit, in front of the facial aspect of the skull, there was a narrow, thin copper plate (unfolded rim of a copper vessel?) with a elongated-triangular hole at one end (a pendant?) (preserved in three fragments); within the fill of the grave pit, without a precise location, there is a small flint knife-like plate (a knife with two blades for cutting soft material and a point as identified by Madina Galimova), originating from the cultural layer of the settlement.

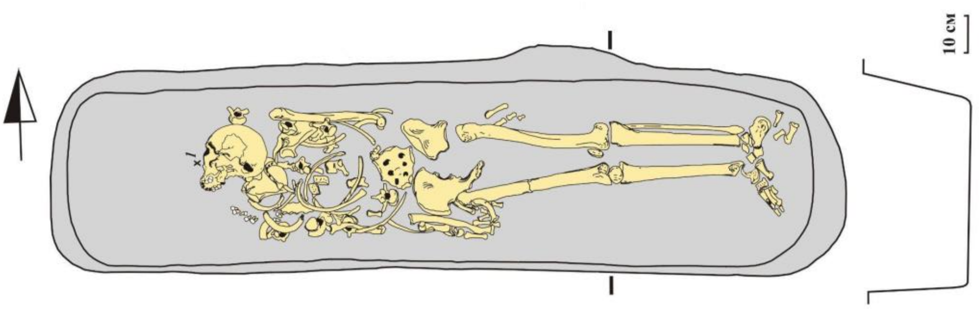

***Fig. SI.B.17.*** *Burial 51 at the Gulyukovo cemetery.*

##### burial 60 (individual ID I19081, Male)

The burial was excavated in 2007. The grave pit has an elongated shape, oriented southwestward (azimuth 250°), with the eastern end destroyed by an edge, and the western end irregularly rounded. The preserved portion measures 195 x 58 cm at the fixation level, while at -220 cm depth, the length increases to 210 cm due to the slope of the edge that destroyed the eastern end of the pit. The fill is heterogeneous: dense dark-gray loam with occasional inclusions of reddish-brown clay (across the entire area of the pit at the primary fixation level, and in the middle part of the pit and below) and variegated fill (dense gray loam and reddish-brown clay in approximately equal proportion) (along the walls and at the bottom, at the contouring level and below). The walls are slightly sloping, almost vertical, and uneven in the western (end) part; the depth from the fixation level ranges from 51 to 54 cm, with a flat bottom.

The skeleton of a male, aged 25-35 years (?), as determined by Ilgizar Gazimzyanov, was positioned elongated on its back with the head facing southwest, and the arms extended along the body, with the left forearm lying on the left hip joint. The skull is lying on the right side, facing south. The hands are missing; the foot bones (all phalanges, part of the metatarsals, and cuneiform bones) are displaced and unevenly distributed in the fill, mainly in the eastern half of the pit: some phalanges and metatarsals were found in the upper fill layer; the majority of the phalanges and metatarsals formed a cluster at the bottom of the eastern end of the pit, primarily between the right shinbone and the southeast wall of the grave; one cuneiform bone

was found at the western end of the grave, behind the skull. This is likely the result of animal burrowing activity, although the outlines of burrows in the fill are not evident.
In the upper fill layer at the western end of the pit, a flint chip without traces of use was found. Additionally, from the fill without a precise location, fragments of flint plates (chisels for wood, cutters, and knives) were recovered, as identified by Madina Galimova, as well as small fragments of ceramic vessel walls from the Volga-Kama Neolithic culture and the Late Bronze Age (?); all findings originate from the cultural layer of the settlement.

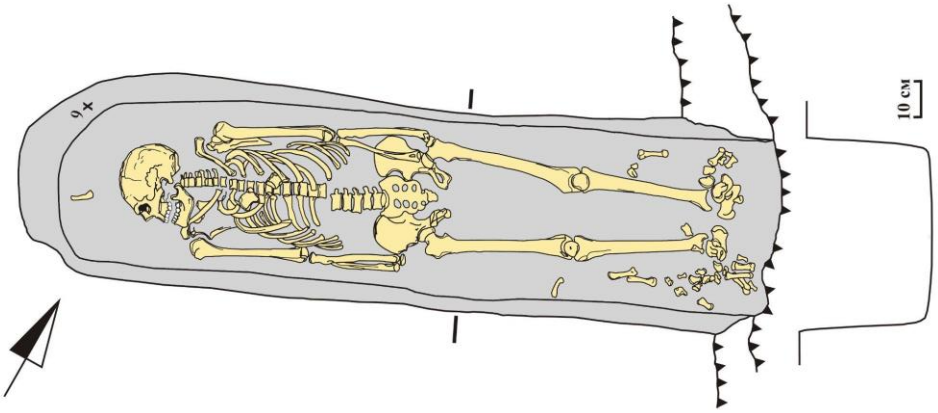

*Fig. SI.B.18.* *Burial 60 at the Gulyukovo cemetery.*

##### burial 63 (individual ID I33730, Female)

The burial was excavated in 2007. The contours of the bottom part of the grave pit, elongated with a rounded end, oriented southwestward (azimuth 247°), were traced at a depth of -182 cm; the majority of the burial remains outside the excavation area, with the portion that entered the excavation measuring 80 x 30 cm at the fixation level. The fill is variegated (dense gray loam with inclusions of reddish-brown clay). The walls were not identified during clearing; the profile of the southeast wall of the excavation reveals overhanging of the end wall of the burial pit. Judging by the profile, the depth of the pit from the surface level is 40 cm, with a flat bottom slightly raised towards the walls.
The skeleton of a child, approximately 5 years old, as determined by Ilgizar Gazimzyanov, partially within the excavation area (upper half down to the thoracic region of the spine), was positioned elongated (?) on its back with the head facing southwest; the left arm is extended along the body. The skull is lying on the right side, facing southeast.

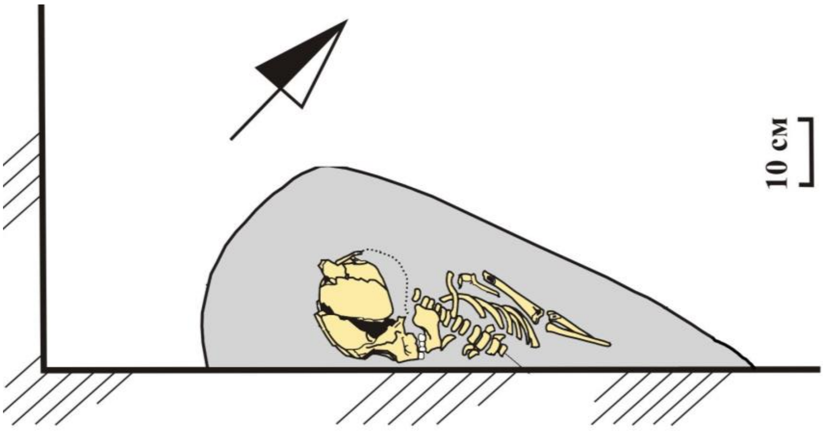

*Fig. SI.B.19.* *Burial 63 at the Gulyukovo cemetery.*

##### I.B.10.2. Ust-Menzel'ya

The hillfort at the Ust-Menzelya locality is known since 1929. The cultural layers and finds from the enclosure determine the Early Iron Age (Pyany Bor culture) as the main period of inhabitance, with layers from the Eneolithic period (Novoilyinka and Garino cultures); sporadic finds include ceramics from the Neolithic period (Kama culture) and from the end of the Bronze Age to the beginning of the Early Iron Age (Lugovoy culture, Textile Ware, or Ananyino culture). The burials in the fortified area were exposed by ravine erosion and documented in 1998 by L.V. Mil’chakov in 1998 (hunting inspector for the Menzelinsk and Aktanysh districts of the Republic of Tatarstan; one partially destroyed non-inventoried burial was inspected). In 2001, during a survey by Dmitry Bugrov, in a control trench approximately 1.5 m2 of heavily damaged child burial (3-4 years old, with undetermined cultural-chronological position) was unearthed. In 2007 Dmitry Bugrov and Madina Galimova carried out excavations of a total area of 183.5 m2, which yielded five burials identified as "early Muslim" and attributed to the early phase of the Chiyalik culture.

###### burial 5 (individual ID I33733, Male)

The outline of the burial pit was not traced; the skeleton of a child aged 8-10 years old as determined by Ilgizar Gazimzyanov, was found in a layer of yellow clay with admixtures of lime crumbs and with streaks and layers of gray-brown clay. It was oriented with the head to the southwest; the legs were extended, the right foot was turned outward, the left was extended; the torso from the pelvic bones and above was turned to the right side, the left arm was slightly bent at the elbow, lying over the body with the hand on the pubic bones, the right arm was extended along the body, the humerus with the proximal end passed under the right ribs, the forearm bones under the outer edge of the right ilium, the phalanges and metacarpals under the greater trochanter of the right femur. The skull lies on the right side, with the facial part to the south.

A fragment of an unornamented wall of a ceramic vessel with admixtures of crushed shell in the molding mass (Pyany Bor culture) was found at the level of excavation; it originates from the cultural layer of the hillfort.

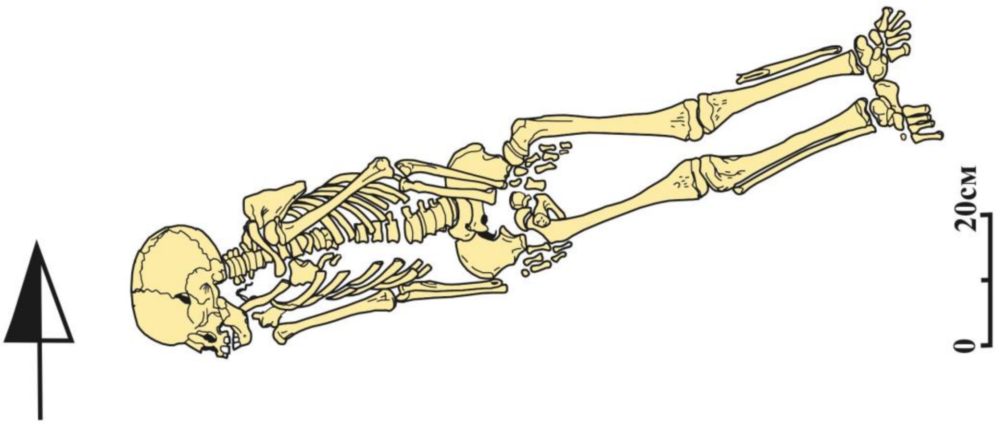

***Fig. SI.B.20.*** *Burial 5 at the Ust-Menzel'ya hillfort.*

##### I.B.10.3. Zuevy-Klyuchi hillfort

The site is located on the right bank of the Kama River, 1.5 km southwest of the village of Zuevy Klyuchi in the Karakulinsky district of the Udmurt Republic. The settlement occupies a promontory with a roughly triangular shape on the slope of the southern exposure of the third floodplain terrace, formed by two steep slopes with a depth of 20-25 m. Excavations conducted

by R.D. Goldina, V.F. Gening, and L.I. Ashikhmina yielded approximately 5000 square meters of the unearthed cultural layer, yet the site remains largely unpublished. Materials from the settlement were published by V.F. Gening and L.I. Ashikhmina in 1984, with a characterization of early Iron Age dwellings identified at the site in 1986. From 1997 to 2006, the hillfort and several adjacent sites were excavated by E.M. Chernykh. To the west, across the ravine, lies the Zuevy-Klyuchi burial ground of the Anan'ino period, excavated by A.A. Spitsyn in 1898 (unpublished).
In the course of the excavations of the Zuevy-Klyuchi hillfort in 1971-1972, at the southeastern edge of the promontory and beneath the cultural layer of the Late Anan'ino period, numerous burials were unearthed. 25 of them were attributed to the Seima-Turbino period, as reported by V.F. Gening in 1975 while the others were dated to the Late Anan’ino and “later” periods. The anthropological material from the excavations was stored in the Tomsk State university.
In total, we sequenced eight individuals from the burial ground at the Zuevy-Klyuchi hillfort. Five of them were dated by the 14C, and all of them show similar ages between 1227 and 1394 CE. The other three individuals are genetically close to the distinkt group of the dated samples. Based on this observation, we attribute all eight individuals from Zuevy-Klyuchi to the Late Medieval period.

##### burial 30 (individual ID I30341, Female)

The grave pit, of sub-rectangular shape measuring 120 x 80 cm, oriented in the direction of WNW-ESE, was identified at a depth of -63 cm. The depth of the pit is -16 cm. The burial contained a skeleton in satisfactory preservation, lying in an extended position, with the head to the WNW. The upper part of the skeleton, from the position on the back, is partially turned onto the right side, the left arm is semi-flexed at the elbow, and the right arm is extended alongside the body. Near the knee joint of the left leg of the deceased, a fragment of pottery was found at a depth of -73 cm from the surface, and a human tooth was found in the NE part of the burial.
The determined radiocarbon age of the burial is 1292-1395 calCE (635±20 BP, PSUAMS-13158) (based on human bone).

##### burial 34 (individual ID I30342, Female)

Human bones were discovered in the sod-plowed layer at a depth of 14 cm in the southeastern corner of the plot. The skeleton is heavily damaged by plowing and is preserved fragmentarily, with better preservation of the leg bones. Due to the damage to the skeleton, it is difficult to judge its orientation. Apparently, it was oriented from S3 to E, with the head to S3. Two pottery fragments were found near the skeleton: the first near the presumed pelvis at a depth of 14 cm from the surface, and the second at a depth of -24 cm near the knee joint of the right leg.
The determined radiocarbon age of the burial is 1303-1401 calCE (605±20 BP, PSUAMS-13394) (based on human bone).

##### burial 43 (individual ID I32547, Female)

The determined radiocarbon age of the burial is 1296-1394 calCE (635±15 BP, PSUAMS-12694) (based on human bone).

##### burial 42 (individual ID I30343, Female)

The determined radiocarbon age of the burial is 1318-1407 calCE (585±15 BP, PSUAMS-13159) (based on human bone).

###### burial 45 (individual ID I32546, Female)

The determined radiocarbon age of the burial is 1227-1278 calCE (775±15 BP, PSUAMS-12708) (based on human bone).

### I.C. The Middle Kama and Cis-Urals (including the Belaya Basin)

#### I.C.1. The Early Iron Age Pyany Bor culture in the Southern Cis-Urals (Belaya_Pyany-Bor)

##### I.C.1.1. Kipchakovo

The Kipchak I kurgan-ground burial ground is located 1 km east of the village of Minnyarovo (Aktanyshsky district, Republic of Tatarstan) and 2.5 km southwest of the village of Kipchakovo (Ilishovsky district, Republic of Bashkortostan), on the main terrace of the right bank of the Syun River, 70-80 m southwest of the outer rampart of the Kipchakovsky settlement.

Over the course of four field seasons (1990–1991, 1994, and 2001), F.M. Tagirov excavated 4 kurgans (Nos. 15, 41, 51, 52) and two adjacent excavations, one near kurgan No. 41 (Excavation I, area 36 sq. m) and the other near kurgan No. 15 (Excavation II, area 618 sq. m). A total of 64 burials were excavated during this time.

Later, the site was excavated by S.E. Zubov. Some reports were not written. The numbering of burials was restarted several times: numbers up to 20 are encountered at least 3 times, and up to 50 at least 2 times. Determining the complex and dating is difficult. Overall, the findings do not extend beyond the Pyany Bor culture.

##### I.C.1.2. Starokirgizovo

Staro-Kirgizovo village, Ilishovsky district, Republic of Bashkortostan. Right bank of the Myadym (Minueshta) River, a right tributary of the Belaya River, a left tributary of the Kama River. Located on the eastern outskirts of the village near the tractor park and fire station, on a high promontory formed by the edge of the right floodplain terrace and a ravine. Discovered by A.Kh. Pshenichnyuk in 1971 (information is not available in reports; materials not deposited in museum collections), two burials were excavated. In 1971-1972, 51 (or 53) burials were excavated. In 2016-2017, work was continued by N.A. Lifanov, who excavated 75 burials. The buried individuals were lying extended on their backs, with their heads to the south or southwest. The burials yielded a rich material assemblage, including adornments (beads, temple pendants, bronze plaques, pendants, openwork overlays), weapons (arrowheads, knives), and ceramics.

The numbering of burials is duplicated by different authors. It is possible that some graves were excavated twice.

###### burial 25 (individual ID I16644, Female)

The burial pit was not documented and had been completely destroyed by several trenchings; two lines of cables (now inactive) ran through the burial. Overall, the skeletal remains retained anatomical order, although some bones were displaced, mixed, or missing. The buried woman, aged between 45 and 65 (closer to 65), was laid out extended on her back with her head to the northeast.

The burial inventory included items made of bronze, iron, and glass. Some of them, apparently, had been displaced from their original positions. For instance, fragments of bronze temple spiral pendants were found in various locations (to the northeast and southwest of the skull, near the right elbow, abdomen, sacrum, and by the right knee). Two round bronze plaques were found to the east of the skull and on the sacrum. Three round bronze discs were discovered near the right pelvic bone, and a bronze rhombic plaque was found under the pelvis. Iron items were concentrated in the lumbar-pelvic region. Fragments of iron items (knives?) were found around the abdomen and under the right pelvic bone. Fragments of iron bracelets were found on both sides of the pelvis, and an iron waist hook was fixed to the southeast of the skull. Yellow glass beads, totaling five specimens, were located to the east of the skull, on the sacrum, and by the right knee. Blue glass beads (two specimens) were found to the east of the skull and on the right side of the chest. Adjacent to the latter were green beads. A cluster of various beads, including white and blue, gilded, as well as polychrome beads ornamented with longitudinal blue and white stripes (totaling 49 specimens), was cleared on the left side of the chest.

The determined radiocarbon age of the burial is 94 calBCE - 58 calCE (2030±20 BP, PSUAMS-10313) (based on human bone).

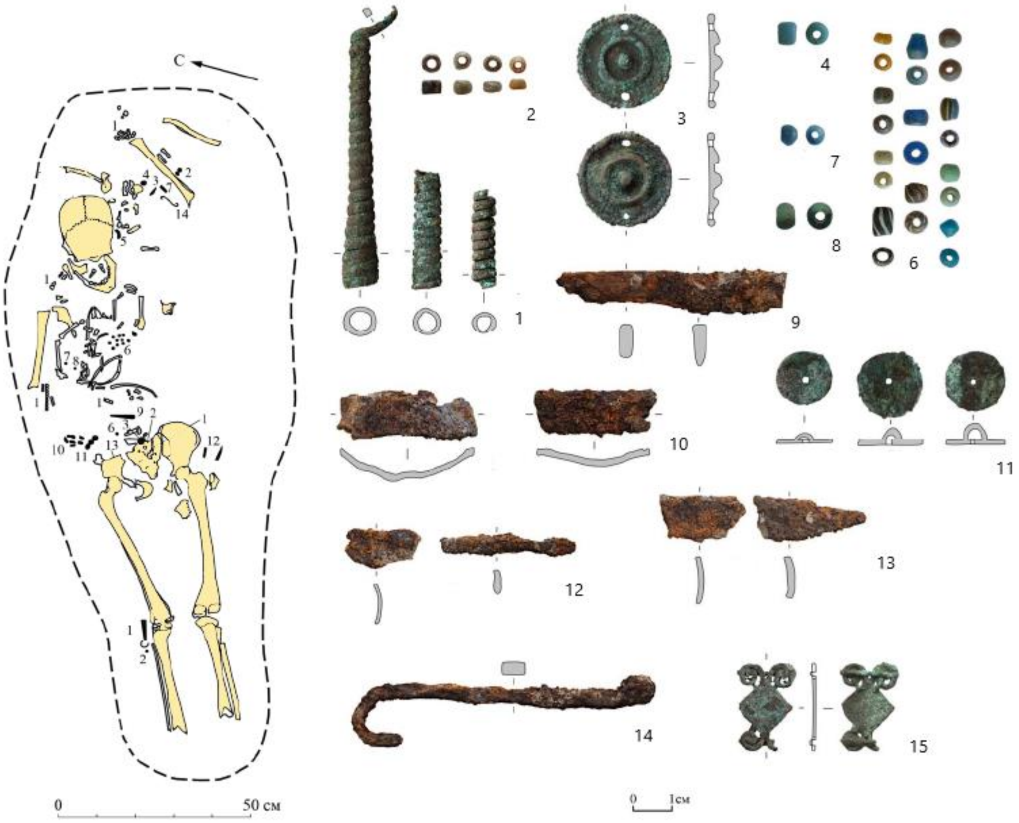

***Fig. SI.C.1.*** *Burial 25 at Starokirgizovo cemetery.*

##### burial 42 (individual ID I16645, Male)

The burial pit was elongated with rounded corners and oriented along an east-west axis. The dimensions of the burial pit were as follows: length – 224 cm; width – 68-75 cm; maximum depth of the pit from the discovery level – 21 cm (at the western wall). The walls were steep, and the bottom was flat.

Overall, the skeletal remains retained anatomical order, although some bones had decayed. A cluster of small chalk fragments was found under the skull. The buried man, aged between 20 and 28, was laid out extended on his back, with his head oriented towards the eastern sector.

The burial inventory mostly retained its in situ position. Fragments of a bone buckle and a wooden object with iron oxides were found between the pelvic bones. Two bone triangular socketed arrowheads, one of which showed traces of iron oxide, were found near the left femur and the western wall of the pit. Fragments of an iron three-bladed tang arrowhead with remnants of a shaft were found between the left tibia and the southern wall of the burial pit. South of each of the major tibia bones, there was a bronze circular framed buckle with a single pin, likely belonging to footwear. Among the bones of the right foot, a bone disc with a through hole in the center was found. In the southwest corner of the pit, adjacent to the bones of the left foot, there was a long bone awl with a hole. Additionally, fragments of burnt animal bone were found near the left scapula and right pelvic bone.

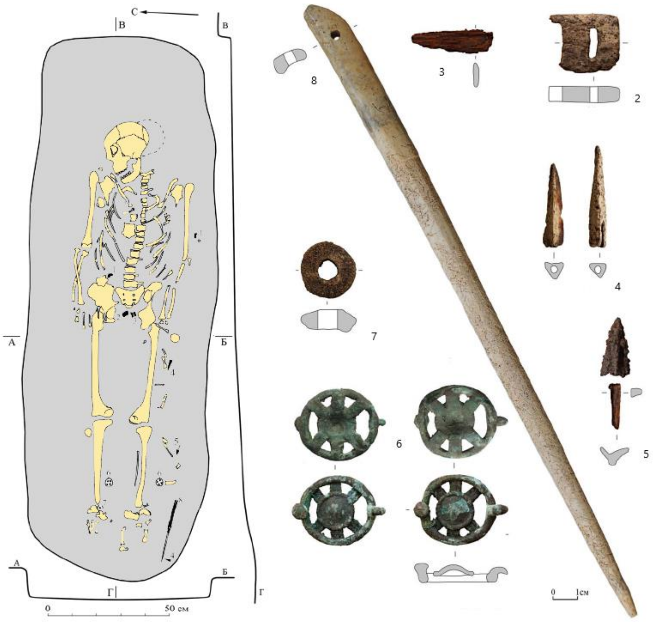

***Fig. SI.C.2.*** *Burial 42 at Starokirgizovo cemetery.*

##### burial 53 (individual ID I16643, Male)

The burial pit was elongated with rounded corners and oriented along a northwest-southeast axis. The dimensions of the burial pit were as follows: length – 200 cm; width – 70 cm; maximum recorded depth of the pit from the discovery level – 19 cm (at the southern wall). The walls were mostly steep, and the bottom was flat, with a slight slope towards the south.

Overall, the skeletal remains retained anatomical order, although some bones were displaced due to earthmoving activities. The buried child, aged 7-8 years, was laid out extended on their back, with the head oriented towards the north-northwest. The arm bones were laid alongside the body, and the skull was positioned on the base of the skull and the occipital bone, slightly tilted to the left side.

To the left side of the skull, a bone socketed arrowhead with a rhombic cross-section was found. In the neck area, west of the skull, two yellow beads and one blue bead were found east of the skull. Near the left shoulder, a small fragment of calcined animal bone was found. In the waist area, there was a rectangular bone buckle with a fixed hook and two through slots perpendicular to the buckle axis for threading a belt. Three bronze elliptical eyelets were found near the left wrist and hand. Another similar eyelet was found between the buried individual's thigh bones, next to which was a triangular bone socketed arrowhead. Another bone socketed arrowhead with a rhombic cross-section was found near the right knee.

On the outer side of the right thigh bone, there was a poorly preserved iron knife approximately 30 cm long. The knife was in wooden sheaths, from which wood decay was detected during clearance. In the area of the handle, near the thigh, there was a trapezoidal bone clasp with a slit in the wide part and two oval-shaped holes in the narrow part. Judging by its position, the clasp served to attach the sheaths to the belt. Four claw phalanges of an adult badger from the front limbs were found on this clasp. Nearby, on the inner side of the right thigh bone, a stone spindle whorl with a hole was found.

Between the knife and the upper part of the thigh bone, from top to bottom, there were the following: a permanent premolar of an adult wild boar from the right mandibula, a canine tooth of an adult horse from the left upper jaw (with an artificial round hole in the root and the end of the root cut off), a fragment of the canine tooth of an adult wild boar from the right upper jaw (with an artificial round hole in the root), and a fragment of a jawbone of a badger with a canine tooth.

In the area of the knife tip, near the right shin, there was a fastening for attaching the end of the sheath to the leg in the form of a composite "bracelet". This consisted of a bronze clasp with a fixed hook of elongated trapezoidal shape, decorative bone plates of bow-like shape with a central through hole for a leather strap (7 pieces), and separating bronze eyelets between them (7 pieces). The terminal part of the attachment was decorated with five bone pieces with a central hole, imitating animal fangs. Below the knife sheath, a circular bronze plate with a hole in the center and two ears on the back for threading a strap was found. This plate was presumably part of the ornamentation system for attaching the terminal part of the combat knife to the leg.

On the inner side of the right thigh bone, there was a complex consisting of cut upper and mandibulas of small predatory animals, originating from at least two badgers, two foxes, and one otter. From top to bottom (from the groin to the knee), cuts made during the detachment of the front part of the skull were evident on the upper parts of the jaws. The cut passed through the ends of the roots of the upper canines.

Between the legs, in the area of the left foot, a cluster of bone socketed (18 pieces) and iron three-bladed tang arrowheads (6 pieces), apparently stored in a quiver, was found. At the end of the shins, two bronze round shoe fasteners with a fixed hook and a corresponding pin were discovered.

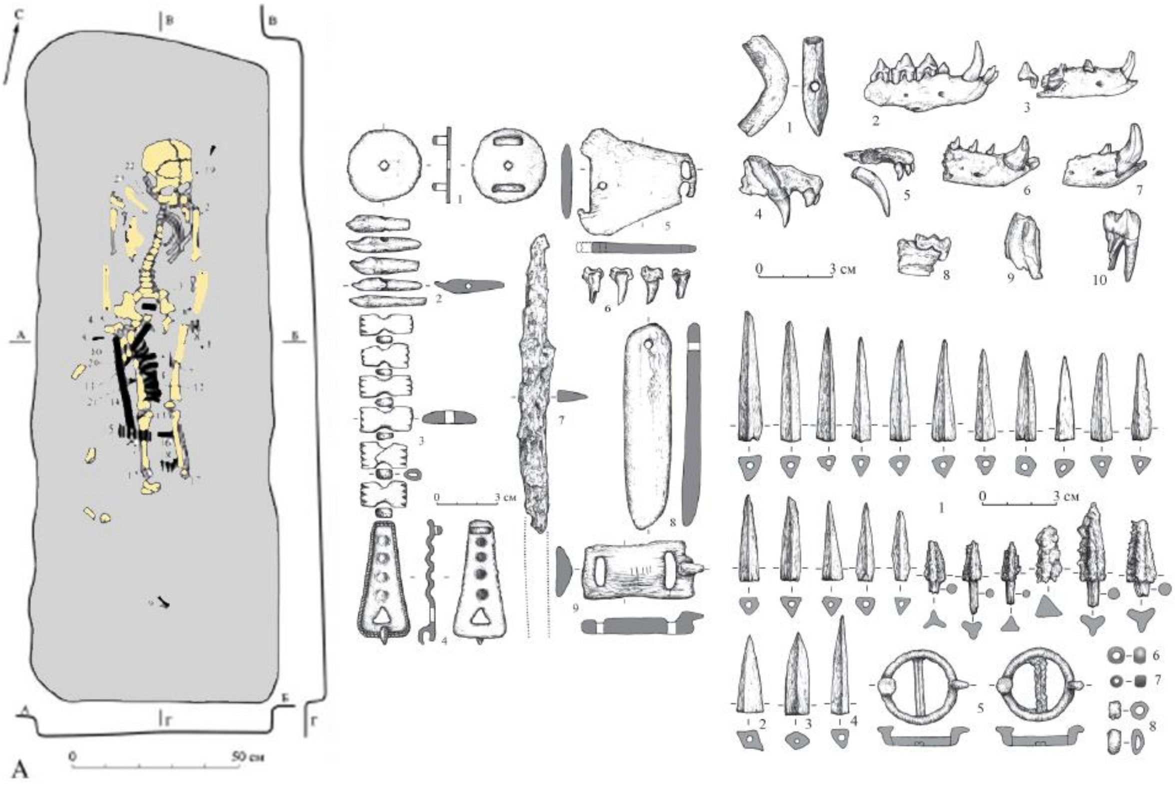

***Fig. SI.C.3.*** *Burial 53 at Starokirgizovo cemetery.*

burial 58 (individual ID I16642, Male)

The burial pit was elongated with rounded corners and oriented along a west-east axis. The dimensions of the burial pit were as follows: length – 164 cm; width in the central part – 56 cm; the maximum recorded depth of the pit from the discovery level – 28 cm (in the western part). The actual depth of the pit from the ancient soil surface, judging by the profiles, was at least 45 cm. The walls of the pit were mostly steep, and the bottom was flat with a slight slope towards the west and south.

The skeletal remains were partially damaged by the cable trench. Four arrowheads were found in the burial: two triangular bone socketed and two three-bladed bronze socketed arrowheads. The bronze arrowhead was positioned in the left part of the chest of the buried individual, with the tip pointing south, while the second bronze arrowhead and both bone arrowheads were scattered disorderly along the left shin of the skeleton, likely displaced as a result of the intervention that also damaged the foot bones.

Along the right thigh of the buried individual, an iron knife was found, with remnants, likely part of its handle, consisting of small fragments of iron and wood, near the right wrist. A fragment of a bone was fixed in the chest area, and a complete round bronze buckle was found near the left thigh. On the right side of the skull, a silver(?) pendant was unearthed.

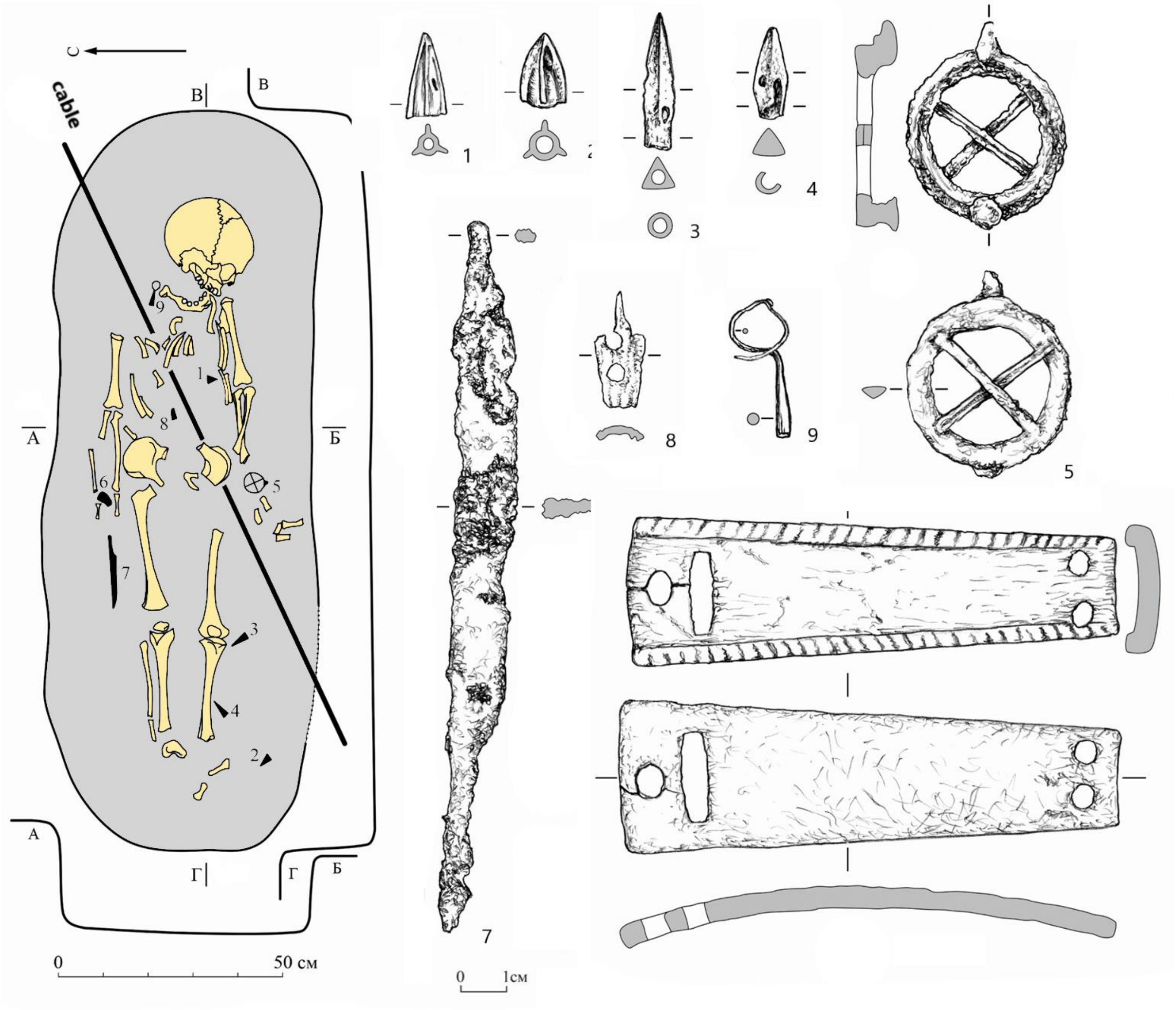

***Fig. SI.C.4.*** *Burial 58 at Starokirgizovo cemetery.*

##### burial 63 (individual ID I16641, Male)

The burial pit was elongated with rounded corners and oriented along a west-east axis. The dimensions of the burial pit were as follows: length – 208 cm; width – 70 cm; the maximum recorded depth of the pit from the discovery level – 17 cm (at the western wall). The walls of the pit were steep, and the bottom was flat with a slight slope towards the west and south.
The skeletal remains of an adult male, approximately 45-55 years old and in good condition, were located closer to the southern wall of the pit. The buried individual was lying stretched out on his back with the head to the east, the right arm stretched along the body, and the left hand on the pelvic bone.
The burial inventory included an iron knife lying above the right femur and a set of quiver remnants near the left shin. The attachment of the quiver to the leg was marked by bronze spiral-wound plates around the tibia bones and a bronze pin. The best-preserved plate was wound three times, while the others were fragmented. The pin had a truncated biconical shape. Judging by the preserved decay, the quiver was made of leather. Its contents consisted of arrowheads made of bronze and bone with partially preserved shafts. There were 18 bronze three-bladed socketed arrowheads and 2 triangular bone socketed arrowheads. Fourteen shaft fragments were found.

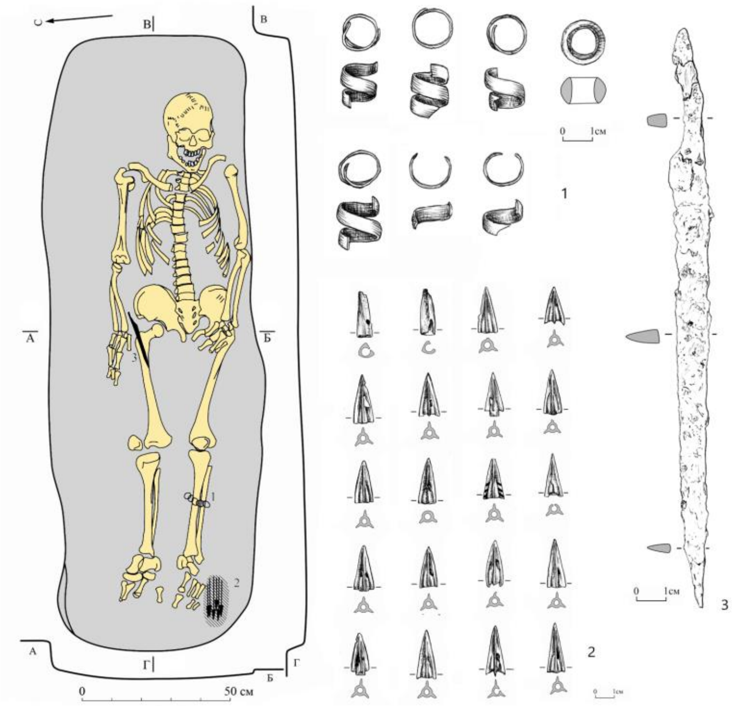

***Fig. SI.C.5.** Burial 63 at Starokirgizovo cemetery.*

#### I.C.2. The Migration period Mazunino culture in the Southern Cis-Urals (Belaya_Mazunino)

The description of the Mayunino culture assemblage of the Lower Kama region was given in the I.B.4 section of this Supplementary material.
In the Cis-Ural area, the Mazunino sites demonstrate similar cultural traits. In this region, however, the Mazunino sites do not disappear after the 5th century CE, but gradually evolve to the Bakmutino cultural tradition.
The first finding of the artifacts later attributed to the Bakhmutino tradition became known in the late 19th century (Bulychev N.I., 1902), and systematic research started from the 1920s (Schmidt A.V., 1929). In the late 1950s and 1960s, the Bakmutino cultural assemblage was

characterized as a distinct phase of the local cultural development (Mazhitov N.A., 1963a/r; 1968).

Fortified settlements with ramparts and ditches attributed to the Bakhmutino culture are located on high floodplain terraces, while settlements are situated on lower ones. Dwellings consisted of above-ground log structures up to 120 square meters with hearths and storage and cash pits. The pottery of the early period is represented by small, low-necked round-bottomed bowl-shaped vessels ornamented with dimples across the entire surface; of the late period - by large vessels of the same type ornamented with dimples and triangular punctures across the entire surface, with incisions on the rim and shoulders. The main branches of the economy were animal husbandry, agriculture, hunting, fishing, iron, and bronze metallurgy.

The Bakhmutino burial grounds are flat, with rows of grave pits. During the early phase of the culture, the deceased were buried in shallow graves in a supine position. The burial inventory consisted of tools (spindle whorls, iron knives, awls, sharpening stones, and sickles), weapons (iron and bone arrowheads, spears, axes), decorations (bronze bracelets, temple pendants, neck rings, appliqué plaques, pendants in the form of ducks, bears, necklaces made of glass beads). The "sacrificial complexes" consisting of women's jewelry are specific details of the Early Bakhmutino burials.

In the late period, the deceased were buried in deep graves (sometimes in wooden log coffins or wrapped in bark) in a supine position. Clay vessels and remnants of sacrificial food (horse and sheep bones) were found at the head, in the corners. Male burials were characterized by the presence of weapons (iron arrowheads, spears, swords), and female burials by jewelry (bronze, gold, silver lunar pendants, zoomorphic temple pendants, bronze mirrors, bronze and iron buckles). For adult burials, the presence of belt fittings (bronze and silver overlays, buckles, pendants, belt tips, including those covered with gilded foil) was characteristic.

We publish four individuals from the Cis-Ural Birsk-2 cemetery, the three are attributed to the Mazunino culture and one to the early phase of the Bakhmutino tradition.

##### I.C.2.1. Birsk-2

The Birsk-2 burial ground is located at the southern outskirts of the city of Birsk and occupies a vast area of the native terrace of the right bank of the Belaya River, 0.6 km northeast of the Birsk (Chertovo) hillfort. The total area of the site reaches 350x450 m or approximately 157,000 m².

The Birsk-2 burial ground has been known since 1902, when N.I. Bulychev described finds stored in the Ufa Governorate Museum (now the National Museum of the Republic of Bashkortostan) (Bulychev, 1902). Over many years of research on the burial ground, conducted by Niyaz Mazhitov, six excavations were conducted, and 692 burial complexes dated to the 8th-10th centuries AD were studied (Mazhitov, 1968). In 2017 and 2021, the work was continued by Rida Ruslanova.

###### burial 694 (individual ID I19264, Male)

The burial pit of a sub-rectangular shape and oriented south-south-east - north-north-west was discovered at a depth of -0.62. It measures 1.9x0.64 m (width at the bottom 0.32 m) with a depth of 0.35 m. An oval-shaped spot of small charcoal pieces (15x24 cm) was detected incide the burial pit infill of black soil with clay inclusions. The grave pit had shoulders, possibly used for construction of wooden covering, which remains were recorded in the profile of the burial pit. Deeper in the infill, fragments of a human mandibula (in the southwest corner) and traces of almost completely decayed skull (in the central part) were unearthed.

On the bottom of the grave pit, a postcranial bones of a skeleton of an adult aged 25-30 years was excavated, positioned supine. It is possible that the skull, previously cleared higher, belongs to this individual. There are no signs of trauma on the cervical vertebrae. Below the skeleton (in its upper part), remnants of organic bedding were found.
In the infill, remains of a collapsed thick-walled clay vessel of reddish-brown color with dreg impurities in the clay, placed along the burial axis, were cleared. On the ledge on the eastern side, a fragment of a red clay vessel with coarse sand impurities and a fragment of black ceramic with dreg impurities were identified.
At the level of the cervical vertebrae, a bronze pendant in the form of a bear with well-defined protruding ears was found. In the area of the cervical vertebrae, a fragment of a ring wire temple pendant with a wound faceted ribbon was positioned. Under the elbow joint of the left arm, a heavily corroded iron buckle with an oval frame and previously movable tongue was identified. Decayed remnants of an iron knife were preserved on the pelvic bones. Above it, parts and decayed remnants of iron bit were discovered.

burial 702 (individual ID I19265_d, Male)

An oval burial pit 2.1x0.63 m oriented along the north-north-west - south-south-east line was recorded at a depth of -0.80 m. The depth of the grave pit was The filling contained scattered calcified bones and 2 teeth (fangs), from which it is possible to determine the approximate age of the buried person as 35-45 years old.
A collapse of a handmade pottery vessel and an iron buckle with a round-shaped frame and a movable tongue were found at the bottom of the grave pit.

burial 710 (individual ID I19266_d, Male)

The grave pit oval in shape oriented along the the north-north-west - south-south-east line was discovered at a depth of -0.59 m. It's dimensions were 2.44x0.90 m, and depth 0.32 m. The grave pit was cover by wooden boards, placed in ledges in the northern, eastern and southern walls.
At the bottom of the grave pit, a skeleton of a 30-35 year old male was found, stretched out on his back and with his arms located along the body. In the area of the ribs, remains of a leather belt and a bronze belt-tip were found made from a bent plate, with an extension in the bent part. A bead of green glass was also found here.
At the knees of the skeleton, a cluster of objects was discovered, including an axe, an iron spearhead, and a belt tip. In the southern part of the burial, near the bones of the right foot, a ritual assemblage was discovered, consisting of a bracelet, inside of which were placed 4 Mazunino-type pendants (2 of them had embedded beads of white and blue glass). A bronze temple wire ring wrapped in a bronze ribbon and an iron tetrahedral awl were also cleared here.

burial 713 (individual ID I19267, Female)

The oval-shaped burial pit measuring 1.4x0.62 meters was discovered at a depth of -0.41 m and had a depth of 0.22 meters. The burial pit had a flat bottom and a ledge in the southeastern wall. The orientation of the burial was northwest-southeast. The skeleton was placed within a wooden structure, secured by decayed material. From the skeletal remains, a crushed skull turned to the left and a shoulder bone were preserved. The age of the individual, approximately 4 years old, was determined based on the discovered bones (teeth).
The skull was likely covered with a cloth with beadwork, which fell over a ritual assemblage placed hear the head of the deceased person. The assemblage included a ring of Mazunino type, a twisted temple ring, rhombic-section torque, beads, chains (small and large), round temple

pendant, and a triple-shaped appliqué. A belt buckle was discovered in the central part of the burial, with two additional buckles found in the southern part of the burial.

##### I.C.3. The Kushnarenkovo culture (Belaya_Kushnarenkovo)

The terms Kushnarenkovo and Karayakupovo cultures refer to cultural traditions of the Cis-Ural population dated to the 6th-9th century CE and distinguished mainly on the features of handmade pottery making. However, these definitions have been actively discussed for more than half a century. Some researchers consider burials and settlements with the Kushnarenkovo-type pottery as an archaeological type (Khalikova, 1976) or clearly defined archaeological culture of the 6th-10th centuries (Vasyitkin, 1968; 1992; Kazakov 1981; 2001). Other researchers believe that this is the early chronological stage of the Karayakupovo culture (Mazhitov 1977; 1981a; 1981b), united Kushnarenkovo-Karayakupovo culture (Botalov, 2019), two independent synchronous cultural groups (Matveeva G., 2007; Ivanov A., 2008) or two asynchronous cultures of the late 6th – early 8th and mid 8th - 9th centuries (Ivanov V., 1999; 2006). The most radical version supposes the origin of handmade Kushnarenkovo-type ware as a product of “traveling potters” (Matveeva N., 2021). This discussion is too far from the ending, therefore we have to use the term "horizon", denoting the time of the highest spread of this cultural phenomenon.

Kushnarenkovo horizon

The distinctive features of Kushnarenkovo pottery were characterized only by the results of the excavations of an ensemble of sites discovered near the village of Kushnarenkovo in the second half of the 1950s, despite some previous discoveries (Gening, 1977). Interestingly, Kushnarenkovo pottery was not predominant or even notably represented on any of these sites, including burial grounds, settlements, and hillforts. Despite the presence of Kushnarenkovo ceramics, systematic characterization of the culture is lacking, and its chronology remains undeveloped. Existing concepts suggest a prehistory of ceramic tradition from the Trans-Urals region, although the date of the culture's origin remains uncertain.

There is evidence of the use of semi-subterranean dwellings at the Kushnarenkovo settlement, while others were likely light surface structures. Ceramic vessels are characteristic, with medium and tall vessels featuring high vertical necks decorated with fine diagonal notches around the rim, horizontal carved lines along the neck and upper body, rows of inclined impressions from toothed stamps, and fine diagonal mesh patterns. Kurgans at burial sites range from 6 to 10 meters in diameter and up to 0.5 meters in height, with 1-3 (rarely up to 5) burials in rectangular or oval pits 40-60 cm deep from the burial soil level. The bodies are placed supine, with the head facing north or west, occasionally ritually disarranged. Among the artifacts are ornaments, horse equipment, weapons; vessels are usually placed at the head, with food remains (horse or sheep bones) nearby. Skulls and limb bones of horses are found under mounds or in graves, possibly with the animal's skin. Other findings include an iron sword in wooden scabbards, arrowheads, fishhooks, bridle rings, stirrups in later complexes; bone inlays for bows; a silver chain necklace (Taktalachuk burial ground); silver and bronze belt fittings, earrings, pendants, rings; beads, including cornelian with white inlay.

The earliest dateable artifacts found together with the Kushnarenkovo-type pottery are belt sets and variuos strap ornaments of “heraldic” styles (late 6th – early 8th centuries). Later the characteristic features of the Kushnarenkovo pottery traditions are observed in complexes with the Karayakupovo-type ware (Matveeva G., 2007; Ivanov V., 2009). However, the

Karayakupovo-type tradition dominate most of these sites with the exception of the Bolshiye Tigany cemetery (9th – early 10th century).

Karayakupovo horizon

The eponymous hillfort of the Karayakupovo culture has been studied since the early 1960s (Matveeva G., 1975; Ivanov V., 1990a; 1999).

Hillforts are located on elevated promontories (some without traces of earthworks; some are shelters), while settlements are on river terraces with a sparse cultural layer. Characteristic pottery is round-bodied, with ovoid, spherical, or pear-shaped bodies and straight, slightly curved necks. The ornamentation consists of diagonal impressions from short smooth or toothed stamps on the rims, horizontal lines on the neck and shoulder, waist belts of pits and protruding "pearls." Karayakupovo-type ceramics are found in subterranean(?) and sub-barrows burials at sites with a broader chronological range (Manyak, Karanaevo, Beketevo, Idelbaevo). The associated context contains spindle whorls of truncated conical shape, crucibles, spoon-molds, fishhooks. The economy was likely based on livestock and farming (charred grains of spelt, millstone fragments were found).

The earliest Karayakupovo-type ware was found in complexes with "heraldic" style ornaments of the late 7 – early 8th centuries CE. Starting from the mid 8th century, the prestigious fashion of the population of the region (belt ornaments, harnesses, weapons etc.) changes under the impulse of the early Saltiv culture, on the other hand - under the influence of the Turkic nomads of Central Asia. In the middle of the 9th century they were both replaced by new nomadic impulses of influence, among which the Proto-Magyar and Srostki culture, and later in 10th century CE also Oghuz-Patzinaks culture.

##### C.3.1. Bustanaevo

The Bustanaevo burial kurgans were discovered in 2011, with initial investigations conducted by Aleksandr Kolonskikh in 2015 (Kolonskikh, 2017a and 2017b). During reconnaissance work, 43 mounds were identified, and the presence of two concentration zones of mounds (NW and SE) was established. The necropolis is located on the edge of the main terrace on the watershed of the Bystry Tanyp and Gareika rivers. We sequenced one individual found in burial 1 under mound 45.

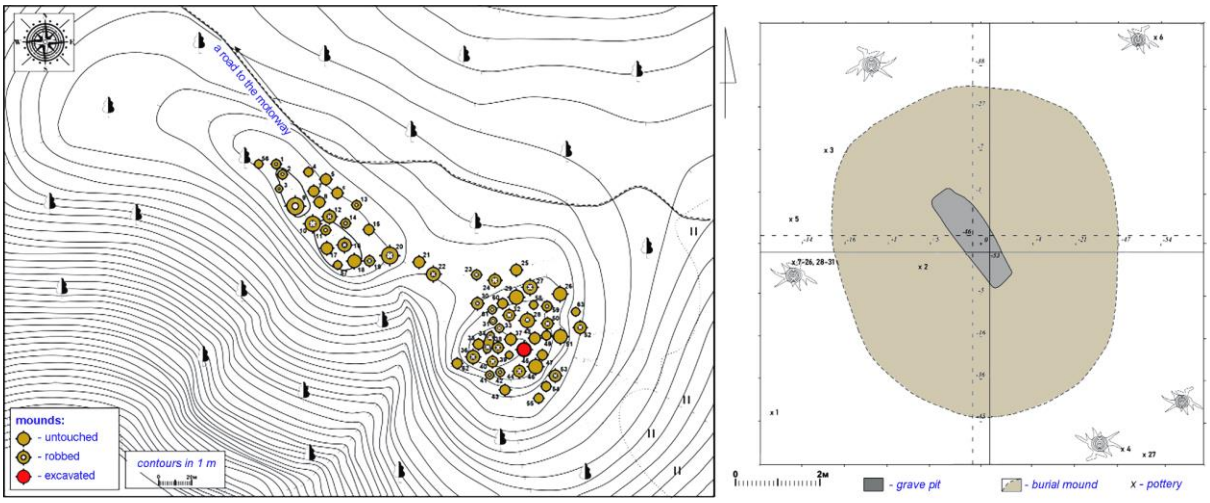

***Fig. SI.C.6.*** *Bustanaevo kurgans. A plan of the kurgan group and a plan of the excavated kurgan 45.*

##### mound 45, burial 1 (individual ID I25531, Male)

Burial 1 under mound 45 had a sub-rectangular shape with rounded corners. The length of the burial pit is 235 cm, the width in the southeast part is 57 cm, in the northwest part is 50 cm, and at its widest point is 64 cm. The depth of the burial pit varies from 79 to 91 cm from the ground level. The bottom of the pit had a slight depression in the center. The walls are vertical, without additional steps, ledges, or shoulders, although a small recess was made under vessel 1 in the western wall. The mound is oriented along the northwest – southeast line, with the deceased placed head to the northwest, which is typical for the Kushnarenkovo burial rite. The skeleton was poorly preserved, only fragments of the skull, shoulder, and femur bones were discovered.

In the burial, flower-shaped four-petal overlays, a buckle fragment, a belt tip, two vessels, five arrowheads, elements of horse equipment, knife fragments, and a pendant were found. The Kushnarenkovo-type vessels were located in the northwest part of the burial: vessel 1 stood west of the skull in the niche at the western wall, vessel 2 laid on its side and was crushed. Fragments of animal bones, likely from sheep, were found in vessel 2, indicating food offerings.

The determined radiocarbon age of the burial is 340-540 calAD (1640±25 BP, DeA-16570) (based on human bone), however, the grave goods confirm its irrelevance. Based on archaeological data, the grave must be dated to 560-630 CE. Pottery confirms that the burial is associated with the Kushnarenkovo archaeological culture.

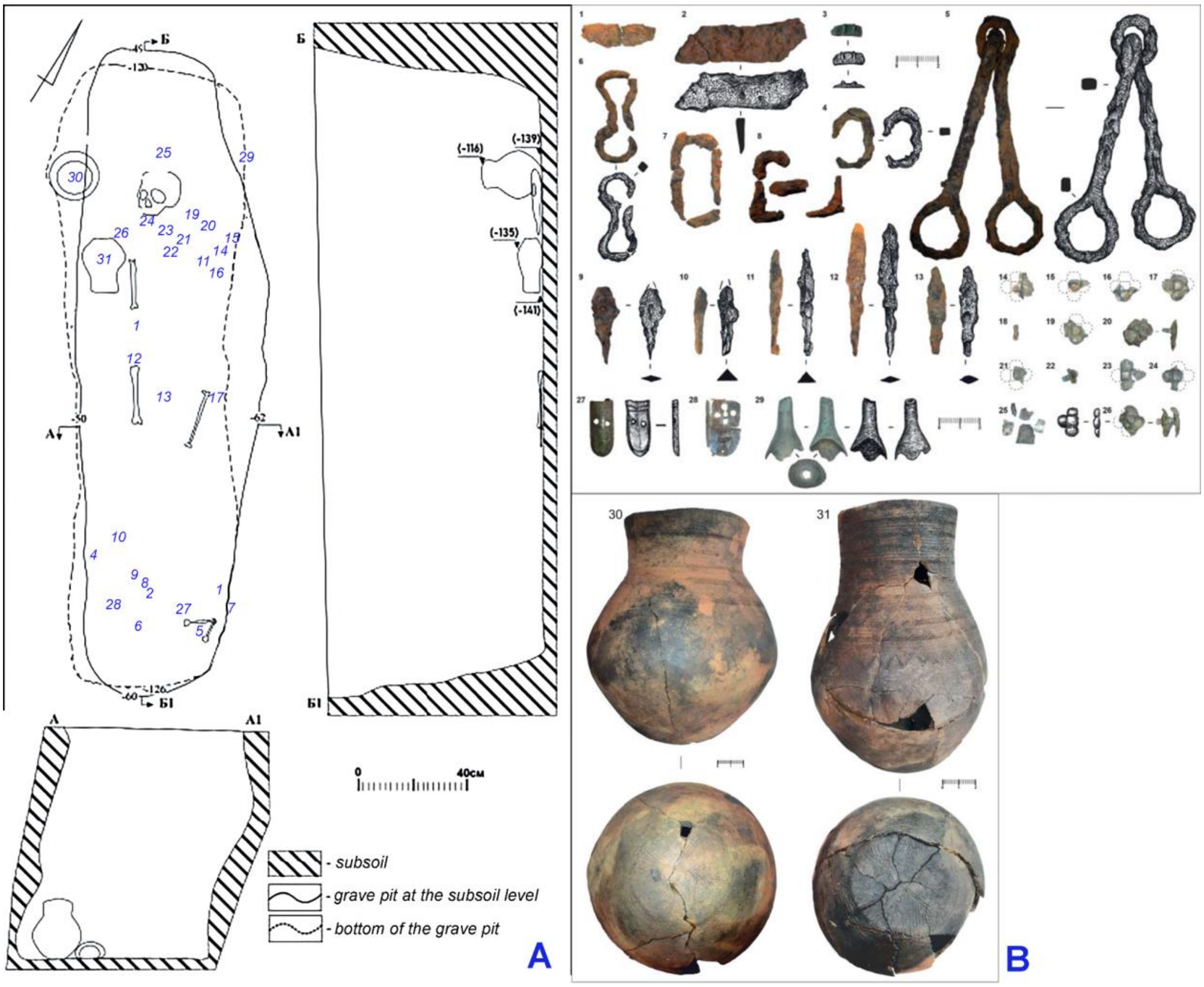

***Fig. SI.C.7.*** *Bustanaevo, mound 45, burial 1. A - a plan of the burial, B - grave goods.*

#### I.C.4. The Nevolino culture in the Cis-Urals (Cis-Urals_Nevolino)

The main features of the Nevolino culture were described in the section I.B.5.

##### I.C.4.1. Bartym

The Bartym-1 burial ground occupies the southwest part of Bartym-1 settlement of the Nevolino culture. In 1980, 1981, and 1983, during excavations conducted by the archaeological expedition of the Udmurt State University, an area of about 2900 square meters was excavated, revealing 19 burials containing 60 skeletons.

The upper part of the cultural layer in this area was completely destroyed by plowing. The burial pits were identified after the removal of the plow layer at a depth of 20-25 cm. The bedrock at the site is limestone with clay lenses. Some burial pits were carved into solid rock (8 graves - 42.1%), while others were dug into clay (11 graves - 57.9%). One of the burials was made within a cash pit of the settlement. Some graves were disturbed by plowing.

###### burial 16, individual B (individual ID I19110, Male)

The burial level is -65 cm. The grave has square outlines with rounded corners, dimensions 183x158x80 cm, oriented along the east-south-east - west-north-west line. At the level of -65-80 cm, nine skeletons were found, oriented with their heads to the ESE. Four skeletons (A, B, C, G) were arranged in a row; four belonging to children (D, E, F, H) were found at the feet of skeletons A, B, and C; the fifth (I), from which only a fragment between the thigh bones was preserved, had 3 bronze plates, a bronze buckle on the right shin, 3 plates, and a ornament consisting of bird-shaped, swollen, and horn-shaped ornaments, 10 glass beads. At the northern end, between the skulls B and F, a set of 9 bronze tubular ornaments and a bronze rattling pendant were found. In the area of the belt of skeleton C, next to the tubular arm bones, 4 glass beads, 2 bronze wheel-shaped pendants, and 3 plates were found. To the left of these tubular bones, 6 bronze plates, a buckle, a belt tip, and a fragment of an iron knife were found. Between the fragments of skeletons C, F, and E, fragments of an iron knife and a bronze buckle were found. Slightly below, at the head of skeleton E, bronze plate fragments and a buckle were found to the left of the legs of skeleton F.

Anthropologically determined sex and age-at death of the buried individuals is as follows: A - male, 18-20 years old; B - male, 35-45 years old; C - male, 30-40 years old; D - sex?, 5 years old; E - gender?, 4 years old; F - gender?, 5 years old; G - gender?, 6 years old; H - gender?, 0.5 years old.

The determined radiocarbon age of the burial is 428-591 calCE (1535±24, DeA16585) (based on human bone from individual D).

#### I.C.5. The Lomovatovo culture (MidKama_Lomovatovo)

The interest in the "cult" finds of the Lomovatovo type emerged in the mid-19th century (Spitsyn, 1902). Systematic excavations primarily targeted burial complexes and were generally characterized by Rimma Goldina in her dissertation (Goldina, 1970s; 1971a/b; 1985; Goldina, Kananyin, 1989). Subsequent work became sporadic, only intensifying in recent years. Synthesizing a broader chronological range led A.M. Belavin to establish a new periodization (Belavin, Krylasova, 2016).

Settlement clusters encompassed hillforts, settlements, and burial grounds. The sites attributed to the Lomovatovo culture also include several small late *kostisches*. Houses had sunken floors, featuring hearths and storage pits, pillar constructions, and adobe floors, with some displaying sidewall trenches and earthbank reinforcements. Pottery was handmade (often shell-tempered), low-profiled, with rounded or flattened bottoms, decorated with comb, cord, and other stamps, lines, forming bands of simple compositions on the upper vessel part, and incisions (including stamping) on the rim.

Burial grounds until the 7th century comprised mounds (sometimes with up to 12 burials) and adjacent (including synchronous) ground burials (with rows discernible), typically featuring supine body placements. Wooden structures and charcoal finds were common; animal bones and pottery appeared in fills (mostly in cremation burials). Burial items included attire details, usually worn, ceramic vessels, occasionally weapons, horse gear, and household items; silver masks near the eyes, silk face covers are known. Attire featured temple pendants, necklaces (beads, coins, etc.), bracelets, rings, belts, often of local Ural types, diverse pendants (numerous jingling ones) and awls. Hoards were numerous, including Iranian and Byzantine metalware, coins, jewelry; a special group featured Permian animal-style artifacts. Economy was based on slash-and-burn agriculture, animal husbandry, with hunting and other crafts being important.

Several local groups and 4 phases are distinguished: Kharino (6th century), Agafonovo (late 6th - 7th centuries), Demenki (late 7th - 8th centuries), Uryino (late 8th - 9th centuries), Lavryata (10th - late 11th centuries). The formation of the Lomovatovo tradition is associated with the colonization of the taiga and subtaiga of the Kama area by the Kharino-Glyadenovo population. The Medieval Rodanovo culture derives from the Lomovatovo tradition.

##### I.C.5.1. Bayanovo

The site was discovered in the spring of 1951 during quarrying activities for the construction of the Levshino-Kizel railway, which partially destroyed the monument. Soldiers from the military unit constructing the road collected some items and reported the findings to the Molotov (Perm) Regional Local Lore Museum, and later to the Molotov State University.

On July 18, 1951, students V.A. Oborin and Z.P. Sokolova visited the site of the finds to survey and collect items. An excavation was initiated at the edge of the quarry in its southeast part. In 1951 and 1953, excavations were conducted at the burial ground by V.A. Oborin. In August 2005, excavations at the burial ground were commenced by A.V. Danich (Danich, 2005). By 2012, a total of 283 burials had been studied across an area of 1472 m2.

###### burial 277 (individual ID I19113, Male)

The burial is oriented along the northeast-southwest axis and was identified at a depth of -0.84 to -1.10 meters from the arbitrary excavation datum. Its dimensions are 1.91 x 0.73 meters, with a maximum depth of -1.29 meters from the excavation datum. The bottom is flat, and the walls are vertical.

The burial contains the remains of a single individual arranged anatomically. In the northern part of the burial, a poorly preserved skull and mandibula are found. The skull lies with its facial aspect upward, while the mandibula stands on its base, oriented vestibularly to the south. The remains include the temporal bones, diaphyses of both forearms, metacarpal bones, and phalanges, as well as fragments of the pelvis and both femurs. The pelvis remains unopened. The long bones are aligned parallel to each other and to the burial's arbitrary axis.

The skeletal remains in the burial belong to a male aged 18-25 years. Based on the position of the bones, the deceased was placed on his back with his head to the northeast.

The burial yielded rich inventory:

1. Two solid bronze beads with a slight widening in the middle and an unornamented edge.
2. Fragments of a silver mask.

3. Fragments of a silver mask.
4. Silver wire temple ring of oval shape with a slight thickening at one end of the wire.
5. Silver pendant depicting a horse rider. The pendant hung around the neck using a bronze spiral-wound pin.
6. Fragment of a solid bronze bead with a slight widening in the middle and an unornamented edge.
7. Flint.
8. Pouch made from the skin of a large animal. Trimmed along the edge with small bronze ferrules. Suspended from the belt using a bronze spiral-wound pin and 6 solid bronze beads with a slight widening in the middle and an unornamented edge.
9. Round wire bronze bracelet, hexagonal in section. Ornamented on the outer side with three rows of circular ornamentation. Located on the left arm.
10. Round wire bronze bracelet, hexagonal in section. Ornamented on the outer side with three rows of circular ornamentation. Located on the left arm.
11. Bronze tube-shaped pipe tamper.
12. Fragments of a leather belt with bronze overlays. The front part of the belt is decorated with small oval overlays adorned with two groups of horizontal lines, with 3 lines per group. The side and rear parts of the belt are decorated with flat figurative overlays with protrusions on the lateral sides.
13. Flint with a bronze handle. The handle is carved with an image of two horse riders riding in different directions. Along the base, the flint is decorated with an ornamental band resembling a wavy line.
14. Solid bronze bead with a slight widening in the middle and an unornamented edge.
15. Knife with a wooden handle.
16. Flat figurative bronze overlay with protrusions on the lateral sides.
17. Flat figurative bronze overlay with protrusions on the lateral sides.
18. Laurel-leaf arrowhead without a socket.
19. Two small solid bronze beads with a slight widening in the middle and an unornamented edge.
20. Bronze openwork buckle depicting a wolf's head, with a rectangular ring and an oval receiver. The buckle is decorated with notches along the edges and the front edge of the ring.
21. 28-31,33. Four arrowheads without sockets.
22. Bronze shield-shaped buckle with an oval ring and a sharpened unornamented rear plate.
23. Coin die with a narrow, elongated, triangular blade and a hammer-like butt. The cross-section of the butt is rectangular.
24. Saber. The length of the preserved portion is 70.3 cm, the length of the preserved double-edged portion is 11.8 cm, the length of the handle is 10 cm, the width of the blade at the hilt is 3.6 cm, the width at the double-edged portion is 2.6 cm, the width of the handle is 2.2 cm, the curvature of the blade is 0.03 cm, the thickness of the handle is 4 mm, and the thickness of the blade is 5 mm. The handle has wooden spherical grips and a slight tilt toward the blade. The saber was housed in wooden scabbards, traces of which are visible along the entire length of the blade. The saber has a block-shaped crossguard with spherical ends, made of two parts. At a distance of 17 cm from the crossguard, there is an iron suspension consisting of two parts, each 1 mm thick, riveted together with two rivets.

The burial is dated to 770-875 calCE (1213±22, DeA11172).

#### I.C.6. The Polom culture (MidKama_Polom)

The first excavations on the sites later attributed to the Polom culture, were conducted in the late 19th century, and their materials remain unpublished. A brief description of the cultural assemblage was provided in the late 1950s, while the key site, the Varni burial ground, has been under study since 1970.

The sites along riverbanks consist of 1-2 fortified settlements, burial grounds, and open settlements. Fortified settlements on promontories with steep slopes feature ramparts and ditches, sometimes in two lines. The houses are rectangular, timber-framed, with areas ranging from 24 to 85 square meters, with clay or sandy-gravel floors and hearths on clay bases with stones, with 1-2 hearths along the walls; a gabled roof is reconstructed. Within or near the dwellings are 1-3 pits lined with boards or bark. Among the outbuildings, timber stables are prominent. Traces of metal and bone processing have been found. The pottery (round-bottomed bowls) is handmade, with shell admixture in the clay; predominant ornamentation includes comb-stamped and geometric patterns, filled with incised lattice stamps, cord impressions.

Burial grounds are large and flat; graves are arranged in rows. In the early part of the burial ground near the village of Varni, there are mounds with circular trenches. The bodies are laid on their backs, in coffins or wrapped in bast or birch bark, often oriented with their heads to the west (early graves) or north (later ones). Clay or metal vessels were placed in the graves, sometimes with antler spoons; pieces of meat were placed at the feet; at a later stage, burial rituals involved the inclusion of horse parts (skull or jaw, leg bones). Male burials contain earrings, bracelets, rings, belts sets, axes, spears, swords, sabers, bows with arrows, chisels, planes, knives, and horse harness items; female burials—earrings, temple and chest pendants, necklaces of beads, bracelets, rings, belts with appliqués and pendants, knives with elaborately decorated sheaths, awls, whorls, spindles. Various rattling pendants, including zoomorphic ones, are common. Southern trade (?) connections of the Polom population are confirmed by finds of "eastern silver" hoards (coins, vessels from Iran and Central Asia). Subsistence activities include slash-and-burn agriculture and animal husbandry, with hunting, including fur trapping, and fishing being important.

The evolution of the Polom culture passed four phases: in the early (Gyrkeshur, 4th quarter of the 4th(?) – 5th-6th centuries; Varni, late 6th-7th centuries) the sites are concentrated in the upper reaches of the Cheptsa River, while in the late stages (Karavaless, late 7th-8th centuries; Mydlan'shay, late 8th-9th centuries) the middle course of the river with its tributaries was developed.

The cultural assemblage demonstrates influence from the Glyadenovo, Azelino, and (less significant) Mazunino traditions. The later evolution of the Polom tradition formed the Cheptsa culture, which spread over the same territory.

##### I.C.6.1. Varni

The Varni burial ground, dated from the 5th century to the first half of the 10th century, is located on the northern outskirts of the village of Varni and occupies a promontory-like part of the gentle southern slope of the high right bank of the river, transitioning into the main terrace of the Cheptsa River. Discovered by V.A. Semenov in 1970, the burial ground underwent systematic excavations in 1970-1973, 1984, and 1990. In 1991, Semenov's work was continued by N.I. Shutova, and in subsequent years, from 1994 to 1998, 2003, and 2006, archaeological research was led by Aleksander Ivanov. By the mid-2000s, an area of about 3303 square meters

had been excavated, and 689 burials had been studied. From 2021 to 2023, excavations at the burial ground were conducted by Tatiana Sabirova.
Here we publish two individuals from the cemetery.

###### burial 158 (individual ID I33979, Male)

The clear outlines of the burial pit, with a quadrangular shape and rounded corners, were observed at a depth of 40 cm. The grave, measuring 292 x 75 x 59 cm, was oriented along the south-north line. At all four corners of the grave, traces of stakes with a diameter of 5 cm were recorded, located 7-10 cm away from them. The profile of the northeast stake was square-shaped, but only for the first 15 cm from the end. The grave had an expanded northern end and very smooth, well-worked walls. At a depth of 100 cm, well-preserved skeletal remains were uncovered. The deceased was laid out on their back, with the head to the north, and the arms extended alongside the body. Traces of birch bedding were found at the bottom of the grave under the iron objects. To the right of the head, closer to the northwest corner of the grave, stood an iron cauldron assembled from several plates, with a massive iron handle. Beneath the cauldron lay a axe-dagger and a flint fragment. On both sides of the skull, near the temporal bones, lay temple pendants. Along the western wall, touching the right hand, lay a sabre. Only the bow holders of the scabbard were preserved. The handle retained traces of wood and the pommel. Near the left femur, lay a whetstone, and below the bones of the left hand, outside of the femur, a compact heap contained a gray-colored sharpening stone made of fine-grained sandstone, a seat in the form of a plate, and a tube for whetstone oil. At the feet of the deceased were the bones/ribs of a horse and fishing tackles.

###### burial 260 (individual ID I33978, Female)

Fairly distinct outlines of a burial pit with a quadrangular shape and strongly rounded corners near the southern wall were discovered at a depth of 70 cm. The burial pit, measuring 200 x 70 x 50 cm with a wider northern end, was oriented along the south-north line. The central part of the grave was overlapped by a grave robber's trench, which partially destroyed the outlines of the eastern wall. At a depth of 90 cm, a burial was uncovered, disturbed in the chest and waist area. The skull with the shoulder girdle, as well as the lower limbs together with the pelvis, remained undisturbed. Judging by their position, the deceased was buried lying on their back, with the head to the north. The position of the arms could not be determined, as the arm bones and part of the ribs were found in a mixed condition near the pelvic bones. Near the skull, to the right of it, stood a vessel, while ordinary twisted pins and a copper bead lay on the pelvic bones, and a knife was found between the thighs. Remnants of bark from a coffin were found between the shins, and occasional coal inclusions were noted in the fill.

#### I.C.7. The sites of the Karayakupovo horizon in the Cis-Urals (Cis-Urals_KH)

##### I.C.7.1. Karanayevo

The burial site is located one kilometer south of the village of Karanaevo in the Mechetlinsky district, three kilometers north of the village of Abdrashitovo in the Duvansky district of the Republic of Bashkortostan, on a high (253.6 m) plowed elevation called Barag, 750 meters to the left of the Ufa–Yekaterinburg highway. The burial mounds covered an area of no less than 10,000 square meters.
The site was discovered and first investigated by Niyaz Mazhitov in 1964 and 1966, with 18 burial mounds excavated, providing rich material on the history and culture of the nomads of the 10th-12th centuries. In 2000 archaeologists were informed about intensive looting of the

cemetery, and the rescue excavations of the site were organized. Despite the fact that the burial ground was previously recoerded as a kurgan site, long-term plowing of the surface led to the complete leveling of the burial mounds in the 2000s, and they were no longer visually identified on the surface. This circumstance was the main reason why excavations were carried out on it as on a flat burial ground. The new excavation block was located on the plan drawn up by Niyaz Mazhitov in 1964, and it was recognized that the new excavation covers the area over three mounds located west of those excavated by N.A. Mazhitov (mounds No. 8, 9, 10). Thus, in 2001, 1080 square meters were excavated at the Karanaevo burial ground and 12 burials were investigated, providing new data on the material culture and burial rites of the population of the Southern Urals in the 10th-12th centuries.

We sequenced four burials from mound 1 and one burial from mound 2.

mound 1

burial 6 (individual ID I25536, Male)

The determined radiocarbon age of the burial is 664-827 calAD (1270±30 BP, Poz-136206) (based on human bone).

mound 2

burial 3 (individual ID I25539, Male)

The determined radiocarbon age of the burial is 885-1016 calAD (1105±30 BP, Poz-136207) (based on human bone).

#### I.C.8. The Medieval Chiyalik culture in the Cis-Urals (Belaya_Chiyalik)

##### I.C.8.1. Gornovo

mound 2, burial 2 (individual ID I25548, Male)

The determined radiocarbon age of the burial is 1178 -1276 calAD (810±30 BP, Poz-136209) (based on human bone).

##### I.C.8.2. Nizhne-Khozyatovo

burial 1 (individual ID I25540, Male)

The determined radiocarbon age of the burial is 1281-1395 calAD (650±30 BP, Poz-136208) (based on human bone).

burial 10 (individual ID I25544, Male)

The determined radiocarbon age of the burial is 1225-1295 calAD (745±30 BP, Poz-136326) (based on human bone).

burial 13 (individual ID I25546, Female)

The determined radiocarbon age of the burial is 1226-1377 calAD (735±30 BP, Poz-136327) (based on human bone).

burial 2 (individual ID I25541, Female)

The determined radiocarbon age of the burial is 1300-1406 calAD (605±30 BP, Poz-136246) (based on human bone).

###### burial 6 (individual ID I25542, Male)

The determined radiocarbon age of the burial is 1280-1395 calAD (655±30 BP, Poz-136324) (based on human bone).

###### burial 9 (individual ID I25543, Male)

The determined radiocarbon age of the burial is 1274-1389 calAD (685±30 BP, Poz-136325) (based on human bone).

### I.D. The Trans-Urals and the Western Siberia

#### I.D.1. The Early Iron Age Sargatka culture (MidIrtysh_Sargatka)

##### I.D.1.1. Bogdanovo-2

The burial site is located in the Kormilovsky district of the Omsk region, near the Om' River, a tributary of the Irtysh River. It was investigated in 1977 by V.A. Mogilnikov. We involve into this study two individuals buried under mound 2, individual ID I30386, Female and individual ID I30387, Female. The central burial under that mound was completely looted, and around its grave pit, on the ancient ground surface, remains of three individuals, including an infant, were found. Based on an earring, a bronze cauldron, and iron chain reins, the burial mound is dated to the 2nd century BCE – 1st century CE. The archaeological context has not been published yet.

##### I.D.1.2. Borovyanka-18

###### mound 13, burial 2 (individual ID I10117, Female)

The determined radiocarbon age of the burial is 385-206 calBCE (2240±20 BP, PSUAMS-9091) (based on human bone).

###### mound 3, burial 1 (individual ID I10118, Female)

The determined radiocarbon age of the burial is 382-206 calBCE (2235±20 BP, PSUAMS-9122) (based on human bone).

#### I.D.2. The Migration period sites of the late phase of the Sargatka culture (MidIrtysh_LSargatka & Tobol_LSargatka)

##### I.D.2.1. Borovyanka-17

###### burial 46 (individual ID I6826, Male)

The determined radiocarbon age of the burial is 262-420 calCE (1680±20 BP, PSUAMS-4827).. (based on human bone).

##### I.D.2.2. Ipkul'

The Ipkul' burial site is located in the Nizhnetavdinsky District of the Tyumen Region, it is situated at the northern shore of Lake Ipkul. The site was discovered in 1971 by M.F. Kosarev and V.F. Starkov (1971), and it was studied in 1983 by L.N. Koryakova (1984; 1988). Subsequent research over two years by the expedition of the Ural State University excavated 7 mounds (1, 2, 3, 6, 14, 28, 39), with another 2 (27 and 44) mounds investigated by trenches (Koryakova, 1988; Koryakova, Morozov, Sukhanova, 1988).

In 2010–2011, I.Yu. Chikunova documented 36 mounds elongated in small groups in the latitudinal direction (Chikunova, 2017). The majority of the necropolis is located on an open, flood-prone clearing overgrown with meadow grass and small shrubs, while the northern part is in a mixed coniferous-deciduous forest. Some mound structures are almost entirely leveled, with a height not exceeding 0.2 m, identified only through instrumental surveying. New numbers have been assigned to the documented mound structures, except for those previously used by the Ural State University expedition. Excavations were conducted on mounds 4, 5, 7, 9, 13, 18, 19, 20, 22, 25, 29, 36, 40, 41 (12), 42 (2), 43 (3). Additionally, the raised areas 10 and 12 were investigated, revealing well-preserved piles of sand and construction debris.

The inventory from the burials and radiocarbon dating indicate that the necropolis dates back to the 2nd–4th centuries CE. The site marks the concluding phase of the Sargatka culture, characterized by elements originating from the taiga zone, which are evident in the Kashina culture and the Sarov phase of the Kulayka culture. These elements subsequently influenced the formation of the Bakalda culture, which flourished during the Migration Period and the early Middle Ages (4th–8th centuries CE).

##### mound 13, burial 1, ind. 1 (individual ID I33844, Male)

The burial in the pit of a rectangular shape was located at the center of the mound. It measured 2.1×0.9 meters with a depth of 0.2 meters and was oriented along the north-south axis. The deceased, a male aged 35-45 years, was laid with his head to the north. His hands were resting on the pelvic bones. Corroded fragments of an iron knife handle, 13 centimeters in length, with remnants of iron sheathing for leather or wooden scabbards, were found on the right side of the pelvic bones. On the left side, near the lower edge of the ribs, a bronze round-frame buckle with a rounded shield and a movable tongue was discovered.

Pit 1, similar in configuration and orientation to burial 1, was positioned parallel to it with a displacement to the southwest. The fill consisted of dark gray loam. In the center, there was an inverted large undecorated pot of Bakalda type without a neck, filled with charcoal. No other finds were made. The burial is dated to 200-400 CE based on the belt bucle.

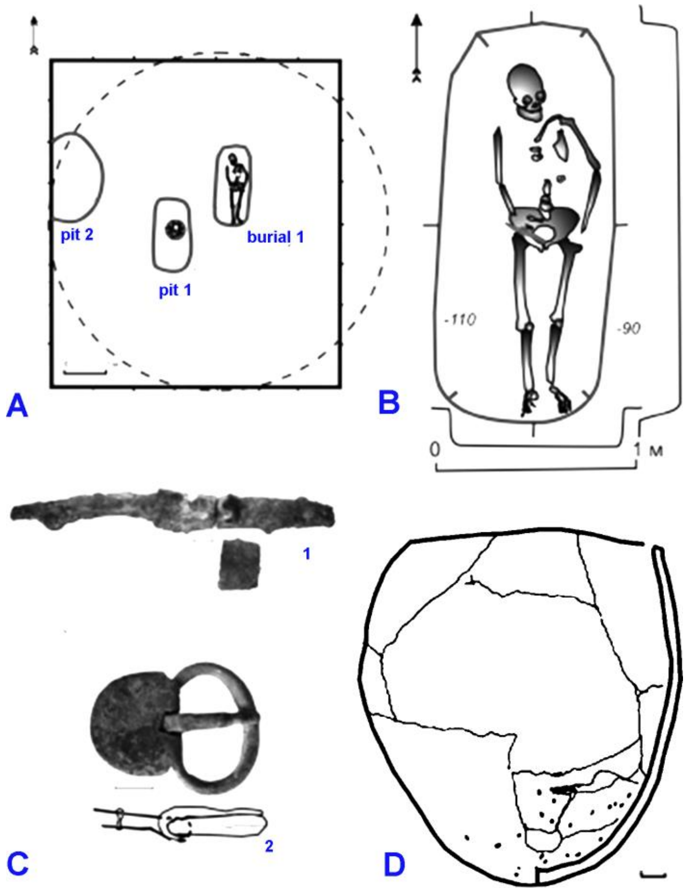

***Fig. SI.D.1.*** *Ipkul' burial site, mound 13, burial 1. A - a plan of the kurgan, B - a plan of the burial, C - grave goods, D - Bakalda-type pottery from the grave.*

##### I.D.2.3. Starolybaevo-4

The burial site is located approximately 4.5 km to the northwest from the village of Staro-Lybaevo, in the Zavodoukovsky district of the Tyumen region. The cemetery occupies an area on the elevated right bank of the Iset River, covered by mixed forest, adjacent to the vicinity of the Schetkovo lake. It was discovered by I. V. Zhilin in 1982 and surveyed in 1995.

A total of 41 mounds have been identified. Undisturbed mounds are situated along the forest edge, stretching in a northwest to southeast direction, well-preserved but showing signs of looting and badger holes. These include 18 mounds numbered 1-13, 16-18, 21, and 22. Additionally, six ploughed mounds were found along a sandy country road parallel to the aforementioned kurgans, within a distance of 15-30 meters, likely belong to the same group. Another group consisting of 15 mounds forms an irregular chain oriented from north to south, extending from the forest edge directly across the field to the Schetkovo lake.

Finds from mounds 31, 34, and 35 date back to the Neolithic-Chalcolithic period, representing remnants of short-term settlements along the shore of the now dried-up lake, to the edge of which the burial ground is presumably associated.

###### mound 34, NE sector 1, ind. 1 (individual ID I32779, Male)

Mound 34 had a diameter of approximately 15 meters and a height of 0.35 meters. The mound was regularly plowed, making it difficult to accurately determine its contours, resulting in the conditional center of the mound being slightly shifted to the southwest. Beneath the mound, a cultural layer from the Chalcolithic period was discovered.

Several post holes indicated the presence of some ancient above-ground structure. During excavations, it was found that the ditch surrounding the platform contained two main burials and had a polygonal shape, presumably hexagonal. The width of the ditch varied from 0.8 to 1.1 meters, with a depth of 0.25-0.35 meters. On the inner side of the ditch from the west, a lens of burning measuring 1.0x0.4 meters was identified. The dimensions of the platform outlined by the ditch were 11x13 meters. In the northwest sector, a collapsed Sargatka pot with scallops was found at a depth of 40 cm below the surface.

The main burials associated with the Early Iron Age were burials 1 and 2. All burials were looted.

Burial 2. The central grave in the mound, with a rectangular shape, had dimensions of the entrance pit, damaged by looters, measuring 4.25x3.0 meters. Debris lenses, created to the north and southeast, slid into the grave. During the excavation of the fill, the contours of the mound contracted and took on a more regular form. The chamber dimensions at a depth of 0.5 meters from the ground level were 2.5x3.2 meters. Particularly, on the northeast side, a flat platform was revealed, possibly serving as a step or resulting from an inlet burial cut into the northeast edge of the central mound. Supporting the latter hypothesis is the discovery of two intact and two damaged skulls in the grave, as well as the finding of articulated pelvic bones, femurs, radius, and ulna of the left arm belonging to a body that had not decomposed at the time of looting. This indicates multiple-stage burials in this chamber. The total depth of the grave was 0.86 meters, and the dimensions of the depression at the bottom were 2.0x1.5 meters. Due to looting, only fragments of pottery were found, the collapse of a small round-bottomed pot with notches along the shoulders, a spindle whorl, fragments of an iron sword blade, and an iron three-bladed arrowhead.

Anthropological analysis suggested the presence of six individuals: two adult women, as well as women approximately 45 years old, 40-50 years old, 30-55 years old, and around 60 years old. However, the genetic analysis confirmed male sex of the individual ID I32779, whose age was anthropologically determined as 40-50 years old.

##### I.D.3. The Nizhneobskaya culture (MidIrtysh_Nizhneobskaya & Ob_Nizhneobskaya)

The Nizhneobskaya culture was distributed across a vast area of the Western Siberian taiga, predominantly along the main river artery, the Ob River, and its tributaries (the Tobol, Ishim, and Irtysh Rivers). Its northern limits extended to the Arctic Ocean, while the southern boundaries reached as far as the northern edge of the forest-steppe. The distinct identification of this culture was accomplished by V.N. Chernytsov in the mid-20th century during fieldwork at the Us-Told settlement. The Nizhneobskaya culture existed from the 4th to the 13th centuries CE and evolved through several phases:

1. Karym, 4th–6th centuries CE
2. Zelenogorsk-Ryolka, 6th–8th centuries CE
3. Kuchiminskaya, 8th–9th centuries CE
4. Kintus, 9th–13th centuries CE

Karym phase of the Nizhneobskaya culture (4th–6th centuries CE) is described based on necropolises such as Kozlov-Mys, Krasnoyarsky-4, Alekseevka-50, and Alekseevka-51, characterizing the Migration period in Western Siberia. The population of these sites resided on the periphery of the forest-steppe region inhabited by the Bakalda archaeological culture. The carriers of Karym traditions engaged in communication and exchange with the Ural

population. This interaction is evidenced by the substantial quantity of imported items of Eastern European origin, such as belt sets, bronze bowls, and others.
It is reasonably inferred that the migration of Karym groups from the taiga zone to the south took place between the 3rd and 6th centuries CE. During this period, characteristic objects associated with these groups, such as pottery and ornaments, began to appear in the forest-steppe and subtaiga sites. However, this influx of population was not protracted and ceased by the 7th century CE. Moreover, this study indicates that there is no basis for positing substantial migrations of southern or eastern groups into the Lower Tobol River territory during the 3rd–6th centuries CE, as evidenced by archaeological materials (such as the emergence of a superstrate nomadic component).
Anthropological research on cranial series from the Bakalda and Karym cultural contexts in the Lower Tobol River region (burial sites Ipkul’, Revda-5, Kozlov-Mys, Pereyma, and Ustyug-1), dated to the 3rd–6th centuries CE, has revealed that these populations exhibited a mixed Eurasian-Mongoloid physical type. A portion of the group was identified as descendants of the Sargatka-associated population, displaying a predominance of Europoid features, while another part aligns with inhabitants of the Western Siberian taiga.

In this study, we included Karym phase Nizhneobskaya population samples from the Ust-Tara-7 burial site, located in the Omsk Trans-Irtysh Region. Regrettably, the cranial remains from the Ust-Tara-7 burial site, due to their poor preservation, were not studied by means of physical anthropology.
Kuchiminskaya Phase of the Nizhneobskaya Culture (8th–9th centuries CE). The period spanning the 8th to 9th centuries CE in the forested northern zone of Western Siberia is characterized by complexes attributed to the Kuchiminskaya phase of the Nizhneobskaya culture. During this phase, main cultural markers (housing construction, fortifications, and pottery production) of the Nizhneobskaya assemblage were standardized, and a large number of imported objects, including those of long-distance origin, indicated a sharp increase in trade and exchange networks. Notably, Iranian bronze vessels were widely distributed in the region (Chemyakin, Karacharov, 2002).
The typical traits of the Kuchiminskaya phase disappear rapidly in the latter third of the 9th century. During this period, magnificent fortifications were abandoned, pottery traditions underwent transformation, and there was an overwhelming dominance of imports from the west. These changes likely reflected dramatic political and economic transformations that occurred in the Cis-Urals and the Middle Volga regions, where the Volga Bulgaria was established.
The results of physical anthropological examination of the Kuchiminskaya phase population interred at Barsov Gorodok show its similarity to the modern Ugric-speaking population groups. However, comparative data from the middle taiga of Western Siberia for earlier medieval periods are currently lacking, complicating the identification of possible ancestral paleopopulations. It is worth noting that the series from the Barsovo burial site includes several individuals who, alongside the common characteristics shared by the entire group, exhibit certain Europoid traits that are clearly not of local origin. The closest similarities for this subseries were found in the Cis-Urals, particularly with series from the Mitino, Demyonki (6th–8th centuries CE), and Birsk-2 (3rd–7th centuries CE) burial sites. This may reflect a migration of a small population group from the Cis-Ural region into the middle taiga of Western Siberia sometime before 700 CE. However, our study examined not the migrants themselves but their descendants who intermixed with the local population. There are substantial arguments supporting the notion that this migration was a brief and momentary impulse, which did not

greatly impact the population of the middle taiga during the medieval period. A comparison between the Barsov Gorodok individuals and contemporary indigenous populations leaves little doubt that they are the direct ancestors of certain Eastern Khanty groups, particularly those along the Salym and Balyk Rivers (Poshekhonova, 2010).
In our study, the population attributed to the Kuchiminskaya phase of the Nizhneobskaya culture is represente by two inivivals from the Barsov Gorodok cemetery. Regrettably, they were not examined anthropologically due to their poor preservation.

##### I.D.3.1. Ust-Tara-7

The Ust-Tara-7 burial site is situated in the southern taiga zone of the Irtysh region, within the Tarsky District of the Omsk Oblast. Comprising eight burial mounds with heights of up to 0.5 meters, the site was excavated by I.E. Skandakov between 1990 and 1994, with the subsequent publication of materials from the burial ground in 1999. The necropolis is tentatively dated to the 4th–5th centuries CE.
Analysis of the material culture, anthropology, and burial customs indicates sustained interactions between the local population and the tribes of the forest-steppe of Western Siberia and the central Kazakhstan steppes. Notably, distinct cultural traits differentiate the Ust'-Tara assemblage from the taiga-based Nizhneobskaya culture. We identify this site as a specific Karym-type tradition and propose its evolutionary trajectory leading to the later Potchevash cultural complex.

###### mound 9, burial 1 (individual ID I19098, Female)

The burial was performed in an oval pit elongated from north to south, with a length and width of 3 x 1.9 m, respectively. The southern half of the pit has a depth of 0.72 m and a flat bottom, which gradually rises near the northern wall, forming a step-like feature approximately 0.25 m high. During the excavation of the fill in the northern part of the grave, fragments of charred wood were cleared, with the largest pieces measuring 24-36 cm in length, 8-12 cm in width, and 2-3 cm in thickness.
The skeleton of a female, approximately 20 years old, was positioned elongated on her back, with the head to the south. Due to the profiling of the pit's bottom, the lower part of the shinbones and the heels of the deceased were positioned at the edge of the step, and a clay vessel was placed near the end of the left tibia. Remains of charred vertical stakes, with a diameter of 4 cm, were also cleared around the feet. A bronze buckle with remnants of a leather belt was found near the left thigh, with a piece of charred birch bark lying on top of it. Among other findings, attention is drawn to a bronze necklace (?), consisting of four plates connected by overlapping iron rivets. To the left of the skull, which exhibited signs of artificial deformation, a wire temple pendant was discovered, while another pendant was found beneath the skull, where one end of the necklace was inserted. Small pieces of charcoal and charred birch bark were also present in this area.
The determined radiocarbon age of the burial is 124-311 calAD (1840±30 BP, Poz-136243) (based on human bone).

##### I.D.3.2. Barsov Gorodok

The burial site of Barsov Gorodok is located at a locality called "Barsova Gora" in the Khanty-Mansi Autonomous Okrug - Yugra, in the vicinity of Surgut city. Geographically, it is situated in the Surgut Lowland, the northern part of the Middle Ob Lowland, which is located in the central part of the West Siberian Plain. The area of Barsova Gora, where the burial site is located, represents a section of a high-rooted terrace that borders the right-bank floodplain of the Ob River. The height of the terrace in this area reaches 40 meters, which explains its name "gora" (mountain).

The predominant natural vegetation consists of pine with some Siberian pine and birch, while lichens and taiga grasses dominate the ground cover, along with various shrubs such as bog bilberry, lingonberry, and blueberry.

In the territory occupied by the Barsov Gorodok burial site, various groups of burials are referred to as different phases of the site. Barsov Gorodok-1 is the term for the Medieval graves, while burials from the Early Iron Age are attributed to Barsov Gorodok-3. The Medieval burials (Barsov Gorodok-1) date back to no earlier than 675-700 CE, while the latest ones are no younger than 1300 CE.

Burials 235 and 236 were excavated in 1989 by an archaeological expedition of the Ural State University. Both burials had been looted in the past, but they were positioned inside the group of graves that date to the 8th-9th centuries CE. The dating of the preserved artifacts from burials 235 and 236 does not contradict the overall dating of the group.

###### burial 235 (individual ID I19095, Male)

The burial pit, oriented along the west-east axis, had a subrectangular shape with rounded corners and a depth of 11-12 cm from the ground level. The western end of the pit was tapered. The width of the burial pit was approximately 50 cm, and the length was approximately 230 cm. The skeletal remains of a man, aged 20-25 years, were positioned with the skull facing the east on the dorsal surface, in such a way that there was some distance between the skull and the foot bones and the western and eastern walls of the pit (20 and 50 cm, respectively). The limb bones were aligned along the long axis of the pit, with the hands resting on the bottom of the grave next to large spits. Based on their position, the body was laid in the grave without any fixation. The skull lay with its facial surface upward, resting on the occipital and parietal bones. The position of the mandibula is unknown. The available skeletal remains were in anatomical order. At the time of excavation, the shin and foot bones, the right pelvic bone, and all the bones of the right upper limb were missing. The bones of the left arm were represented by decay, except for the distal ends of the radius and ulna, as well as the metacarpal and wrist bones. No traces of the burial construction were preserved. Between the skull and the eastern wall of the pit, a silver plate with protuberances was cleared, and remnants of an iron object (possibly a chisel) were found beneath the skull. Around the skull and near the left knee joint, eight silver appliqués were discovered. It can be assumed that these silver items have a Cis-Uralic origin.

The determined radiocarbon age of the burial is 131-336 calAD (1810±30 BP, Poz-136239) (based on human bone). Due to archaeological reasons, the date is irrelevant and likely biased by the freshwater reservoir effect.

###### burial 236 (individual ID I19096, Female)

The burial pit was oriented along the northeast-southwest axis and had a subrectangular shape with rounded corners, with a depth of 10-11 cm from the ground level. The southwest end of the pit was tapered. The width of the burial pit was approximately 38 cm, and the length was approximately 190 cm, nearly matching the height of the interred individual. The skeletal remains of a woman, aged 45-50 years, were positioned with the skull facing the northeast. The limb bones were initially aligned along the long axis of the pit. Based on their position, the body was laid in the grave without any fixation. However, during the looting of the burial, the left femur became dislocated, and the other leg bones were damaged. Only the shoulder bones of the arms were preserved. The skull, rotated to the left, rested on the base and basal surface

of the mandibula. The remaining skeletal remains were in anatomical order, except for the left femur. No traces of the burial construction were preserved. Near the left half of the mandibula, a small ceramic vessel was cleared, and a large iron knife was placed southwest of the non-preserved bones of the right forearm. A silver appliqué, similar to the previous one, was found in the abdominal area of the interred individual. A fragment of pottery was discovered in the vicinity of the non-preserved foot bones.
The determined radiocarbon age of the burial is 261-535 calAD (1655±30 BP, Poz-136240) (based on human bone).

#### I.D.4. The Potschevash culture (MidIrtysh_Potchevash)

The sites attributed to the Potchevash culture are primarily distributed in the southern taiga and forest-steppe regions along the valleys of the Ishim and Irtysh rivers, encompassing the Tyumen, Omsk, and Novosibirsk regions. The identification of this culture can be credited to V.N. Moshinsky, who based his classification on the assemblage of sites located at Chuvashsky Mys, near the city of Tobolsk, during the mid-20th century. Dated to the 6th-8th centuries CE, the Potchevash culture exhibits a heterogeneous character, incorporating elements from the Kulay, Sargatka, Nizhneobskaya, and Bakalda cultures, as well as early Turkic traditions, reflecting the cultural admixture of its population.

##### I.D.4.1. Vikulovo

The burial ground of Vikulovo Cemetery is situated in the western part of a ridge-shaped outcrop that extends from west to east on the terrace of the Ishim River. The cemetery was excavated by Viktoria Ilyushina in 2008.

###### collective burial 1

The burial was situated in the eastern sector of the excavation area. It consisted of a sub-rectangular grave pit with rounded corners, discovered at a depth of approximately 0.6 m from the present-day surface. The vault of the skull and clavicle were located near the northern wall of the grave, while the tibia was found near the southern wall. The grave itself measured 1.75 × 2.0 m in size, with a depth ranging from 0.04 to 0.08 m from the mainland level. Its orientation followed a north to south axis. Three pits were observed on the surface of the burial fill, likely the result of later disturbances unrelated to the original burial event. Within the grave, the remains of an estimated four individuals were discovered. The cultural layer of the archaeological site primarily consisted of loam; however, the condition of the bones was poor, likely due to disturbance caused by subsequent construction activities in the Russian period.
Skeleton 1 (individual ID I19094, Male) (referred to as "skeleton 2" in the publication) was located near the western wall of the burial. The skeletal remains belonged to an adult male, aged approximately 35-40 years at the time of death. It is presumed that the individual was originally placed in a supine position, with the head oriented towards the north. The mandibula was damaged, the skull fragmented, and various skeletal elements, including cervical vertebrae, collarbones, ulna, humerus, radius, femur, and a fragment of the pelvic bone, were preserved. However, due to disturbances, it is not possible to determine the original body position during burial (Ilyushina, 2009). Of particular interest is a significant chop wound observed on the parietal and occipital bones of the male individual, suggesting a potential warrior status (unpublished data from the author).
The determined radiocarbon age of the burial is 27-213 calAD (1915±30 BP, Poz-136238) (based on human bone). The radiocarbon date contradicts artefactual data and likely was bias due to freshwater reservoir effect.

#### I.D.5. The sites of the Karayakupovo horizon in the Trans-Urals (Trans-Urals_KH)

##### I.D.5.1. Uyelgi

The Uelgi kurgans are located in the Southern Urals, at the junction of the forest-steppe and steppe natural zones, within the modern Kunashaksky District of the Chelyabinsk Oblast, Russia. The site is situated on one of the terraces between the Uelgi and Saigerly lakes. Most of the burials under the mounds were conducted using the inhumation rite, with cremations being rare.

The cultural and chronological attribution of the site is based on a group of distinctive artifacts, which include:

1. Plaques of the Bekeshev and Husainov types (9th century),
2. Belt sets of the Srostki type (second half of the 9th century – first half of the 11th century),
3. Horse harnesses and weapons dating from the late 8th to the 11th centuries,
4. Potter ofy the Kushnarenkovo, Karayakupovo, and Petrogrom types.

The chronological attribution is confirmed by a dozen radiocarbon dates. Based on the combined archaeological and C14 data, the site is divided into two chronological phases:

1. 9th century,
2. Late 9th – early 11th centuries.

The cultural influence of the Srostki assemblage, evidenced at Uelgi by changes in burial practices and the appearance of gilded, ornamented overlaid plaques, raises the question of a possible infiltration of a population of Altai origin into the Southern Trans-Urals during the Karayakupovo horizon period.

###### mound 7 (33), burial 5 (individual ID I19121, Male)

Burial mound 7 (33) was excavated in 2011. Burial 5 features a sub-rectangular pit with a depth of 0.15 m, with remnants of a wooden coffin around the perimeter of the burial chamber. The pit contained one individual, laid out supine. Discovered artifacts include hemispherical bridle plaques, belt plaques with gilding and plant motifs characteristic of the Srostki culture, silver brackets for the front arch of a saddle, and an earring. The pit had been looted, preserving only fragments of long bones and the skull, making it impossible to determine the age of the interred. Based on the archaeological context, the burial dates from the late 9th to the first half of the 11th century CE. Radiocarbon analysis provided a combined date of 879-1150 calCE (1053±50, SPB_854; 888-1151 cal AD, 1040±50, SPb_852, 95.4% probability). The burial is attributed to the Srostki group.

###### mound 9, burial 5 (individual IDs I19115, Male and I19116, Male)

The grave pit of this burial has been heavily damaged due to looting, making it difficult to determine the original elements precisely. The depth of the pit was 0.25 m, and its shape is deformed; bones are scattered and many are lost, presumably belonging to two individuals, likely laid out supine. The presence of wooden logs or coffins is not established. Among the artifacts found are a silver belt tip and pottery of the Petrogrom type. According to the archaeological context, the burial dates from the 9th to 10th centuries, although the grounds for cultural attribution are not determined.

###### mound 28, burial 5 (individual ID I19114, Female)

Mound 28 was excavated in 2011. Under the burial mound, consists of a pit with a depth of 0.25 m, originally sub-rectangular in shape. The pit's shape was disturbed in the western part and oriented along the WSW–ENE line. The pit contained one individual, laid out supine with the head to the WSW. No burial structures were found. Among the artifacts discovered was a silver plate. The age of the interred could not be determined, but it is presumed to be an adult. Based on the archaeological context, the burial dates from the 9th to 10th centuries. Cultural attribution suggests a possible connection to the Srostki-influenced group of the Uelgi population.

###### mound 29, burials 1 and 2 (disturbed) (individual ID I19118, Male)

In the central part of the mound, two pits were discovered, deformed and of an amorphous shape. Both pits contain remnants of a wooden structure along the inner perimeter. Judging by the size and proportions of the pits, it was assumed that each pit contained no fewer than 2–3 individuals, however, the pits were heavily looted, bones scattered, so they are combined into one complex. Fragments of the skull were found in the western part, indicating a western orientation of the bodies, likely laid out supine. Among the artifacts found were silver buckles, buttons, overlays, and remnants of a quiver belt. According to the archaeological context, the burial dates to the second half of the 9th century, and radiocarbon analysis provided a calibrated age of 774–883 AD (95.4% probability). The burial is attributed to the Srostki-influenced group.

###### mound 30, embankment (individual ID I19119, Male and individual ID I19120, Male)

Mound 30, excavated in 2011, on the embankment (above ground level), a skull and a gilded terminal belt overlay of the Srostki type were discovered. The pit, likely associated with the discarded skull, was looted, heavily deformed, and empty. No burial structures were found. There were no significant findings among the artifacts in this mound. No deformations of the skull or other features were identified. According to the archaeological context, the burial dates to the second half of the 9th to 10th centuries. Based on the artifacts found, the burial likely belongs to the Srostki-influenced group.

###### mound 32, burial 12 (individual ID I19117, Male)

No burial structures were found. The depth of the pit is 0.35 m, its shape is sub-triangular, deformed due to looting. The pit contained one individual, likely laid out supine with the head to the west. Among the artifacts found were overlays of the Husainovo and Bekeshev types (9th century). The age of the interred could not be determined, but it is presumed to be an adult. According to the archaeological context, the burial date is from the 9th century, possibly the first half of the 9th century. The burial is likely associated with the group of nomads from the 9th century associated with Bekeshev cultural tradition. The determined radiocarbon age of the burial is 771-937 cal AD (1184±22 BP, DeA_11458) (based on human bone), which confirms archaeological observations.

##### I.D.6. The Medieval Ust'-Ishim culture (MidIrtysh_Ust-Ishim)

In the 1950s, V.N. Chernetsov attributed the Medieval sites of the Lower Ob region and the southern taiga of the Omsk part of the Irtysh River to the Kintus phase of the Nizhneobskaya culture. However, further investigations conducted in the Omsk part of the Irtysh River led V.A. Mogilnikov and B.A. Konikov to identify a distinct cultural tradition in the region, dated

to the 9th-13th centuries CE. B.A. Konikov demonstrated that the Medieval people of the Irtysh Basin Southern Taiga constructed kurgan burials and based their economy on cattle breeding, in contrast to the fishers attributed to the Nizhneobskaya culture, who buried their dead in flat burial grounds.
The sites of the Ust-Ishim culture are primarily located in the sub-taiga and southern taiga zones of Western Siberia, from the forest-steppe boundary in the south to the Vasyugan River in the north, and from the Tobol River in the west to the Tara River, a tributary of the Irtysh, in the east. The chronological frame of the Ust-Ishim Medieval tradition spans from 800 to 1240 CE. This relative chronology is supported by the presence of imported ornaments from Volga Bulgaria, Old Rus, and the Medieval Kama region. The Ust-Ishim culture is believed to be associated with the ancestors of the southern Khanty, who had close contact with the Turkic-speaking population, likely the Srostki culture population. However, physical anthropological examinations suggest that the Siberian Tatars are the most probable descendants of the Ust-Ishim groups.
The total collection of Ust-Ishim anthropological material includes 46 male and 22 female skulls from Ivanov Mys-1 and Panovo-1, as well as additional samples from the burial grounds of Masarly-1, Nugai-1, and Dolgovsky-1 settlement. The series appears homogeneous and likely represents a single paleopopulation, resembling the groups buried in the forest-steppe variety of the Kulai cultural context and the West Siberian Mongoloids from the Sargatka tradition burials. Among the modern ethnic groups of Western Siberia, the Ust-Ishim people are closest in anthropological type to the Tobol-Irtysh Tatars, particularly the Tyumen and Kourdak-Sargatka groups, and are considered their direct ancestors.
A slight variability of features within the series of skulls from Ivanov Mys-1 and Panovo-1 possibly indicates the presence of a small immigrant population and their descendants. The closest similarities for this group were found in various sites associated with the population attributed to the Srostki and Basandaika cultures (Sanatorny-1 burial ground, 1000-1250 CE; Gilevo burial ground, 700-1000 CE, and others). The influx of migrants from South Siberia to the territory of the Irtysh Basin Southern Taiga ceased in the 14th-16th centuries (Poshekhonova, 2011).

##### I.D.6.1. Ivanov Mys-1

The Ivanov Mys-1 burial mound is located near the village of Ivanov Mys, Tevriz District, Omsk region, 300 km north of Omsk, and consists of more than 50 burial mounds up to 14 m in diameter and 1 m high.The burial ground dates back to the 13th century CE. We sequenced 3 individuals from under mound 10, one - from under mound 12, and one individual from unidentified burial (individual ID I19091, Male). This last individual was dated by C14, and its age was determined as 416-545 calAD (1600±30 BP, Poz-136215), which is likely biased by freshwater reservoir effect.

###### mound 10, burial 3 (individual ID I19087, Male)

The burial was located in the north-east part of the mound, in a rectangular pit oriented northwest-south-east. The size of the grave was 2.15 x 1.25 m, the walls were steep, and the corners were smoothly rounded. At the bottom of the grave rested two dead people stretched out on their backs. Both bodies were oriented with their heads to the northwest. In the southeast corners, two clay vessels stood upside down, one of which was placed on the grave slab and the other on the bottom of the pit. Fragments of wooden planks and birch bark cloth were preserved on the skeletons. On the man's head there was a bronze half-tambourine, and on the sides there was one lapis lazuli figurative pendant, on the inner side of the left radial bone of the skeleton there was a single-bladed iron knife. Between the skulls there was a bronze chime. The determined radiocarbon age of the burial is 1165-1267 calAD (835±30 BP, Poz-136210) (based on human bone).

mound 10, burial 6 (individual ID I19088, Male)

The disturbed burial was found in the western part of the mound. The skull was lying with the parietal bone to the north-north-west facial bones upwards. To the west of it, on its side, with its mouth to the north-west, was a clay bowl-shaped vessel, under which a bone petiole arrowhead lay.
The determined radiocarbon age of the burial is 250-411 calAD (1720±30 BP, Poz-136211) (based on human bone). Due to archaeological reasons, the date is irrelevant and likely biased by the freshwater reservoir effect.

mound 10, burial 9 (individual ID I19089, Male)

The determined radiocarbon age of the burial is 356-57 calBCE (2160±30 BP Poz-136212) (based on human bone). Due to archaeological reasons, the date is irrelevant and likely biased by the freshwater reservoir effect.

mound 12, burial 1 (individual ID I19090, Female)

The burial was located in a central rectangular continental pit, oriented northwest-south-east. It was robbed. The walls are steep, the corners are rounded, and the bottom is flat. The skeleton is disturbed. The deceased was stretched out on his back with his head to the northwest. An iron flat petiole arrowhead lay to the corner of the skull. Between the tibiae was an oval wooden object 7 cm in diameter. A broken bone ornamented plate lay parallel to the tibia. In the southern part of the grave fragments of plaques were discovered. A single-bladed knife in a wooden scabbard, a bronze lunette, and bone ornamented plates were placed in the grave.

I.D.6.2. Panovo

The Panovo-1 burial mound is located 4.5 km southwest of the Panovo village, Ust-Ishim region, Omsk Region, on the first supra-flood terrace of the right bank of the Irtysh River. It consists of 77 embankments up to 20 m in diameter and 1.2 m high.

burial 1 (individual ID I19097, Female)

The determined radiocarbon age of the burial is 210-353 calAD (1785±30 BP, Poz-136241) (based on human bone). Due to archaeological reasons, the date is irrelevant and likely biased by the freshwater reservoir effect.

mound 1, burial 30 (individual ID I19092, Female)

The determined radiocarbon age of the burial is 641-775 calAD (1350±30 BP, Poz-136236) (based on human bone). Due to archaeological reasons, the date is irrelevant and likely biased by the freshwater reservoir effect.

mound 67, burial 3 (individual ID I19093, Male)

The determined radiocarbon age of the burial is 647-775 calAD (1335±30 BP, Poz-136237) (based on human bone). Due to archaeological reasons, the date is irrelevant and likely biased by the freshwater reservoir effect.

##### I.D.7. The Late Medieval and Early Modern burials (Tobol_Tatars & MidIrtysh_Modern)

###### I.D.7.1. Putilovo

unidentified burial (individual ID I10116, Male)

The determined radiocarbon age of the burial is 1290-1397 calCE (635±25 BP, PSUAMS-9121) (based on human bone).

### I.E. The Carpathian Basin

#### I.E.1. The Conquest period sites

The Hungarian Conquest Period culture, first identified in 1834, has been extensively studied by scholars such as Fettich Nándor, László Gyula, István Fodor, Csanád Bálint, László Révész, and István Erdélyi. This archaeological heritage of the Old Hungarians is primarily located in the Carpathian Basin, with significant presence along the Subbotsi type sites. With approximately 40,000 graves dating from the late 9th century to the early 12th century, the culture is characterized by inhumation in normal gravepits, typically containing male individuals with classic partial horse burials. Key artefacts such as archery equipment and horse harnesses serve as chronological indicators. The culture, primarily associated with pastoralism, exhibits no significant dietary indicators affecting radiocarbon dating or reflecting migrations. While written sources and chronology suggest an ethnic attribution to the Old Hungarian people, archaeological genetic studies point to an eastern origin, with nine graves showing no biological connections. The conventional periodization places the culture in its early phase during the first half of the 10th century, with distinct variations observed in Transdanubia and the Great Hungarian Plain.

###### I.E.1.1. Balatonújlak

The site, discovered by Zsuzsanna Siklósi in 2003, is located on the north-northeastern edge of the Marcali hill, facing Lake Balaton. This site includes 26 graves arranged in a W-E orientation. No radiocarbon data is available, but chronological indicators such as saddles, harnesses, lozenge-shaped mounts, and caftan mounts date the grave to the mid-10th century. The cemetery represents a fully excavated site with inhumation burials, characteristic horse harnesses, and other specific artifacts. There is no evidence of cultural admixture, and the main subsistence activity inferred is pastoralism. The cemetery is attributed to the early Hungarian culture based on archaeological evidence, though no specific historical or linguistic data is provided.

grave 10 (individual ID I19075, Female)

Grave 10, described by Attila Türk and László Révész, is part of an Early Hungarian cemetery from the 10th century in the Carpathian Basin. Burial 10 is oriented west-east and belongs to a burial row. The grave is a rectangular pit without any constructions such as stone slabs or timber coverings. It contained a stretched skeleton laid on its back with no traces of a coffin. Artifacts found in the grave include characteristic horse harnesses, pear-shaped stirrups, lozenge-shaped mounts, and caftan mounts. No food or dye was discovered. The grave contained the skeleton of an adult woman and an infant, but their exact ages are unknown. There is no information about cranial deformation, odontological details, or dietary habits that might affect radiocarbon results. The mounts date the burial to the mid-10th century, though no C14 dates are provided. Cultural attribution details are not specified, but Western European coins found in the cemetery suggest significant cultural interactions.

burial 17 (individual ID I19076, Male)

Burial 17, excavated in the burial row with a W-E orientation, features a rectangular burial pit without any constructed elements. The grave contained a stretched skeleton of an adult woman laid on her back, with no trace of a coffin. The grave goods included a characteristic braid ornament and earrings with bead-row pendants. Based on the mounts found within, the burial is dated to the mid-10th century.

I.E.1.2. Harta

The Early Hungarian cemetery Harta from the 10th century Carpathian Basin, first described by Attila Türk and László Révész, is situated in the Danube-Tisza interfluve region of Bács-Kiskun County. This fully excavated site, discovered by Rozália Kustár and Péter Langó in 2002, comprises 21 graves arranged in three rows, located on a hillside. The cemetery, featuring inhumations with characteristic caftan mounts dating to the mid-10th century, demonstrates no signs of population admixture and includes typical pastoralist artifacts. The assemblage, particularly the richer female graves compared to poorer male graves, aligns with the Early Hungarian cultural practices, without invoking significant debate regarding its ethnic or linguistic identity.

burial 9 (individual ID I19077, Female)

Burial 9 is oriented W-E and lacks any visible grave marker or construction. The grave is a simple, rectangular pit with a depth of 7.5 cm and a skeleton length of 161 cm. The skeleton, a female estimated to have been 30-40 years old at death, was found lying on its back in a stretched position with the skull turned to the right and arms outstretched beside the body. No tree-trunks or coffins were present. The grave contained caftan mounts and textile remains, but no evidence of food. The burial, dated to the mid-10th century based on the artefactual evidence, specifically the caftan mounts, aligns with the Early Hungarian cultural practices. Despite being disturbed by animals, which displaced stones and finger bones, the remains were well-preserved. There is no cranial or odontological data, nor dietary details available. Paleopathological analysis indicated no battle injuries. The presence of small beads in the cemetery suggests the use of Byzantine luxury textiles, reinforcing the cultural attribution to the Early Hungarians.

burial 19 (individual ID I19070, Female)

Burial 19 with a grave depth of 32 cm, length of 250 cm, and width of 87.5 cm. The skeleton, a female estimated to have been 61-67 years old at death, measures 153 cm in length and is oriented W-E (280-100). The skeleton was found lying on its back in a stretched position with the skull turned to the left. The arms were placed next to the body, broken at the elbows, and bent over the left pelvic floor, with some phalanges displaced by animals. There were no traces of a coffin, although the position of the deceased does not exclude the possibility of one. The grave contained a silver ring but no evidence of food. Paleopathological analysis indicated no battle injuries, and there is no available cranial or odontological data, nor dietary details. Based on artefactual evidence, specifically the silver ring, the burial is dated to the mid-10th century. The cemetery is attributed to the Early Hungarian culture, with the presence of small beads suggesting the use of Byzantine luxury textiles, reflecting possible cultural interactions.

##### I.E.1.3. Szeged-Öthalom, V. homokbánya

The site, discovered during a rescue excavation by the Móra Ferenc Múzeu`m in 2009, is situated atop sand dunes, characteristic of the period. Referred to as a flat cemetery, it features inhumations without distinct rows, with graves positioned close to the sand dune tops. The graves typically contain single individuals in simple chamber gravepits. Radiocarbon dating from all graves suggests a chronological placement in the first third of the 10th century or the end of the 9th century, with some graves indicating a later usage around the middle of the 10th century. Artefactual evidence, including grave goods with parallels from Eastern Europe, indicates cultural connections, while the presence of west European coins from military campaigns to North Italy in the early 10th century suggests broader trade networks. Notably, some graves exhibit atypical features, such as an adult woman buried with a child 150 km away, suggesting rapid movement or shifting community dynamics. The site's overall phase spans the first half of the 10th century, with evidence pointing to possible changes in community usage over time (Türk et al 2015).

###### grave 36 (individual ID I19071, Male)

Grave 36 belongs to the flat cemetery and lacks any burial constructions. The grave is rectangular with rounded corners, oriented NNE-SE (335-155º), measuring 240 cm in length, 70 cm in width at the shoulders and feet, and 110 cm in depth. The buried individual, a male aged 40-45, was found in a supine, stretched position with the skull slightly tilted to the left side and the shoulders contracted. The arms were tightly against the ribs, while the legs were extended and brought together at the knees. Artefacts recovered from the grave include elements of horse harnesses, archery equipment, a silver hair ring, a finger ring, and some silver fragments. The presence of bones positioned higher up from the bottom of the pit suggests the use of a rounded tree coffin. No cranial deformation or odontological descriptions were provided, and no dietary details affecting radiocarbon results or reflecting migrations were identified. The burial has been culturally attributed to the classic Old Hungarian stage, with no apparent import influences. A radiocarbon date (Poz-42783) obtained from the burial falls within the range of 780-874 cal AD (68.2%) and 766-895 cal AD (87.7%) (Türk et al 2015).

###### grave 124 (individual ID I19072, Male)

The burial, identified as Szeged-Öthalom, V. homokbánya, Grave 124 is rectangular with rounded corners, oriented NE-SE (318-138º), measuring 235 cm in length and 90 cm in width at the shoulders, with a depth ranging from 7 to 34 cm. The grave contained the skeletal remains of a mature male aged 50-59 lying on his back in a stretched position, with mechanical damage to the trephined skull and disturbance on the right side of the skeleton from an animal passage. Artefacts recovered from the grave include elements of horse harnesses, archery equipment, flintstone, kresalo, two silver hair rings, sabretashes with bronze mounts, and an iron knife. Additionally, the upper leg of a sheep was found, possibly indicative of funeral food. The presence of bones positioned higher up from the bottom of the pit suggests the use of a rounded tree coffin. No cranial deformation or odontological descriptions were provided, and no dietary details affecting radiocarbon results or reflecting migrations were identified. A radiocarbon date (Poz-42782) obtained from the burial falls within the range of 780-877 cal AD (68.2%) and 768-897 cal AD (88.8%). The burial is culturally attributed to the classic Old Hungarian stage, with no apparent import influences in burial rites or artefacts (Türk et al 2015).

grave 132 (individual ID I19073, Male)

The burial, identified as Szeged-Öthalom, V. homokbánya, Grave 132 is rectangular with rounded corners, oriented NNW-SSE (304-124º), measuring 210 cm in length and 70 cm in width at the shoulders, with a depth of 20 cm. The grave contained the skeletal remains of a mature male aged 50-59 lying on his back in a normal position. Artefacts recovered from the grave include elements of horse harnesses, archery equipment, flintstone, kresalo, two silver fragments, sabretashes with bronze mounts, an iron knife, and a bronze tool for determining the size of belts. Additionally, the right part of a sheep shoulder was found, possibly indicative of funeral food. The presence of bones positioned higher up from the bottom of the pit suggests the use of a rounded tree coffin. No cranial deformation or odontological descriptions were provided, and no dietary details affecting radiocarbon results or reflecting migrations were identified. A radiocarbon date (Poz-42778) obtained from the burial falls within the range of 782-891 cal AD (68.2%) and 774-900 cal AD (83.1%). The burial is culturally attributed to the classic Old Hungarian stage, with no apparent import influences in burial rites or artefacts. A separate radiocarbon date (Poz-42793) obtained from a horse found in the burial suggests a date range of 872-967 cal AD (68.2%) and 805-973 cal AD (91.1%) (Türk et al 2015).

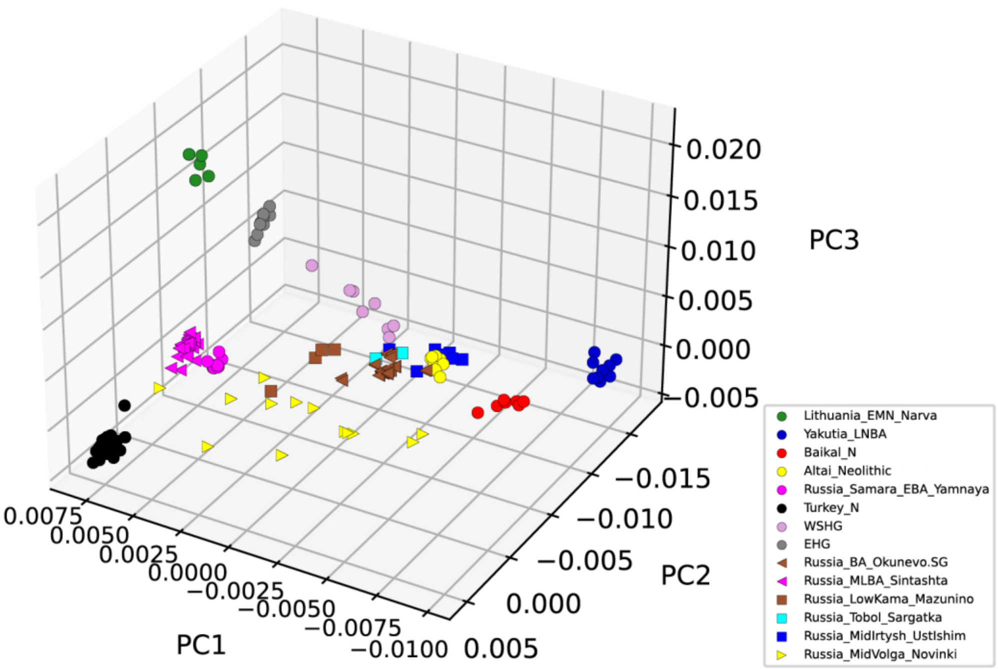

**Figure S1:** 3D PCA scatterplot (PC1/PC2/PC3) showing the Neolithic and Bronze Age proxy source groups used for the supervised *ADMIXTURE* analysis (shown as circles). Bronze Age reference groups used in Fig. 4A as references for the *f4*-statistics are also projected on the PCA plot (shown as left-facing triangles). From the newly sequenced groups, those which were used for the qpAdm analysis (Fig. 4B) are plotted in this figure. The Mid-Irtysh Ust-Ishim and Mid-Volga Novinki groups, which showed distinct ancestral components compared to the other Volga-Uralian groups, are also shown.

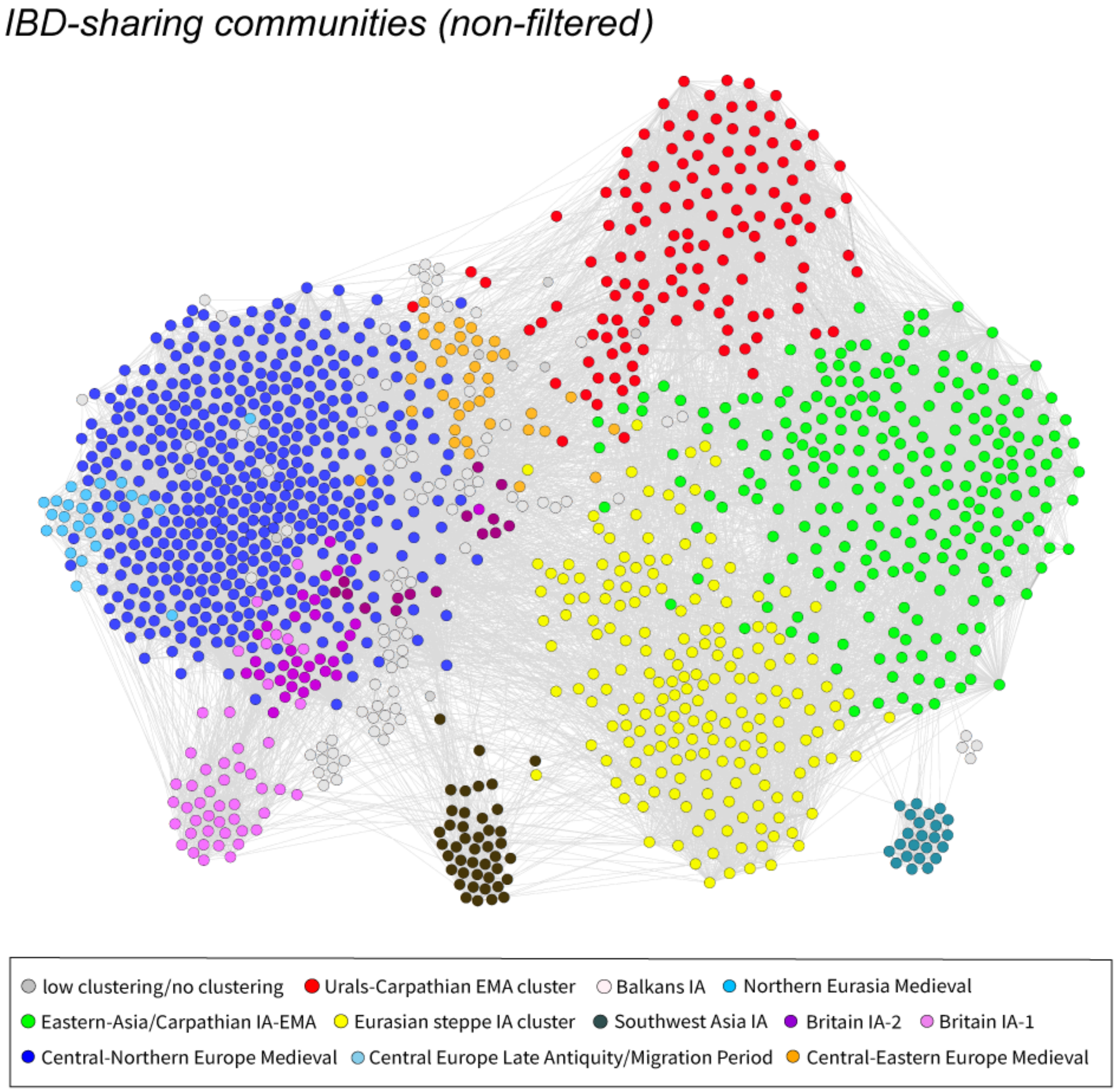

**Figure S2:** Non-filtered version of the IBD network presented in Figure 3. For filtering parameters see Methods.

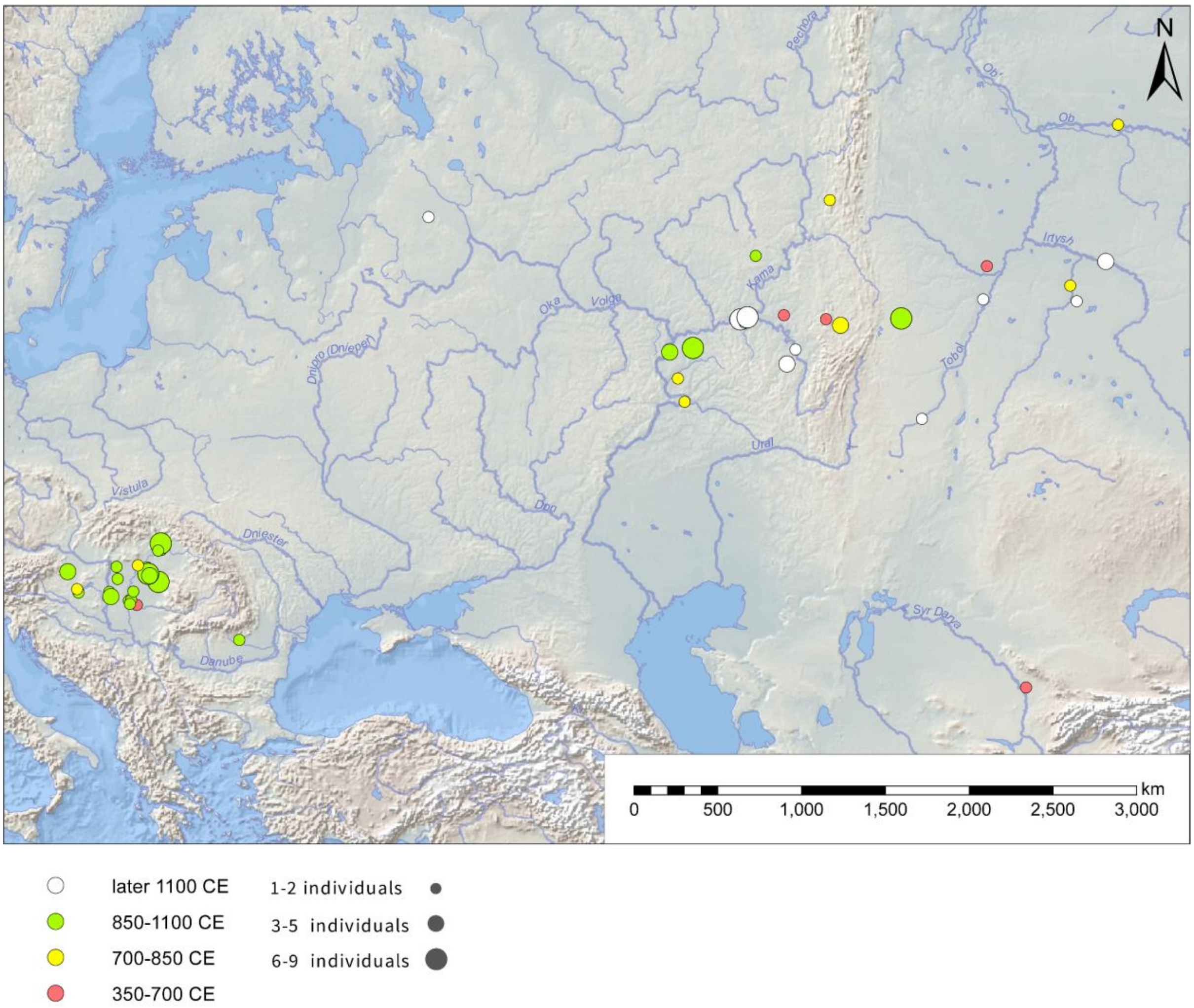

**Figure S3**: Geographical distribution of the individuals from the *Urals-Carpathian EMA* IBD-sharing community.

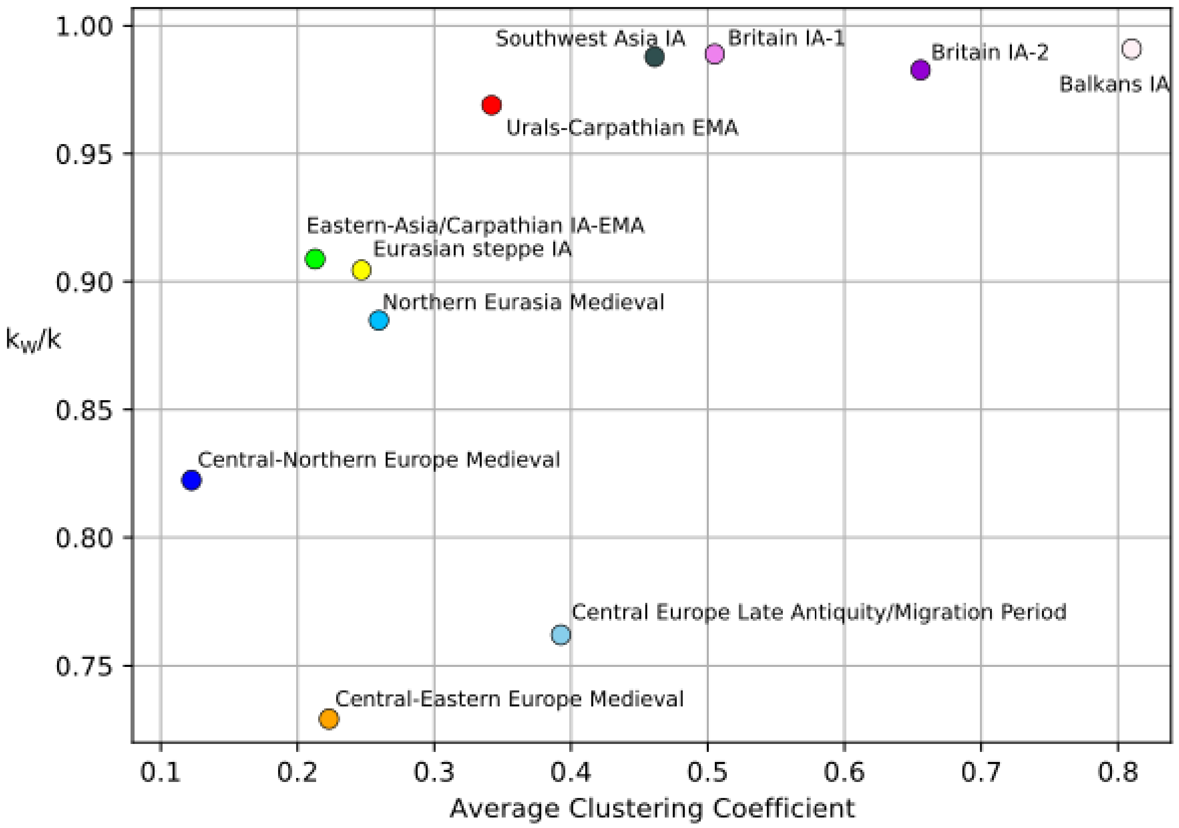

**Figure S4:** A scatterplot displaying the average clustering coefficient of each cluster versus the ratio of within-module connections ($k_w$) to the total degree centrality (k) of the cluster. Each cluster is identified using the Leiden algorithm, which determines community structure in the network. The 'modules' in this context refer to the clusters defined by this algorithm.

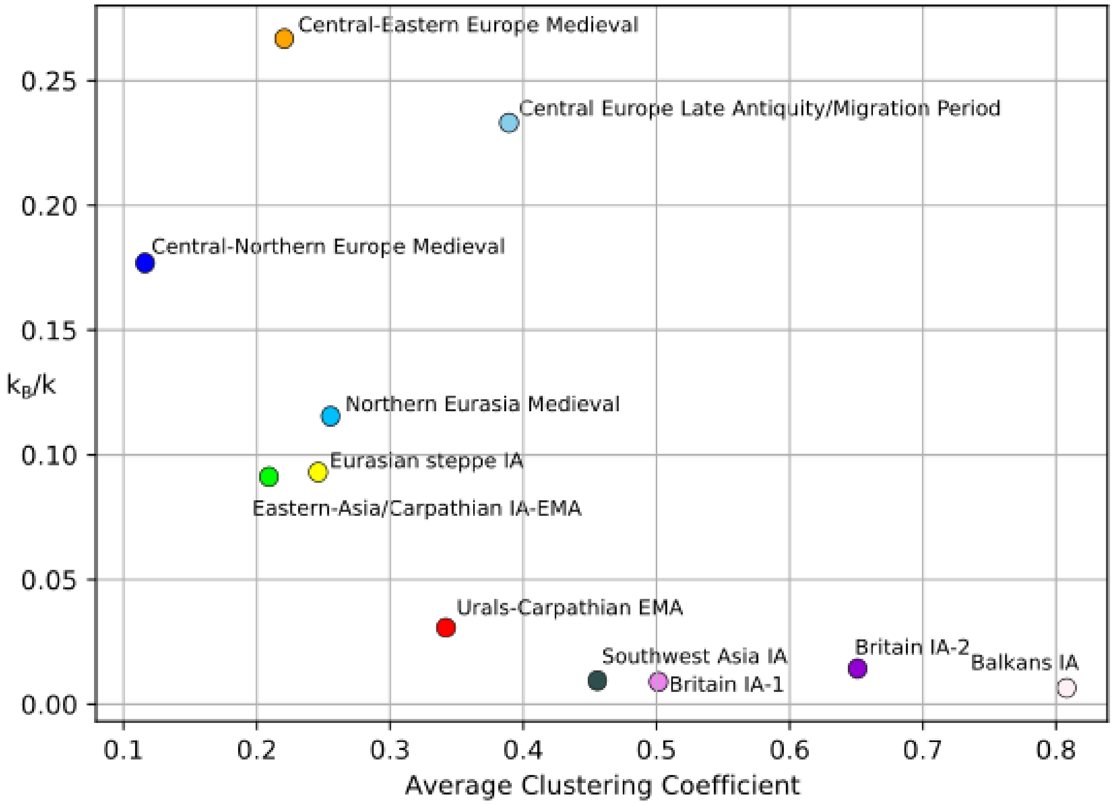

**Figure S5:** A scatterplot displaying the average clustering coefficient of each cluster versus the ratio of within-module connections ($k_B$) to the total degree centrality (k) of the cluster. Each cluster is identified using the Leiden algorithm, which determines community structure in the network. The 'modules' in this context refer to the clusters defined by this algorithm.

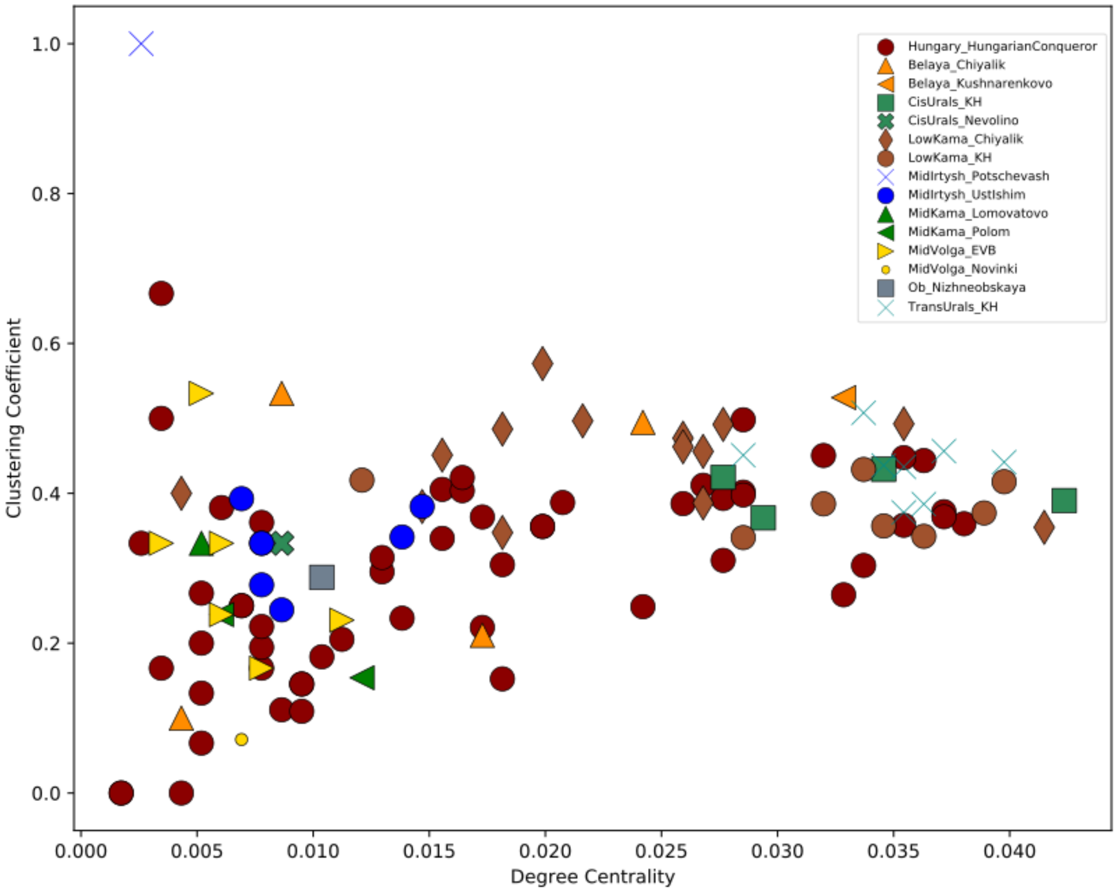

**Figure S6:** Scatterplot of degree centrality (k) versus clustering coefficient for all individuals in the Urals-Carpathian EMA cluster.

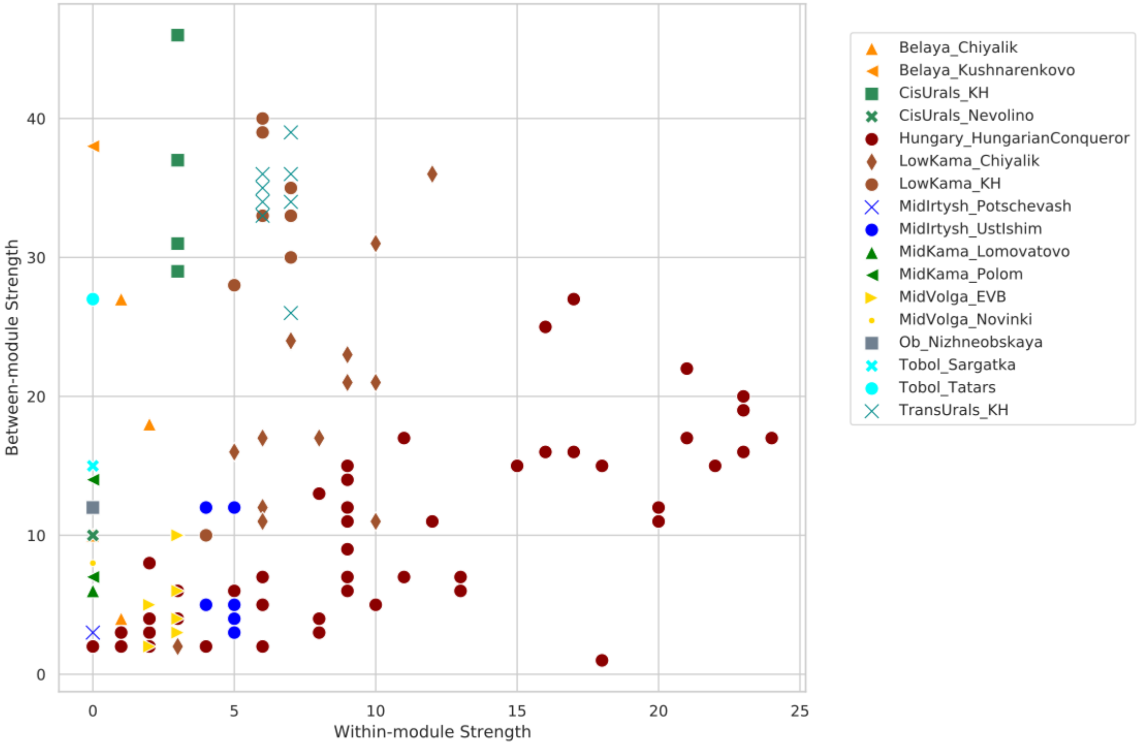

**Figure S7:** Scatterplot of Between-Module Strength versus Within-Module Strength in the *Urals-Carpathian EMA* Cluster (modules defined by groups)

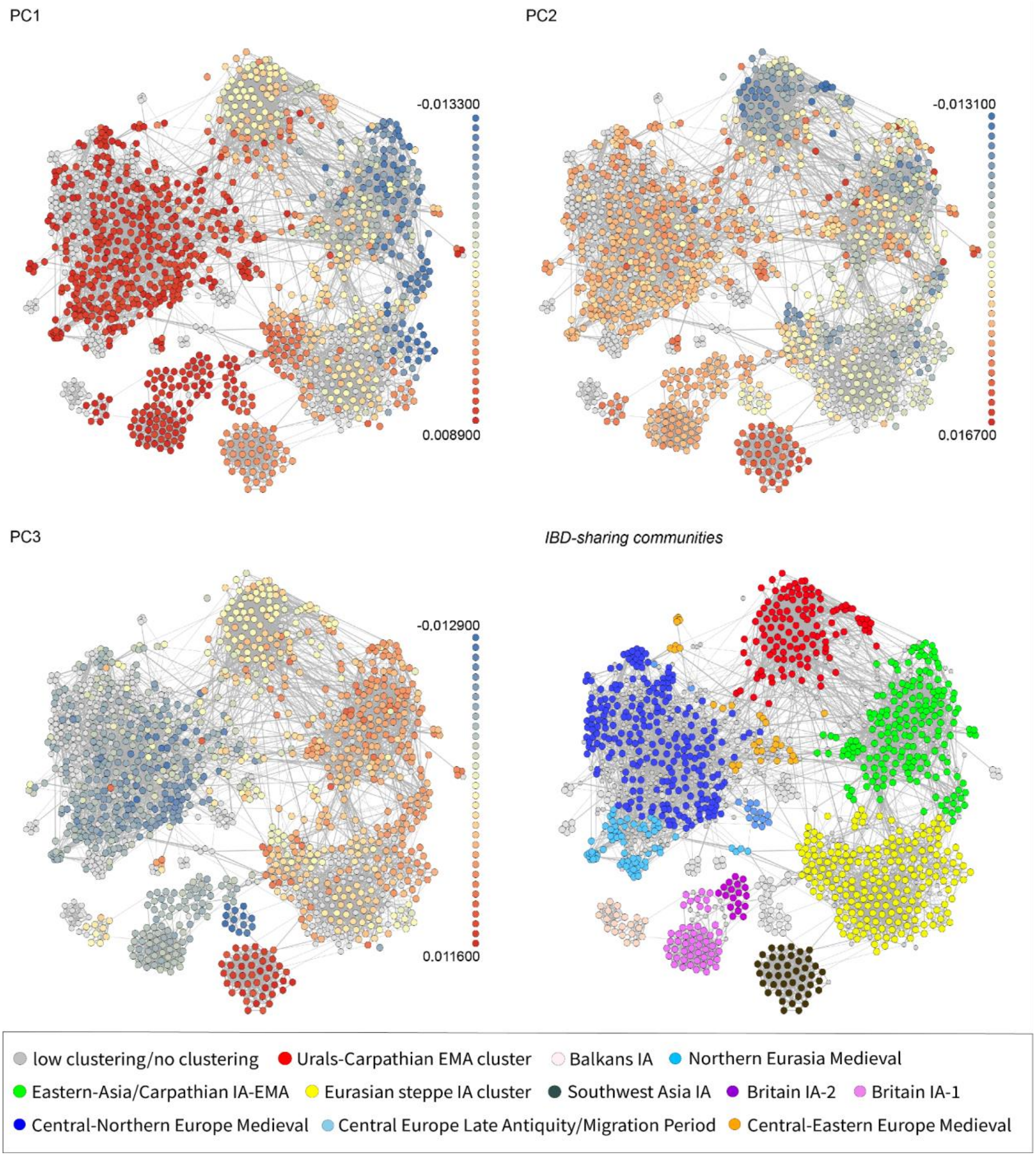

**Figure S8:** Individuals of the IBD-sharing network presented in PC1/PC2/PC3 spaces, projected on modern Eurasian individuals(*50*).

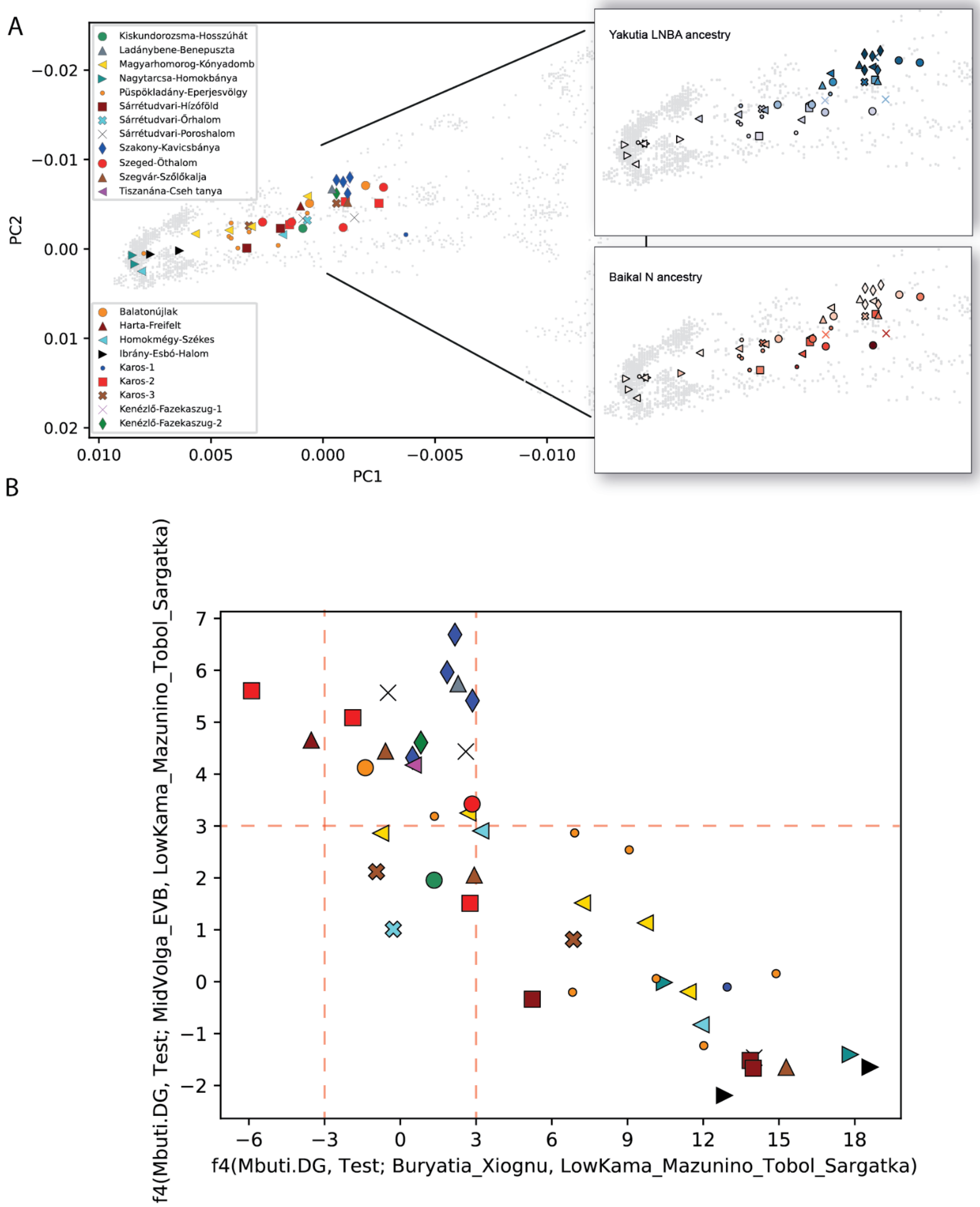

**Figure S9:** Dual ancestry of the 10th century Carpathian Basin individuals (EMMs) within the *Urals-Carpathian EMA* IBD-sharing cluster. **A:** PCA projection with estimates of the Yakutia LNBA and Baikal Neolithic ancestry components from the supervised *ADMIXTURE* analysis presented in Fig. 2B are shown on the right. In the case of the Yakutia LNBA ancestry, deeper *blue* color means higher proportion of this ancestry, while deeper *red* color means higher proportion of the Baikal Neolithic ancestry. **B:** 2D scatterplot illustrating *f4*-statistics. The test was carried out for combined Uralic Migration Period ancestries (Low-Kama Mazunino, Tobol Sargatka) in order to test allele-sharing differences from this joint Volga-Uralic component.

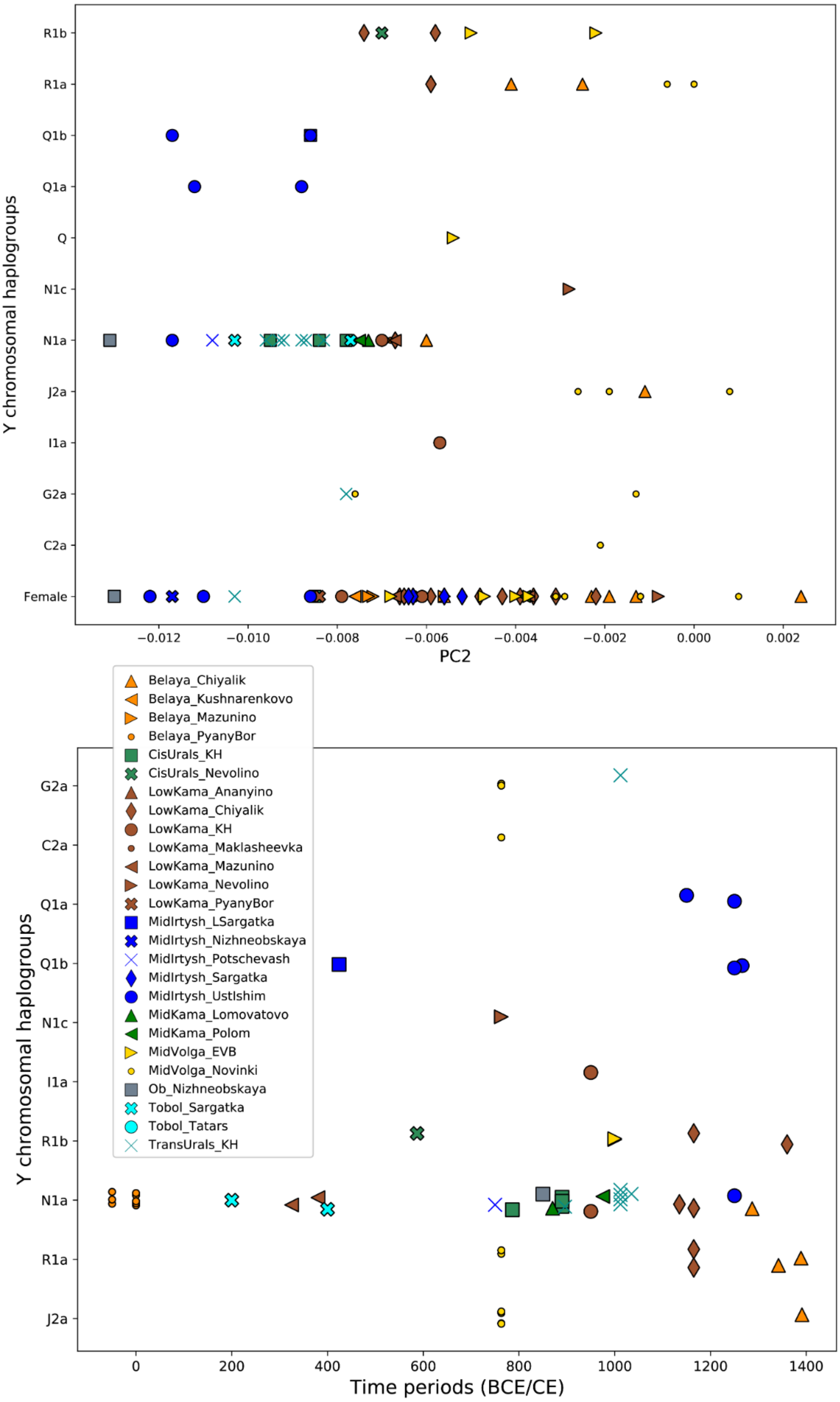

**Figure S10:** Scatterplots showing distribution of Y-chromosomal haplogroups of Volga-Uralic individuals, along PC2 and in different time periods.

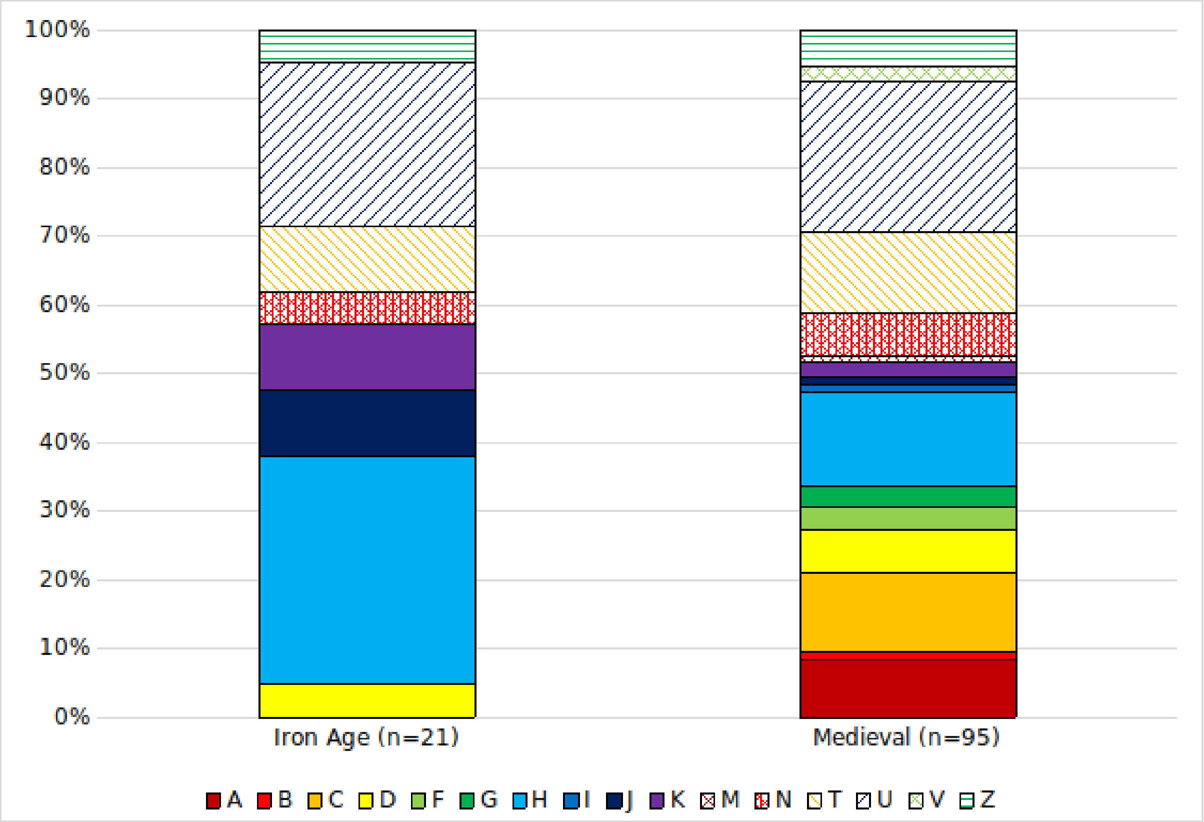

**Figure S11:** Frequencies of mtDNA haplogroups across different time periods, from all circum-Uralic regions under study.

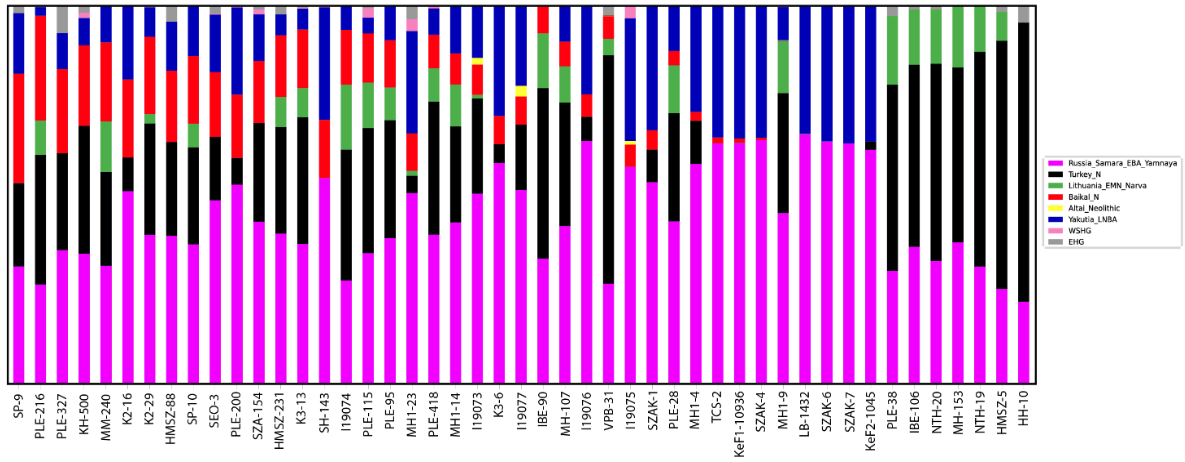

**Figure S12:** A supervised *ADMIXTURE* (K=8) analysis of 10th century Carpathian Basin indviduals from the *Ural-Carpathian EMA* IBD-sharing community.

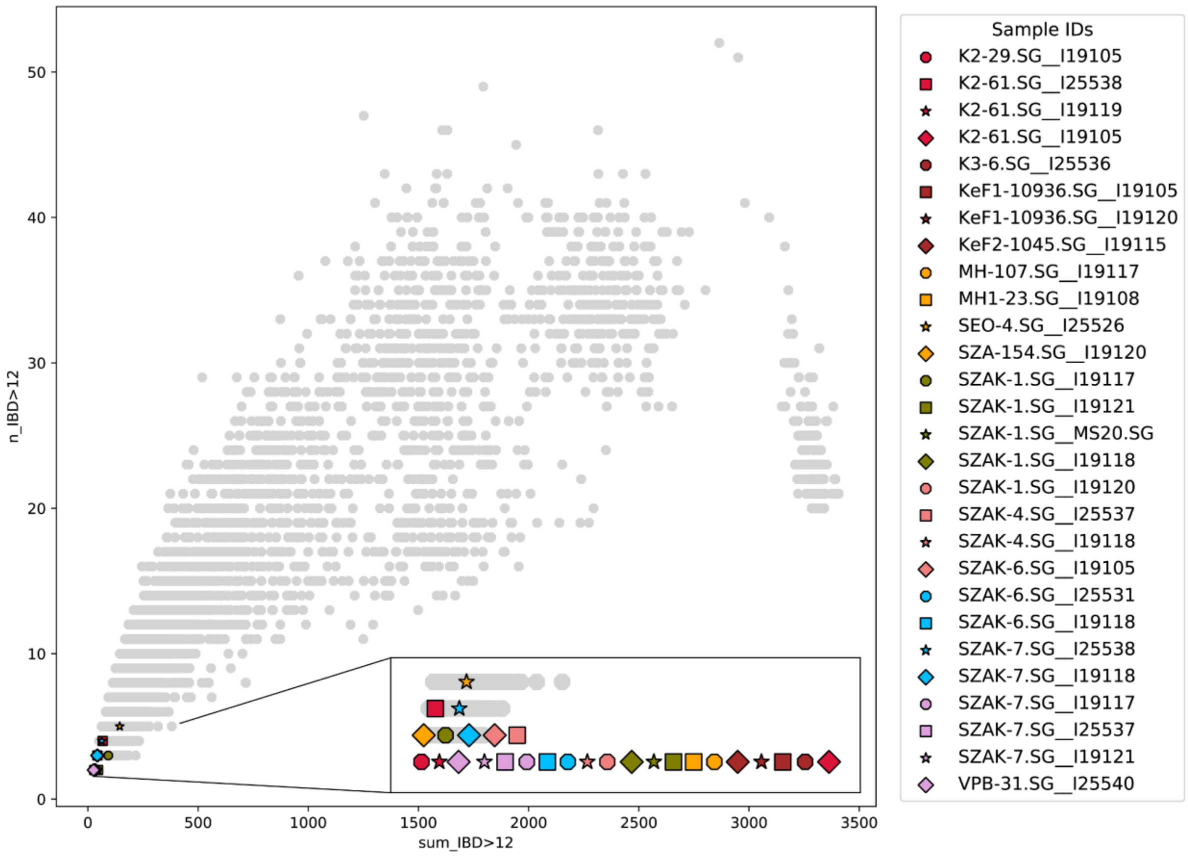

**Figure S13:** Distribution of IBD segments in individuals from the Volga-Ural region and Early Medieval Carpathian Basin (Table S1). Only pairs of individuals sharing at least two segments longer than 12 cM were considered.

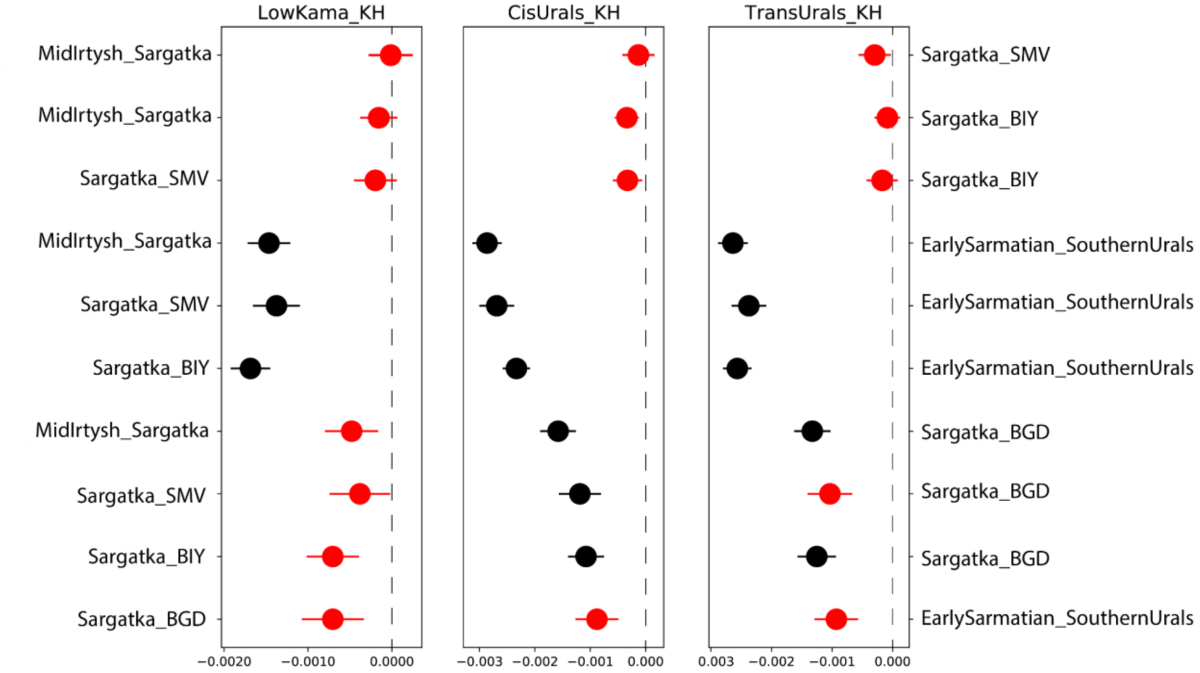

**FIgure S14:** *f4*-statistics *(*Mbuti, KH_group; EIA_test_1, EIA_test_2*)* assess allele-sharing among KH groups and various Early Iron Age references. This analysis includes individuals from the Sargatka horizon at the Shmakovo (SMV), Bogdanovka (BGD), and Mountain Bitiya (BIY) sites as described by Gnecchi-Ruscone et al. (2021). Additionally, it features groups from the Southern Uralic and Sarmatian cultures as reported by Jarve et al. (2019), alongside our newly introduced early Iron Age groups. Red markers denote |Z-score| < 3

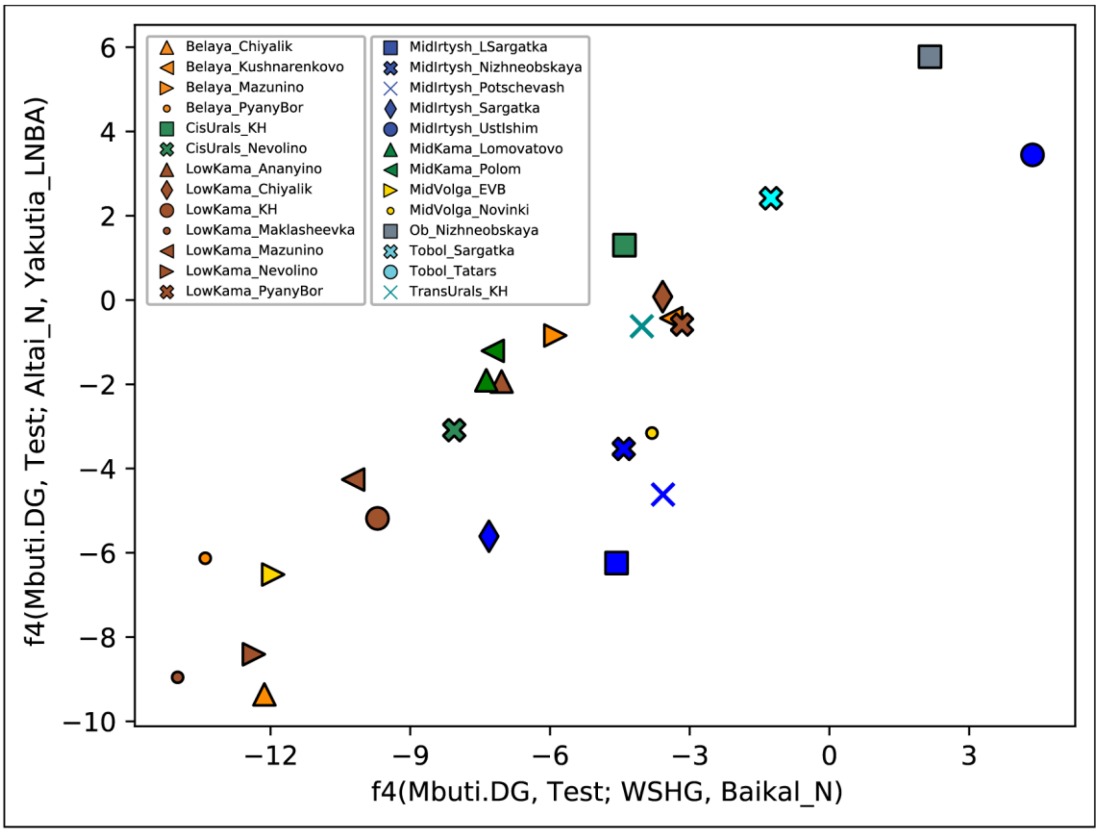

**Figure S15:** A scatterplot illustrating *f4*-statistics (Mbuti, Test; WSHG, Baikal Neolithic) and (Mbuti, Test; Altai Neolithic, Yakutia LNBA).

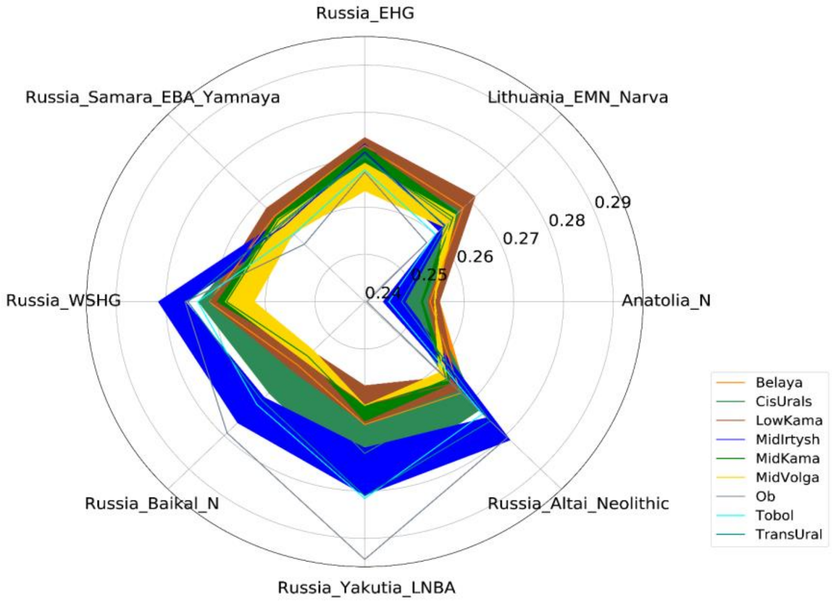

**Figure S16:** Outgroup *f3*-statistics (Mbuti; Test group, Reference group), comparing reference populations with newly sequenced ones (detailed in Supplementary Table 3) from the Volga-Ural region. Samples are grouped together by ecoregions. On this polarity map, the wider line means higher standard deviation for the f3 values. Groups from the east of the Ural Mountains generally share more alleles with WSHG, Altai Neolithic, Baikal Neolithic, and Yakutia LNBA. An exception is the Mid-Irtysh Niznheobskaya individual, which aligns with Low-Kama groups. Specifically, Mid-Irtysh Sargatka and Ob Nizhneobskaya exhibit affinity to Yakutia LNBA and Baikal Neolithic; the Ust-Ishim and Potchevash groups are closer to WSHG. There is low allele sharing with the Anatolian Neolithic in correlation with geographic proximity to this reference.

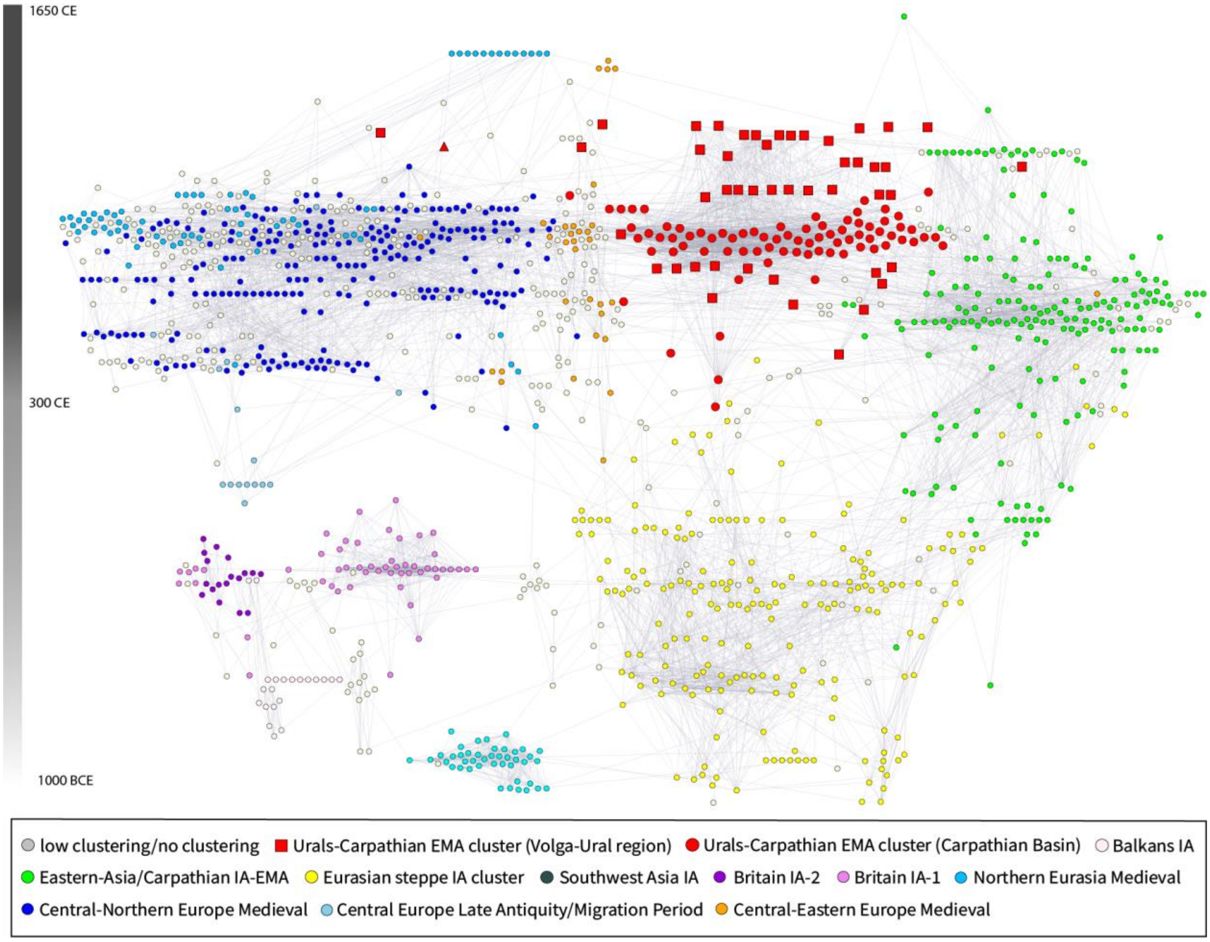

**Figure S17:** Time-ordered IBD-sharing network version of Figure 3.

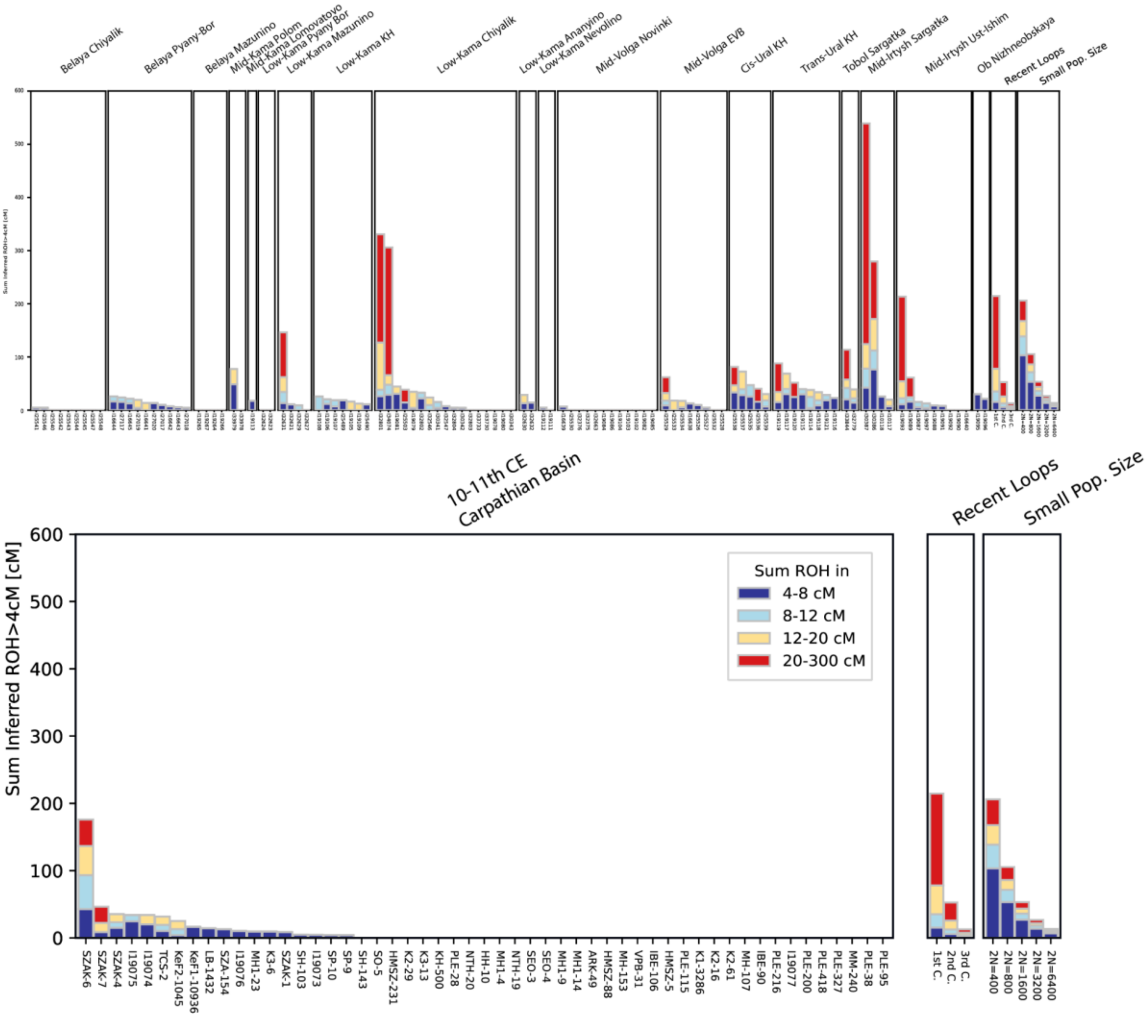

**Figure S18:** *hapROH* results for the Volga-Uralian and 10-11th CE Carpathian Basin invividuals(both newly presented and previously published in *Maróti et al. 2022*. )

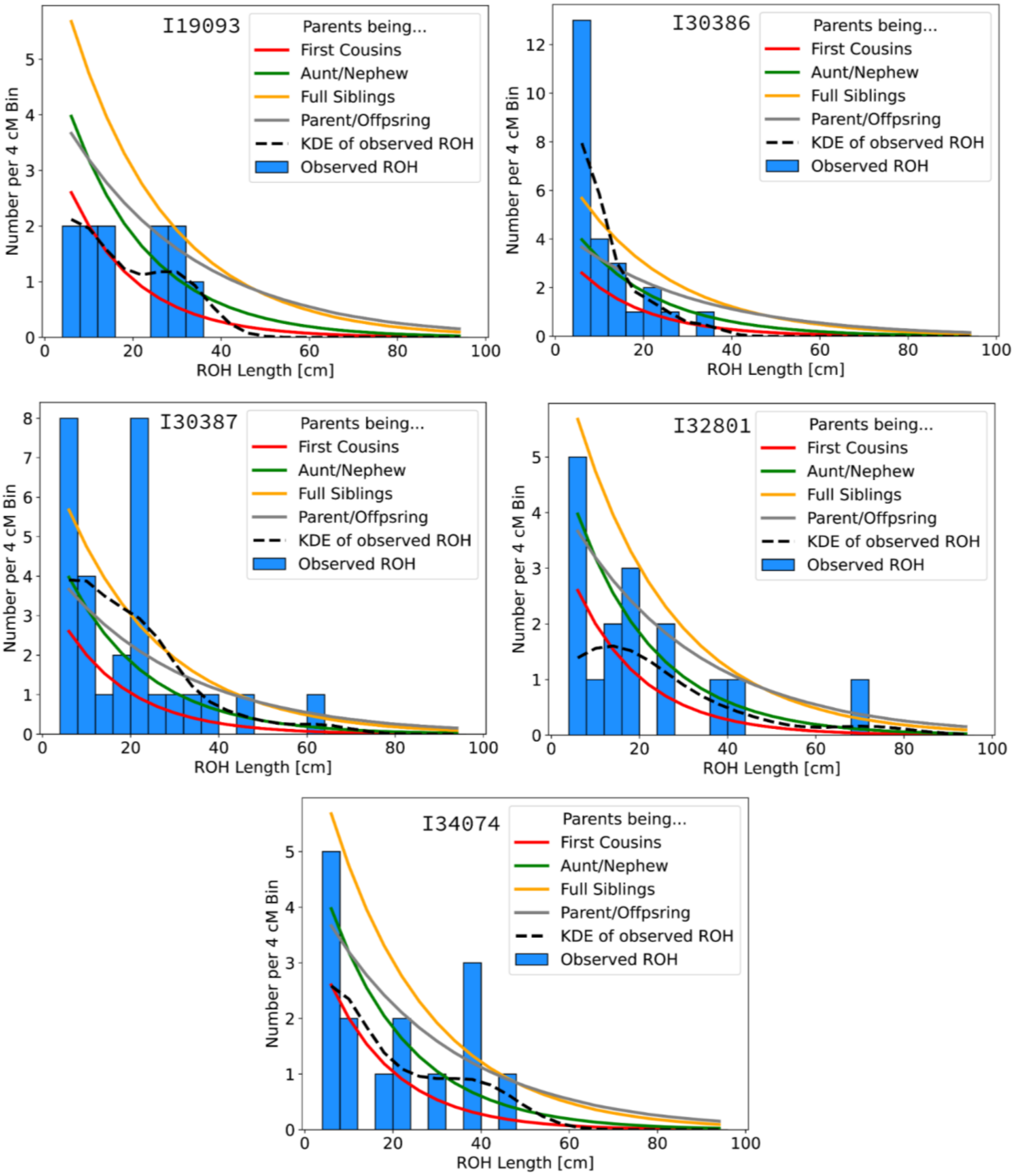

**Figure S19:** Individual ROH histograms of individuals that show high (>50cM SUM IBD from >20cM segment) signal of parental relatedness. They show marriages of at least first cousin level relatives.

| ***GroupID1*** | ***Ind1*** | ***Date1*** | **Sex** | **Y/mtDNA** | ***GroupID2*** | ***Ind2*** | **Sex** | **Y/mtDNA** | ***Date2*** | ***total length of shared IBD segments >2 x 12 cM*** |
|---|---|---|---|---|---|---|---|---|---|---|
| 10-11th century CB (EMM) | SEO-4 | 900-1000 CE | male | G2a/T2g1a | Mid-Volga EVB | I25526 | male | Q/B5b4 | 850-1050 CE | 144 |
| 10-11th century CB (EMM) | SZAK-1 | 900-1000 CE | male | N1a/T2d1b1 | Trans-Urals KH | I19117 | male | N1a/N1a | ***771-937 calCE*** | 92 |
| 10-11th century CB (EMM) | K2-61 | 900-950 CE | male | R1a/U4d2 | Cis-Urals KH | I25538 | male | N1a/U5a1g1 | 664-1016 CE | 67 |
| 10-11th century CB (EMM) | SZAK-7 | 900-1000 CE | female | -/D5a1 | Trans-Urals KH | I19118 | male | G2a/A+152 | 772-1152 CE | 42 |
| 10-11th century CB (EMM) | SZAK-7 | 900-1000 CE | female | -/D5a1 | Cis-Urals KH | I25538 | male | N1a/U5a1g1 | 664-1016 CE | 63 |
| 10-11th century CB (EMM) | SZAK-4 | 900-1000 CE | female | -/HV4a2a | Cis-Urals KH | I25537 | male | N1a/H6a1b | 664-1016 CE | 43 |
| 10-11th century CB (EMM) | SZA-154 | 900-1000 CE | female | -/B5b4 | Trans-Urals KH | I19120 | male | N1a/A12a | 772-1152 CE | 42 |
| 10-11th century CB (EMM) | SZAK-6 | 900-1000 CE | female | -/A16 | Low-Kama KH | I19105 | female | -/A12a | 850-950 CE | 45 |
| 10-11th century CB (EMM) | SZAK-1 | 900-1000 CE | male | N1a/T2d1b1 | Trans-Urals KH | I19121 | male | N1a/U5a1a1 | ***879-1150 calCE*** | 46 |
| 10-11th century CB (EMM) | K3-6 | 900-1000 CE | female | -/B4d1 | Cis-Urals KH | I25536 | male | N1a/C4a2 | ***664-827 calCE*** | 46 |
| 10-11th century CB (EMM) | SZAK-1 | 900-1000 CE | male | N1a/T2d1b1 | Trans-Urals KH | MS20(Gerber et al.) | male | N1a/D4j | 772-1152 CE | 45 |
| 10-11th century CB (EMM) | K2-61 | 900-950 CE | male | R1a/U4d2 | Trans-Urals KH | I19119 | male | N1a/C4a1 | 772-1152 CE | 39 |
| 10-11th century CB (EMM) | SZAK-4 | 900-1000 CE | female | -/HV4a2a | Trans-Urals KH | I19118 | male | G2a/A+152 | 772-1152 CE | 37 |
| 10-11th century CB (EMM) | KeF2-1045 | 900-1000 CE | male | N1a/N1a | Trans-Urals KH | I19115 | male | N1a/H40b | 772-1152 CE | 32 |
| 10-11th century CB (EMM) | SZAK-6 | 900-1000 CE | female | -/A16 | Trans-Urals KH | I19118 | male | G2a/A+152 | 772-1152 CE | 31 |
| 10-11th century CB (EMM) | SZAK-1 | 900-1000 CE | male | N1a/T2d1b1 | Trans-Urals KH | I19118 | male | G2a/A+152 | 772-1152 CE | 29 |

| | | | | | | | | | | |
|---|---|---|---|---|---|---|---|---|---|---|
| 10-11th century CB (EMM) | SZAK-7 | 900-1000 CE | female | -/D5a1 | Cis-Urals KH | I25537 | male | N1a/H6a1b | 664-1016 CE | 29 |
| 10-11th century CB (EMM) | KeF1-10936 | 900-1000 CE | male | Q1a/C4b | Low-Kama KH | I19105 | female | -/A12a | 850-950 CE | 29 |
| 10-11th century CB (EMM) | KeF1-10936 | 900-1000 CE | male | Q1a/C4b | Trans-Urals KH | I19120 | male | N1a/A12a | 772-1152 CE | 27 |
| 10-11th century CB (EMM) | SZAK-7 | 900-1000 CE | female | -/D5a1 | Trans-Urals KH | I19117 | male | N1a/N1a | ***771-937 calCE*** | 38 |
| 10-11th century CB (EMM) | SZAK-6 | 900-1000 CE | female | -/A16 | Belaya Kushnarenkovo | I25531 | male | G2a/U2e1 | ***340-540 calCE*** | 37 |
| 10-11th century CB (EMM) | K2-61 | 900-950 CE | male | R1a/U4d2 | Low-Kama KH | I19105 | female | -/A12a | 850-950 CE | 35 |
| 10-11th century CB (EMM) | K2-29 | 900-1000 CE | male | N1a/J1b | Low-Kama KH | I19105 | female | -/A12a | 850-950 CE | 33 |
| 10-11th century CB (EMM) | MH1-23 | 900-1000 CE | male | D1a/N1a | Low-Kama KH | I19108 | male | I1a/T1a1 | 850-950 CE | 32 |
| 10-11th century CB (EMM) | MH-107 | 1000-1100 CE | female | -/H11a7 | Trans-Urals KH | I19117 | male | N1a/N1a | ***771-937 calCE*** | 29 |
| 10-11th century CB (EMM) | SZAK-7 | 900-1000 CE | female | -/D5a1 | Trans-Urals KH | I19121 | male | N1a/U5a1a1 | ***879-1150 calCE*** | 29 |
| 10-11th century CB (EMM) | SZAK-1 | 900-1000 CE | male | N1a/T2d1b1 | Trans-Urals KH | I19120 | male | N1a/A12a | 772-1152 CE | 26 |
| 10-11th century CB (EMM) | VPB-31 | 700-800 CE | female | -/HV10 | Belaya Chiyalik | I25540 | male | J2a/N1a | ***1281-1395 calCE*** | 26 |

**Table S1:** Extended table of IBD connections between 10-11th century Carpathian Basin and Volga-Ural Early Medieval individuals with at least two IBD segments longer than 12 cM.

| *10-11th century CB (EMM)* | *left population1* | *left population2* | *p-value* |
|---|---|---|---|
| ARK-49 | Cis-Urals KH | ‘Eu cline’ | 0.309141745446696 |
| IBE-90 | Cis-Urals KH | ‘Eu cline’ | 0.992447693535791 |
| VPB-31 | Cis-Urals KH | ‘Eu cline’ | 0.798162013182211 |
| K3-6 | Cis-Urals KH | x | 0.472779170208264 |
| LB-1432 | Cis-Urals KH | x | 0.222102840227951 |
| SZA-154 | Cis-Urals KH | x | 0.435764838950309 |
| SZAK-1 | Cis-Urals KH | x | 0.818811036570204 |
| SZAK-4 | Cis-Urals KH | x | 0.645515885839159 |
| SZAK-6 | Cis-Urals KH | x | 0.584706439353114 |
| HMSZ-231 | Low-Kama KH | x | 0.0893143301884131 |
| HMSZ-88 | Low-Kama KH | x | 0.30933688991315 |
| K2-16 | Low-Kama KH | x | 0.496642082875645 |
| PLE-216 | Low-Kama KH | x | 0.138722225078596 |
| SEO-3 | Low-Kama KH | x | 0.0778270880465192 |
| KeF2-1045 | Trans-Urals KH | x | 0.748075446806715 |
| SZAK-7 | Trans-Urals KH | x | 0.780828377183826 |
| MH1-9 | Trans-Urals KH | ‘Eu cline’ | 0.0866722081786074 |

**Table S2:** Cladality test of the 10-11th century Carpathian Basin individuals with a feasible model (p-value>0.05) from the *Urals-Carpathian EMA* cluster.

| | $N_e$ estimate | std.error |
|---|---|---|
| Belaya Pyany Bor | 2,513 | 551 |
| Trans-Urals KH | 1,022 | 161 |
| MidIrtysh Ust-Ishim | 2,483 | 605 |
| Low Kama Chiyalik | 1,831 | 384 |
| Low Kama KH | 1,576 | 355 |
| MidVolga EVB | 3,252 | 905 |

**Table S3:** Effective population sizes ($N_e$) for groups with at least 8 individuals estimated with *hapROH Ne vignette, considering 4-20 cM segments*.
